# Sphingomyelin−Cholesterol Synergy as a Lipid Chaperone Driving Cytolysin A Pore Formation

**DOI:** 10.64898/2026.01.27.702101

**Authors:** Achinta Sannigrahi, Debayani Chakraborty, Rimjhim Moral, Vishwesh Haricharan Rai, Avijeet Kulshrestha, Dibyajyoti Maity, Muhsin Vannan Challil, Ganapathy Ayappa, Sandip Paul, Rahul Roy

**Affiliations:** Department of Chemical Engineering, Indian Institute of Science, CV Raman Road, Bengaluru, Karnataka 560012, India; Department of Chemistry, Indian Institute of Technology Guwahati, Guwahati, Assam 781039, India; Department of Bioengineering, Indian Institute of Science, CV Raman Road, Bengaluru, Karnataka 560012, India; Department of Nature Sciences, City University of New York, New York, NY 10017; Current address: University of Texas Southwestern Medical Center, 5323 Harry Hines Blvd, Dallas, Texas 75390, USA

**Keywords:** pore-forming toxin, sphingomyelin, cholesterol, lipid membrane, protein folding, molecular chaperone, host-pathogen interaction

## Abstract

Bacterial pathogens rely on pore-forming toxins (PFTs) to breach host barriers, a feat that requires water-soluble monomers to spontaneously metamorphose into membrane-inserted pores. In the extracellular milieu, these toxins must navigate complex folding landscapes without ATP-dependent chaperones. Here, we show that the synergistic interplay of host sphingomyelin (SM) and cholesterol (CHOL) acts as an intrinsic “lipid chaperone” that drives the conformational maturation of the bacterial toxin Cytolysin A (ClyA). Using leakage assays on vesicles and suspended lipid bilayer (SuLBs) arrays, atomistic molecular dynamics (MD) simulations, cell membrane permeabilization assays and biophysical measurements, we demonstrate that SM-CHOL synergy accelerates the rate-limiting unfolding of the toxin’s membrane bound β-tongue motif into a reactive “molten globule” intermediate and guides its subsequent refolding into functional α-helical conformation crucial for pore formation. We identify conserved lysine residues (K175 and K206) as critical molecular sensors that detect this specific lipid signature to drive productive transformation. This assembly process reciprocally remodels the host membrane and dismantling liquid-ordered domains, suggesting a mechanical coupling between toxin folding and the disruption of host lipid homeostasis. Our findings establish a paradigm of lipid-mediated chaperoning, revealing how pathogens co-evolve to exploit host lipid complexity to overcome the energetic barriers of membrane insertion.

## Introduction

The evolutionary arms race between bacterial pathogens and their mammalian hosts is frequently fought at the plasma membrane interface (Casadevall & Pirofski, 1999; Kostow & Welch, 2023). To breach this barrier, pathogens have evolved a diverse arsenal of pore-forming toxins (PFTs), virulence factors capable of compromising cellular integrity to facilitate invasion (Vadia *et al*, 2011). These toxins present a unique biophysical paradox: they are secreted as stable, water-soluble monomers but must spontaneously undergo a radical structural metamorphosis to assemble into thermodynamically stable, hydrophobic pores within the target membrane. Inside the bacterial cytoplasm, such profound conformational transitions would typically be governed by ATP-dependent protein chaperones (Hartl & Hayer-Hartl, 2002). However, in the extracellular milieu, toxins must navigate this complex folding landscape autonomously. This raises a fundamental question: how do PFTs autonomously negotiate the formidable energetic landscape of membrane insertion without catalytic chaperones?

The mammalian plasma membrane is distinguished by the synergistic interplay between sphingomyelin (SM) and cholesterol (CHOL) (Das *et al*, 2014; Lingwood & Simons, 2010). The distinct biochemical membrane signature is precisely conserved through tightly coupled metabolic feedback loops, where the host synchronizes SM and CHOL levels via the ER-resident sterol sensor Scap and SREBP transcription factors (Gatt & Bierman, 1980; Scheek *et al*, 1997). Such rigorous regulation ensures that SM and CHOL are not merely coincident but exist in a functionally coupled composition where SM depletion actively suppresses cholesterol biosynthesis (Slotte & Bierman, 1988). While this strict conservation effectively compartmentalizes cellular signaling, it paradoxically generates discrete biophysical environments—characterized by unique interfacial tension and molecular packing— that serve as high-energy templates for membrane-active agents (Róg & Pasenkiewicz-Gierula, 2006; Smaby *et al*, 1994). Consequently, the metabolic conservation of this lipid architecture establishes an evolutionarily invariant target for pathogens, which exploit the host’s coupled SM-CHOL composition to facilitate precise membrane targeting. This co-evolutionary strategy is exemplified by the exquisite specificity of diverse virulence factors that navigate this landscape. For instance, viruses such as Influenza hijack cholesterol-rich domains to facilitate cellular entry (Li *et al*, 2022). Similarly, specialized toxins including Lysenin, Equinatoxin-II, and Ostreolysin A have evolved to distinguish dispersed SM, SM clusters, or SM–CHOL complexes (Endapally *et al*, 2019; Ishitsuka *et al*, 2004; Makino *et al*, 2015). However, while such studies demonstrate that lipid synergy dictates where pathogenic proteins target and bind, the mechanistic role of this concerted lipid action in actively catalyzing how these proteins fold remains largely unexplored.

Cytolysin A (ClyA), an *α*-PFT secreted by *Escherichia coli* and *Salmonella pathovars*, serves as an exemplary model to investigate how specific membrane environments facilitate complex protein maturation. ClyA assembly involves one of the most dramatic conformational rearrangements known in toxin biology: a transition from a monomeric β-tongue motif to a membrane-inserted helix-turn-helix and associated translocation of the N-terminal *α*-helix to insert and puncture the membrane during oligomeric pore formation (Mueller *et al*, 2009). This profound structural transition is characterized by a dramatic and indispensable dependence on host cholesterol (Sathyanarayana *et al*, 2018). Prior biophysical investigations and molecular dynamics (MD) simulations have established that cholesterol acts not only to stabilize oligomeric pore intermediates but also to actively catalyze the local unfolding of the hydrophobic β-tongue motif (Sathyanarayana *et al*., 2018). Moreover, intrinsic conformational flexibility has been identified as a governing determinant of lytic competency, distinguishing functionally plastic toxins from rigid, inactive mutants (Kulshrestha *et al*, 2023a). Despite these insights, a critical disparity remains: ClyA displays orders-of-magnitude greater lytic kinetics against native-cell plasma membranes compared to reductionist binary lipid models (Sathyanarayana *et al*., 2018), implying that efficient assembly requires multicomponent lipid cooperativity beyond the solitary action of sterols. Specifically, how the synergy between cholesterol and other host lipids, such as sphingomyelin, modulates the thermodynamic trajectory of ClyA membrane insertion and pore formation remains largely unresolved.

Here, we combine ensemble biophysical and cell lysis kinetics, cellular membrane imaging, high-throughput single-microwell suspended lipid bilayer measurements, and atomistic molecular dynamics simulations to demonstrate that SM and CHOL act in concert as a “lipid chaperone” for ClyA. We show that this lipid synergy dramatically stimulates ClyA’s lytic activity accelerating the unfolding of the ClyA monomer into a critical molten globule-like intermediate and guiding its refolding into a lytic-competent intermediate. We identify conserved lysine residues (K175 and K206) as the molecular sensors for this interaction, linking lipid recognition directly to conformational plasticity. Finally, we observe that this lipid dependent ClyA assembly process actively remodels the host membrane and perturbs the sterol homeostasis, dispersing lipid rafts and triggering lipid droplet biogenesis. Our findings not only demonstrate a mechanistic link between lipid composition, protein structure and toxin function, but also the broader significance of SM–CHOL synergy in regulating membrane protein dynamics, cellular homeostasis, and host defense. This paradigm suggests that co-evolution of host and pathogen has driven bacterial toxins to exploit unique host lipid synergies, refining molecular interactions at the membrane interface for precise cellular targeting.

## Results

### Sphingomyelin–Cholesterol Synergy Selectively Enhances ClyA Pore Formation

While cholesterol is essential for stabilizing ClyA intermediates, binary lipid models fail to replicate the rapid lytic kinetics observed in native membranes(Sathyanarayana *et al*., 2018). For instance, the profound ClyA activity loss following methyl-β-cyclodextrin (mβCD) treatment was at least an order larger than what cholesterol addition alone could enhance(Sathyanarayana *et al*., 2018). It is well-established that mβCD not only extracts cholesterol but also fundamentally disrupts the synergistic balance between sphingomyelin (SM) and cholesterol (CHOL)(Zidovetzki & Levitan, 2007). This led us to hypothesize that ClyA might specifically target this synergistic organization. Supporting this view, a sequence alignment of ClyA across diverse bacterial species revealed a high conservation of lysine and arginine residues (**Figure S1A**). This cationic enrichment is a hallmark of other known SM-binding proteins, suggesting that ClyA may have evolved a specific “cationic sensor” to detect the electrostatic and packing signatures of SM-rich domains (**Figure S1B**). As sphingomyelin and cholesterol are intractably coupled, with SM levels strictly regulating the distribution of cholesterol into sequestered and accessible pools (**Figure S2A**) (Das *et al*., 2014), the profound lytic deficits observed upon mβCD treatment suggest that SM could be responsible for modulating cholesterol mediated ClyA functional assembly. To test this, we exposed RAW 264.7 macrophages to wild-type ClyA and assessed pore formation using fluorescence microscopy with propidium iodide (PI) uptake as a marker of membrane permeabilization (**Figure 1A** and **Figure S2B**). RAW 264.7 cells exhibited a pronounced increase in PI mean fluorescence intensity, confirming efficient pore formation (**Figure 1A** and **Figure S2B**). Similar, ClyA dependent PI uptake was seen for Vero epithelial cells but we noted that macrophages undergo faster and more extensive membrane permeabilization compared to Vero cells (**Figure S2C**). This result aligns with recent findings that ClyA from *Salmonella Typhi* exhibits selective lysis of macrophages and erythrocytes while sparing epithelial cells(Krone *et al*, 2024). While thin-layer chromatography (TLC) verified similar total SM–CHOL content in both cell types (**Figure S2D**), imaging of GM1-lipid domains (enriched in SM and CHOL (Becucci *et al*, 2014; Vanderroost *et al*, 2023; Yuan *et al*, 2002)) with CtxB-Alexa647 revealed a substantially high microdomain richness in macrophages (**Figure S2E**). Therefore, we conjecture that the spatial organization of lipid domains, rather than the overall abundance, may underlie the heightened sensitivity of macrophages to ClyA.

**Figure 1.**
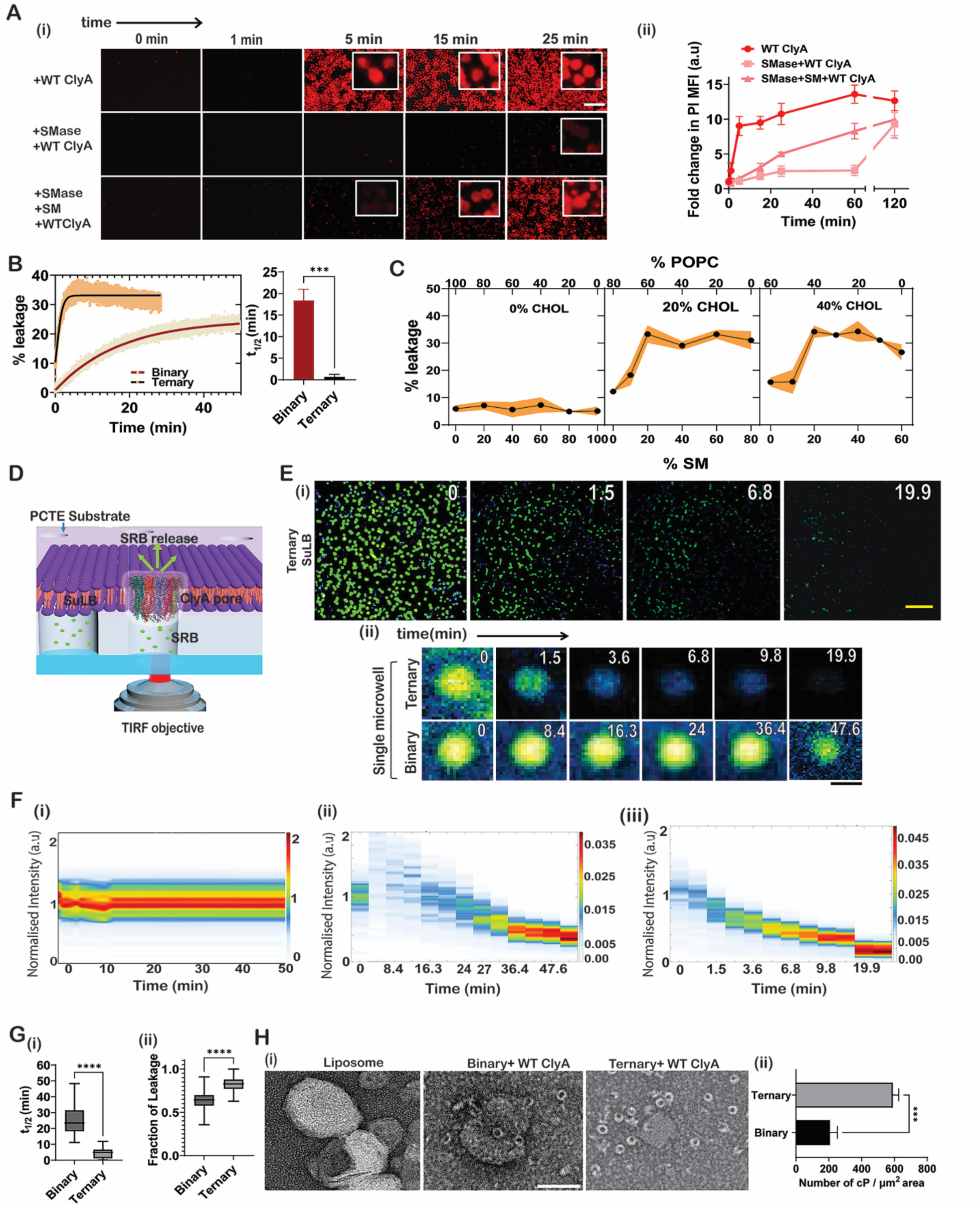
SM-CHOL Membranes Enhance the Lytic Activity of ClyA. (A) (i) Representative fluorescence micrographs of RAW 264.7 cells subjected to three treatment conditions: 100 nM WT ClyA alone; pre-treatment with sphingomyelinase (SMase, 50 mU/mL, 30 min, 37°C) followed by WT ClyA; sequential treatment with SMase, SM-liposomes (20 µM, 6 h), and WT ClyA. Membrane permeabilization was assessed via propidium iodide (PI) uptake. RAW 264.7 cells (2×10^4^/well) were seeded in 96-well plates, cultured in DMEM supplemented with 10% FBS and 1% penicillin-streptomycin (Day 0) and treated as indicated (Day 1). Following SMase exposure, cells were washed thrice with PBS before subsequent treatments. PI incorporation was monitored by fluorescence microscopy. Scale bar is 100 μm. (ii) Quantification of PI mean fluorescence intensity (MFI) over time in treated cells. Data represent mean ± SD (n = 3 independent experiments). (B) Kinetics of calcein release from binary (POPC-CHOL, 70:30) and ternary (POPC-CHOL-SM, equimolar) small unilamellar vesicles (SUVs) upon WT ClyA exposure(left). The right plot shows the comparison of the half-life of calcein leakage in each liposomal formulation. Data represent mean ± SD (n = 3). Statistical analysis was performed using unpaired t-tests, ***P=0.0003. (C). Percentage of calcein leakage from liposomes entrapping calcein, composed of varying mol% of SM and POPC, at three different CHOL concentrations (0%, 20%, and 40%). Liposomes (final concentration: 800 μM) of various compositions were incubated with WT ClyA (working concentration: 500 nM) for 5 minutes. The fluorescence intensity of dequenched calcein was measured, and leakage percentages were calculated relative to total leakage induced by 1% Triton X-100 treatment. Data are presented as mean ± standard deviation (SD), with *n* = 3 representing the number of experimental repeats. The shaded area in the graph indicates the standard deviation across the three independent experiments. (D) Assessment of ClyA pore formation in suspended lipid bilayer (SuLB) systems constructed on polycarbonate track-etched (PCTE) substrates. Microwells (1 µm in diameter) were filled with 100 nM sulforhodamine B (SRB) as a fluorescent reporter. Total internal reflection fluorescence (TIRF) microscopy was used to track SRB release following treatment with 100 nM monomeric WT ClyA. (E) (i) Time-lapse TIRF images of SRB release from ternary SuLB systems following WT ClyA addition. Scale bars, 10 µm. (ii) Single microwell imaging comparing binary and ternary SuLB systems post-WT ClyA exposure. Scale bar is 1μm. (F) Graph shows the temporal well density distributions: (i) ternary SuLB treated with NaP-buffer; (ii) binary and (iii) ternary SuLB following WT ClyA exposure. Color bar indicates relative probability density. (G) (i) Box plots depicting the half-life of SRB release from individual 1 µm cavities in binary and ternary SuLB systems following WT ClyA treatment. Data represent mean ± SD (n = 3). Statistical significance was assessed using unpaired t-tests, ****P =0.00035. (ii) Plot showing the extent of SRB leakage from binary and ternary SuLBs following WT ClyA treatment. Number of cavities analyzed: 515 (ternary) and 454 (binary). Data represent mean ± SD (n = 3). Statistical significance was assessed using unpaired t-tests, ****P <0.0001. (H) (i) Negative-stain transmission electron micrographs of untreated liposomes, binary liposomes + WT ClyA, and ternary liposomes + WT ClyA. Scale bar is 50 nm. (ii) Quantification of complete WT ClyA pore structures per µm² grid area in binary and ternary systems. Data are presented as mean ± SD (n = 3). Statistical significance: ***P = 0.0073.

To establish the direct role of SM in ClyA-mediated pore formation, we selectively depleted SM from RAW 264.7 cell membranes using sphingomyelinase (SMase) prior to toxin exposure. This enzymatic degradation resulted in a pronounced (∼12-fold) reduction in the rate and extent of ClyA-driven pore formation, as measured by PI uptake (**Figure 1A** and **Figure S2F**). Remarkably, supplementing SMase-treated cells with SM-containing liposomes significantly restored ClyA activity, confirming that SM is indispensable for efficient ClyA-induced pore formation (**Figure 1A**). Such dramatic deficits in WT ClyA activity in SM depleted RAW macrophages were consistent across a range of toxin concentrations (**Figure S2G**), suggesting that higher toxin concentration cannot compensate for sphingomyelin dependence. Similarly, SMase pretreatment in Vero cells significantly attenuated WT ClyA-induced PI uptake (**Figure S2H**), reinforcing that SM is a central rate-limiting determinant of ClyA lytic function in diverse cell types. Collectively, these findings establish that SM and the organization of SM-enriched microdomains is critical for the efficiency of ClyA pore formation on mammalian cell membranes.

To further dissect how SM modulates ClyA function, we measured ClyA mediated calcein leakage from binary POPC–CHOL and ternary POPC–SM–CHOL liposomes. Ternary membranes demonstrated an approximately 18-fold enhancement in WT ClyA-induced dye release kinetics relative to binary and PC-only controls (**Figure 1B** and **Figure S3A**), with maximal activity defined by the SM–CHOL ratio and plateauing around 1:1 composition (**Figure S3A)**. Lipid compositions rich in both SM and CHOL displayed increased leakage, while PC contributed minimally. Additionally, leakage across liposomes with varying POPC and SM ratios and fixed CHOL levels revealed that increasing SM content synergistically elevated ClyA activity only in the presence of sufficient CHOL (**Figure 1C**). This enhancement closely mirrored the SM–CHOL binding profile observed for OlyA (Endapally *et al*., 2019), reinforcing the role of SM-CHOL synergy in promoting ClyA activity. Temperature-dependent assays showed that liposomes abundant in both SM and CHOL losing ClyA activity at elevated temperatures, further signifying lipid complexation as a critical driver towards ClyA’s lytic activity (**Figure S3B**). Among all tested compositions, equimolar POPC-SM-CHOL (33.3:33.3:33.3 molar ratio) composition, which closely emulates the plasma membrane outer leaflet(Lorent *et al*, 2020; Verkleij *et al*, 1973), exhibited the strongest ClyA activity (**Figure S3A**) and was therefore designated as our ternary model membrane.

Although the ternary membrane substantially increases ClyA-induced membrane rupture (**Figure 1B**), the overall extent of leakage remains modest—about 35% in ternary versus ∼20% in binary liposomes—compared to near complete lysis observed for erythrocytes by WT ClyA(Sathyanarayana *et al*., 2018). This discrepancy suggests that traditional liposome-based model membranes may not adequately replicate plasma membrane properties. Common experimental systems—SUVs, GUVs, and SLBs—are limited by factors such as high curvature, high heterogeneity, and surface artifacts, which complicate the analysis of protein-membrane interactions and pore formation. To overcome these limitations, we adapted a robust, easily fabricated suspended lipid bilayer (SuLB) platform using commercially available PCTE membranes with 1 μm microwells developed recently(Sannigrahi *et al*, 2022). The SuLB platform enables high-throughput ClyA pore formation kinetic measurements in synthetically controlled lipid compositions (**Figure 1D, Supplementary Material SM1** and **Figure S4**). To determine the lipid requirements for ClyA-mediated membrane disruption, we monitored the release kinetics of sulforhodamine B (SRB) from SuLB-capped microwells (**Figure 1D,E** and **Figure S5**). WT ClyA induced robust SRB efflux from ternary SuLBs, compared to binary POPC-CHOL SuLBs (**Figure 1E,F**). The absence of SRB release in buffer-only controls confirmed that membrane permeabilization was strictly dependent on the presence of both the toxin and a permissive lipid environment. Kinetic analysis revealed membrane pore formation rates in ternary systems to be ∼5-fold higher and improved lysis (> 75%) of the pores compared to the binary lipid membranes, paralleling the trends observed in liposome and cell assays (**Figure 1G**).

To test the hypothesis that the ternary lipid composition facilitates the efficient assembly of ClyA into its closed dodecameric pore structure, we employed atomic force microscopy (AFM) and negative-stain TEM to directly visualize the oligomeric state of WT ClyA on model membranes. AFM imaging of ternary SuLBs revealed a significant presence of ClyA oligomers, which were significantly reduced on binary SuLBs (**Figure S6A,B**). The abundance of these oligomers on ternary SuLBs corresponded closely to the SRB leakage kinetics and the observed microwell depressions following WT ClyA exposure (**Figure S6C**), linking structural assembly to functional activity. Similarly, analysis by negative-stain TEM, after incubating binary and ternary liposomes with WT ClyA, confirmed that complete pore structures were readily observed on ternary membranes but markedly diminished on binary lipid membranes (**Figure 1H**). The density of complete pores per μm² grid area was roughly threefold lower in the binary membranes compared to the ternary system, reinforcing the idea that delayed or reduced oligomerization in binary membranes underlies their slower membrane rupture. Collectively, these structural observations provide a strong functional link between optimal ternary lipid composition and the efficient and rapid assembly of functional ClyA pores.

### Early Stage Unfolding and Refolding of ClyA β-tongue is Facilitated by SM-CHOL

To pinpoint the mechanistic basis of SM–CHOL synergy, we first determined if the ternary lipid environment preferentially stabilizes the final assembly. All-atom molecular dynamics (MD) simulations of the dodecameric pore revealed that the mature complex exhibits comparable structural persistence and global integrity when embedded in either binary (POPC:CHOL) or ternary (POPC:SM:CHOL) bilayers (**Figure S7**). Convergent RMSD and *R_g_* trajectories confirm that the pore maintains structural stability across both environments. This indistinguishable stability implies that SM–CHOL synergy does not act as a thermodynamic stabilizer of the final pore assembly, but rather functions as a modulator of the upstream binding and oligomerization landscape.

Given that the final assembly is energetically insensitive to sphingomyelin, we hypothesized that the synergistic enhancement must stem from the stabilization of the initial membrane-bound monomeric conformation. Indeed, sedimentation assays revealed that ternary multilamellar vesicles (MLVs) recruited WT ClyA more efficiently than binary controls (∼72% versus ∼49% bound) (**Figure 2A**). While this represents a moderate increase in total protein association, it corresponds to a ∼2.7-fold enhancement in the binding equilibrium constant (**Figure 2A**). This thermodynamic advantage suggests that SM–CHOL synergy extends beyond simple electrostatic recruitment, potentially creating a favorable energetic landscape that primes the toxin for downstream conformational progression upon membrane binding.

**Figure 2.**
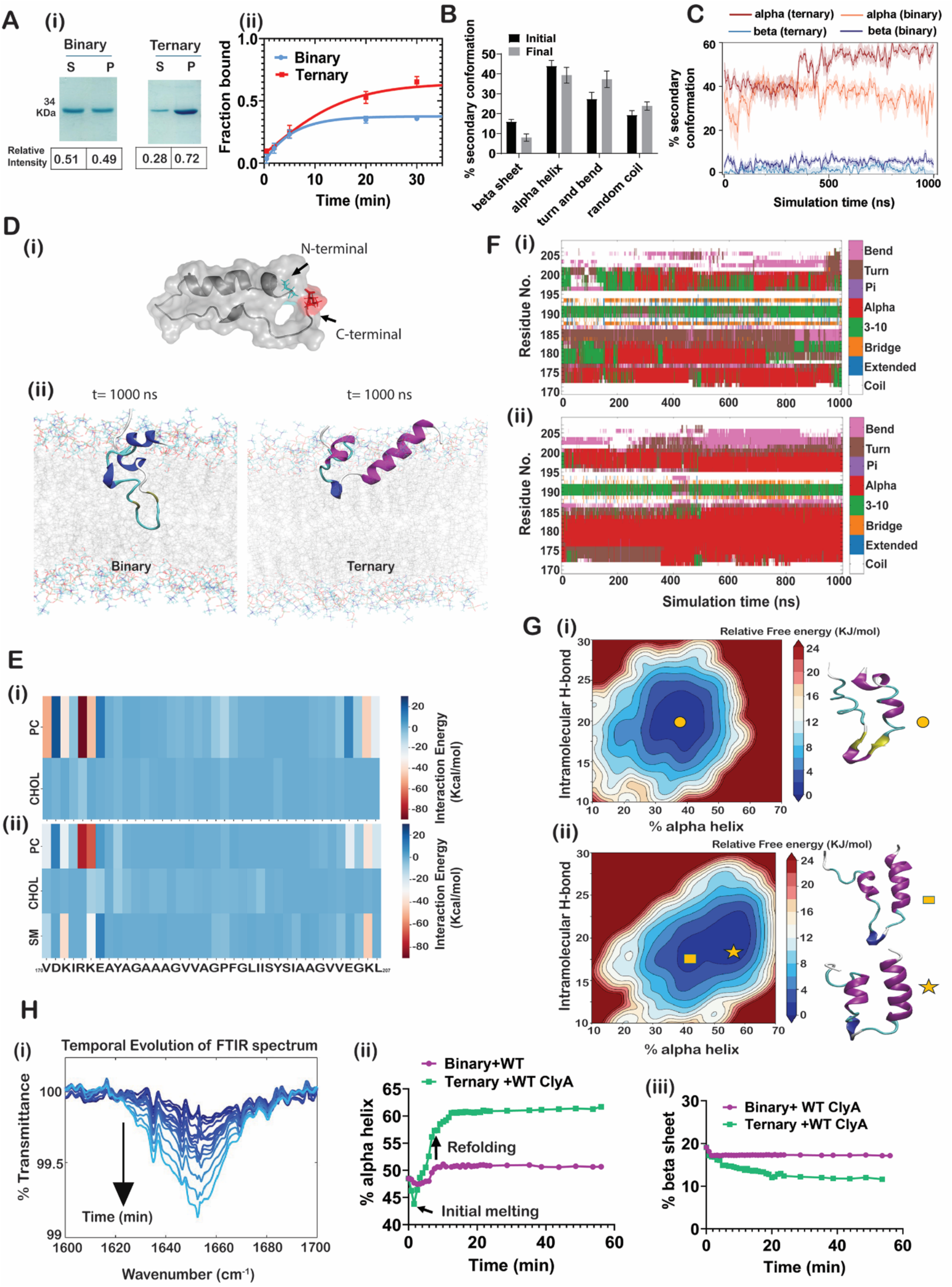
Ternary membranes enhance ClyA binding, induce early-stage unfolding, and facilitate refolding. (A) (i) Membrane binding of WT ClyA monomer was assessed using a sedimentation assay. SDS-PAGE shows WT ClyA in the supernatant (S) and pellet (P) fractions after incubation with binary and ternary multilamellar vesicles (MLVs) at 37°C for 30 minutes, followed by high-speed centrifugation at 23,000 × *g* for 1 hour. (ii) Time-dependent membrane binding of Cy5-labeled ClyA(Q56C) was analyzed with binary and ternary MLVs using sedimentation assays. Membrane-bound and unbound fractions were quantified at multiple time points. Data are presented as mean ± SD (n = 3, representing three independent experiments). (B) Molecular dynamics (MD) simulations show changes in the percentage of β-sheet content within the β-tongue domain of ClyA in binary vs. ternary membrane environments. The plot compares secondary structure content at the initial (0 ns) and final (300 ns) time points. Data are represented as mean ± SD (n = 3). (C) Secondary structure dynamics of the partially unfolded “molten globule”-like intermediate of the ClyA β-tongue were monitored via MD simulations. The shaded area in the graph indicates the standard deviation across the three replicates for each group. (D) (i) Structure of the unfolded molten globule (MG) intermediate of ClyA β-tongue. (ii). Final Snapshots of the molten globule-like β-tongue motif after simulations in binary and ternary membranes. (E) Heatmaps show residue-wise initial interactions of the molten globule-like ClyA β-tongue intermediate with membrane components: (i) PC and cholesterol (CHOL) in the binary system, and (ii) PC, sphingomyelin (SM), and CHOL in the ternary system. (F) Residue-level secondary structure changes of the molten globule-like intermediate during MD simulations in (i) binary and (ii) ternary membrane environments. A similar behavior of secondary structural change was observed in replicates. (G) Free energy profiles (in kJ/mol) illustrate the relationship between the number of intramolecular hydrogen bonds and the percentage of α-helical content in the molten globule intermediate, comparing (i) binary and (ii) ternary membrane systems. Snapshots highlight α-helix formation in each system. (H) Conformational transitions of WT ClyA in the presence of ternary membranes: (i) Time-resolved ATR-FTIR spectra (amide I region) track secondary structure evolution. (ii) Temporal changes in α-helical and (iii) β-sheet content were derived via deconvolution of ATR-FTIR data.

We reasoned that the β-tongue represents the primary energetic bottleneck for this membrane conformation stabilization process. As the initial point of membrane contact, it must physically engage the lipid bilayer while undergoing the most radical conformational metamorphosis in the assembly pathway (Giri Rao *et al*, 2016; Mueller *et al*., 2009). Several lines of evidence suggest that the β-hairpin to a helix-turn-helix conformation change of the β-tongue upon membrane insertion is essential for pore formation. The indispensability of this motif is underscored by the fact that restricting its conformational freedom—specifically through the rigidifying Y178F mutation or partial deletions— completely abolishes lytic activity (Fahie *et al*, 2018; Kulshrestha *et al*., 2023a). This is despite efficient membrane binding and complete ring assembly by these β-tongue mutants. On the other hand, mutations in the N-terminal helix display slower pore formation but complete lysis, whereas mutations in the extracellular domain have no effect on the lytic activity, emphasizing the predominant role of the β-tongue’s conformation on efficient pore formation (Kulshrestha *et al*., 2023a; Sathyanarayana *et al*., 2018).

To dissect the molecular choreography of this transition, we tracked the trajectory of the isolated β-tongue motif using atomistic simulations. While binary membranes induced only partial destabilization of the β-tongue architecture (characterized by decreased β-sheet and increased α-helical content as seen in **Figure S8A,B**), the ternary lipid environment triggered a profound conformational melting, characterized by a rapid and broad transition to random coil and turn elements (**Figure 2B** and **Figure S9A**). This the ternary environment triggered conformational melting, visually captured by the progressive loss of β-tongue architecture in residues V185–S197 (**Figure S10A**). Analysis of the interaction energy landscapes suggests a sequential “hand-off” mechanism driven by lipid–lipid competition. In binary membranes, cholesterol maintains persistent, high-affinity contacts with the hydrophobic core of the β-tongue (residues 172-193; **Figure S8C**), effectively acting as a denaturant to destabilize the native state as observed in longer MD simulations (Kulshrestha *et al*, 2023b). However, the introduction of sphingomyelin dramatically remodels this interaction profile with near complete loss of cholesterol interactions with the β-tongue core (**Figure S9B**). As shown in the simulation snapshots, sphingomyelin specifically recruits cationic sensors (Lys/Arg) while simultaneously forming complexes with cholesterol (**Figure S10A**). Our data supports a model where sphingomyelin competes for cholesterol, forming specific SM–CHOL complexes that effectively “frisk” or sequester sterols away from the protein interface. This distinct lipid choreography appears to drive a two-step activation process: cholesterol first lowers the energetic barrier for unfolding, but its subsequent sequestration by sphingomyelin clears the interface, allowing the molten intermediate to resolve into the lytic-competent α-helical state within the hydrophobic membrane bilayer. Experimental validation using tryptophan fluorescence to monitor urea-induced unfolding confirmed this model, revealing that the ternary lipid environment significantly accelerates the unfolding trajectory of WT ClyA relative to binary controls (**Figure S10B,C**).

Having established that the ternary environment promotes the initial unfolding of the β-tongue, we next asked whether it also orchestrates the subsequent structural refolding to the final β-tongue conformation in the membrane. To test this, we performed MD simulations starting not from the native structure, but from the partially unfolded β-tongue —a conformation serving as a proxy for the reactive “molten globule” (MG) intermediate previously identified in cholesterol-rich membranes (**Figure 2C** and **2D**; (Kulshrestha *et al*., 2023b)). This distinct starting configuration allowed us to isolate the refolding trajectory from the initial melting event. In binary (POPC–CHOL) membranes, this intermediate remained energetically frustrated. Electrostatic interactions favored cholesterol contacts with residues D171, R174, and K175, while van der Waals forces engaged the hydrophobic stretch Y178–203 (**Figure 2Ei**, **Figure S11**). Despite these interactions, residue-specific analysis revealed only partial helical gains in the N-terminal end (K172–K175) and transient 3_10_-helix formation in residues S197–A199 (**Figure 2Fi**, **Figure S12**). Effectively, the peptide failed to progress toward the folded state, remaining trapped in a shallow energy basin characterized by limited α-helical content (**Figure 2C** and **2D**). In stark contrast, the ternary environment acted as a conformational catalyst, driving significant refolding into an α-helical protomer-like conformation (**Figure 2C** and **2D**). Structural decomposition reveals that this refolding is driven by a cooperative “anchor-and-pack” mechanism. Sphingomyelin electrostatically tethers the N- and C-terminal “cationic sensors” (residues K172, K175, K206), stabilizing the boundaries of the folding domain (**Figure 2Eii, Figure S13**). Simultaneously, cholesterol engages the central hydrophobic core (residues 178–194) through strong van der Waals interactions, underscoring its critical role in facilitating refolding (**Figure 2Eii, Figure S13**). This bipartite lipid engagement allowed regions spanning D171–A181 and Y196–G201 to adopt stable α-helical conformations (**Figure 2Fii, Figure S14**), collectively forming an assembly-competent protomer structure. Free energy landscape analysis confirms this chaperone function: while binary membranes restricted the protein to a shallow, disordered basin (∼40% helicity), the ternary environment remodeled the landscape, revealing a second, deeper minimum at ∼60% helicity that drives the spontaneous transition to the folded state (**Figure 2G**).

Finally, we experimentally validated these simulation-derived dynamics by monitoring the conformational transitions of WT ClyA in real-time. Circular dichroism (CD) spectroscopy revealed that the addition of ternary liposomes triggered a swift increase in α-helical content without the transient structural loss observed in binary controls, possibly due to the accelerated folding pathway in ternary lipids (**Figure S15A**). This behavior stands in sharp contrast to the inactive mutant Y178F, which maintained a stable, high-helicity profile across both lipid environments (**Figure S15B**). Further characterization by ATR-FTIR spectroscopy confirmed that the Y178F mutation constitutively increases α-helical content while suppressing β-sheet elements (**Figure S15C**), indicative of a “locked” conformational switch that leads to functional loss akin to similar traps in other systems (Ha & Loh, 2012; Sannigrahi *et al*, 2018). For the wild-type toxin, time-resolved ATR-FTIR tracking of amide I vibrations (1600–1700 cm⁻¹) provided a granular view of the assembly process (Sannigrahi *et al*, 2021; Sannigrahi *et al*, 2024).

In binary membranes, liposome addition induced a rapid decline in both β-sheet and α-helical content, marking the formation of a partially unfolded intermediate (I) reminiscent of states seen in detergent-induced perturbations (Benke *et al*, 2015). Subsequent time-dependent recovery of α-helical content indicated structural maturation toward an assembly-competent protomer, culminating in formation of the complete pore (cP) (**Figure 2H, Figure S15D**). While similar intermediates formed in ternary liposomes, the kinetics of transition to intermediate and protomer states were markedly accelerated (**Figure 2H**) and α-helical content adopted in ternary liposome is dramatically higher that that observed in binary membrane supporting the notion that ternary lipid environments powerfully facilitate ClyA folding and functional assembly. Conversely, longitudinal ATR-FTIR of the Y178F mutant showed minimal secondary structure evolution (**Figure S15D**), reinforcing its inability to navigate this pathway. Collectively, these data demonstrate that the SM–CHOL environment facilitates ClyA folding and functional assembly by lowering the energetic barrier for the monomer-to-MG transition and stabilizing the subsequent protomer formation.

### Molten Globule Intermediate State of ClyA Favors the Pore Formation in SM-CHOL Membranes

The rapid unfolding of ClyA upon membrane contact points to the existence of a transient’molten globule’ (MG) intermediate, reminiscent of unfolded monomer states previously observed during detergent-induced ClyA assembly kinetics (Benke *et al*., 2015). To structurally interrogate this elusive species, we exploited the slow folding kinetics of ClyA in n-dodecyl-β-D-maltoside (DDM), using the detergent as a kinetic trap to stabilize the intermediate. Validating this mimetic approach, atomistic MD simulations revealed that DDM preferentially invades the hydrophobic interface of the β-tongue region, mirroring the lipid-driven engagement observed in our membrane models (**Figure S16A,B**). This interaction effectively dismantles the ordered secondary structure; simulations of the isolated β-tongue motif within a DDM micelle displayed a pronounced collapse of helicity and a rise in structural deviation (RMSD ∼1 nm), consistent with the’melting’ required for insertion (Kulshrestha *et al*., 2023a) (**Figure S16C,D**). Energy decomposition further identified the conserved hydrophobic residues Y178 and F190 as the primary loci of DDM-induced instability, confirming that the detergent micelle faithfully recapitulates the hydrophobic destabilization forces of the lipid bilayer (Kulshrestha *et al*., 2023b).

To experimentally capture the functional properties of this transient species, we leveraged the slow assembly kinetics of ClyA in (0.1%) DDM to isolate distinct conformational snapshots (**Figure S17A**; (Benke *et al*., 2015)). Monitoring α-helical acquisition via circular dichroism (CD) allowed us to define three kinetic populations: a partially unfolded intermediate (’I’, at ∼1 min), an incomplete pore (’InP’, at ∼15 mins), and the predominantly complete pore (’cP’, 60 mins) (**Figure S17B,C**). Structural characterization with negative stain TEM confirmed the stability of these states, with arc-like assemblies appearing in InP and complete pores dominating at 60 minutes. On the other hand, ‘I’ state was predominantly a distinct, disordered monomeric species devoid of higher-order oligomers (**Figure 3A**). FTIR deconvolution revealed reduced α-helical and β-sheet content with increased disorder in the ‘I’ state, whereas both ‘InP’ and ‘cP’ exhibited increased α-helical content, reflecting progressive structural reorganization during assembly (**Figure 3B**, **Figure S17D**).

**Figure 3:**
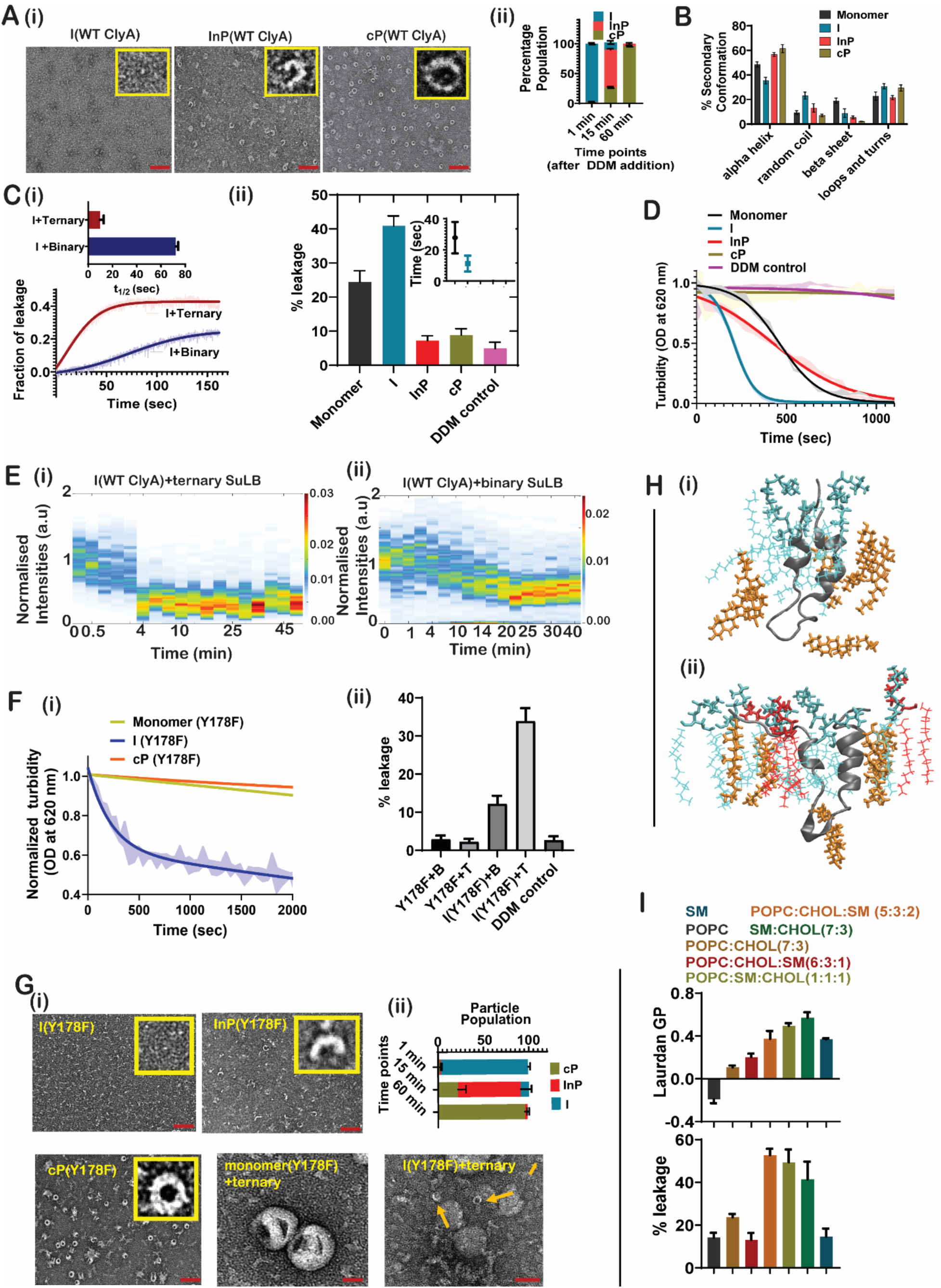
DDM-induced molten globule-like intermediate of ClyA exhibits sensitivity to ternary membrane compositions. (A) (i) Negative-stain TEM images showing distinct structural states of WT ClyA following DDM treatment: molten globule-like intermediate (‘I’, 1 min), incomplete pore (‘InP’, 15 min), and complete pore (‘cP’, 60 min). Insets show magnified views of representative particles from each state. Scale bar is 30 nm.(ii) Quantification of the relative populations of each state based on negative-stain TEM analysis. Data are presented as mean ± SD (n = 3). (B) Bar plot representing the percentage of secondary structural elements (α-helix, β-sheet, etc.) in each DDM-induced WT ClyA state, as determined from deconvolution of ATR-FTIR spectra. Data are shown as mean ± SD (n = 3). (C) (i) Calcein leakage kinetics from calcein-loaded liposomes composed of binary and ternary lipid mixtures following addition of the WT ClyA ‘I’ state. Inset shows the half-lives of leakage kinetics for each lipid composition, determined by Boltzmann sigmoidal fitting. Data are presented as mean ± SD (n = 3). (ii) Extent of calcein leakage from ternary liposomes upon treatment with various DDM-induced ClyA states, monomeric ClyA, and 1% DDM control. Inset shows half-lives of pore formation derived from leakage kinetics. Data are shown as mean ± SD (n = 3). (D) Turbidity-based hemolysis assay showing red blood cell (RBC) leakage kinetics in response to WT ClyA monomer and DDM-induced intermediate states. Data were represented as mean ± SD (n = 3). n denotes the number of replicates, and the shaded area indicates standard deviation. (E) 2D histograms representing SRB leakage from SuLB platforms: (i) ternary and (ii) binary lipid compositions treated with 100 nM ClyA ‘I’ state. Color bars indicate the probability density of leakage events across microwells. (F) (i) Turbidity-based RBC leakage kinetics after treatment with DDM induced Y178F mutant forms: monomer, ‘I(Y178F)’, and ‘cP(Y178F)’. Data were represented as mean ± SD (n = 3). n denotes the number of replicates, and the shaded area indicates standard deviation. (ii) Bar graph showing calcein release from ternary liposomes treated with DDM-induced Y178F states: ‘I(Y178F)’, ‘InP(Y178F)’, and ‘cP(Y178F)’. Data are shown as mean ± SD (n = 3). (G) (i) Negative-stain TEM images of DDM-induced Y178F mutant states (‘I(Y178F)’, ‘InP(Y178F)’, and ‘cP(Y178F)’), along with ternary liposomes treated with monomeric Y178F and ‘I(Y178F)’. Inset depicts magnified views of representative structures. Scale bar is 30 nm. (ii) Quantification of the relative populations of Y178F states, shown as mean ± SD (n = 3). (H) Snapshots of the molten globule-like intermediate ‘I’ interacting with (i) binary and (ii) ternary membrane systems. Lipid components are color-coded: PC (blue), cholesterol (orange), and sphingomyelin (red). (I) Bar plots showing (i) the percentage of calcein leakage induced by WT ClyA ‘I’ state across different lipid compositions, and (ii) generalized polarization (GP) values of the lipid phase-sensitive dye Laurdan in untreated liposomes of corresponding compositions. Data are shown as mean ± SD (n = 3).

Strikingly, functional mapping identified this disordered intermediate—not the mature pore or oligomers—as the primary agent of membrane permeabilization, as evidenced by a ∼7-fold higher calcein release from ternary liposomes compared to binary systems (**Figure 3C**). In contrast, InP and cP states failed to rupture membranes, indicating that the ternary lipid composition specifically facilitates membrane lysis by the unfolded intermediate state (I) (**Figure 3C**). However, liposomes possess high intrinsic curvature that can artificially lower the energetic cost of insertion. To rigorously validate this phenotype in a planar geometry devoid of curvature stress and substrate-induced artifacts, we deployed our high-throughput suspended lipid bilayer (SuLB) platform (Sannigrahi *et al*., 2022). Mirroring the vesicle data, the’I’ state drove rapid sulforhodamine B (SRB) efflux from ternary SuLBs, exhibiting a ∼3-fold increase in both pore formation rate and extent relative to binary bilayers (**Figure 3E**, **Figure S18A**). Crucially, the structurally mature’cP’ state exhibited negligible leakage in this artifact-free system (**Figure S18B**), confirming that lytic competence is transiently acquired during the unfolding phase and lost upon final oligomerization. This preferential activity of the unfolded ‘I’ species was also corroborated by red blood cell hemolysis assays where the intermediate species is even more potent than the structured monomer in lytic activity while fully assembled pores do not rupture the cells (**Figure 3D**).

The critical requirement for this “melting” transition was further illuminated by the behavior of the rigid, non-lytic Y178F mutant. While the mutant typically remains kinetically trapped, DDM exposure triggered a gradual acquisition of helicity akin to the wild-type assembly pathway (**Figure S18C**). Strikingly, this DDM-induced’I’ state displayed substantial lytic efficacy against ternary liposomes and erythrocytes (∼40% lysis)—a gain-of-function that was significantly attenuated in binary lipid environments (**Figure 3F**). Conversely, the fully matured’cP’ state remained inert in both functional assays (**Figure 3F**). Structural analysis by negative-stain TEM corroborated this functional recovery, revealing that the rescued’I’ intermediate successfully self-assembled into membrane-inserted oligomers, whereas the inactive cP state formed clustered assemblies indicative of reduced stability (**Figure 3G**). This chemical resurrection provides definitive mechanistic proof that the defect in Y178F stems specifically from its failure to transiently access the reactive molten globule intermediate required for membrane permeabilization.

Finally, we sought to define the biophysical features of the host membrane that potentiate this reactive intermediate. Atomistic snapshots of the MG-like state embedded in ternary bilayers visualized a striking preservation of lipid packing, contrasting with the disordered environment observed in binary systems (**Figure 3H**). This implies that the SM–CHOL complex provides a structured, ordered template that is essential for the intermediate’s function. We experimentally validated this “order-driven” hypothesis using Laurdan generalized polarization (GP) to map lipid packing across varying membrane compositions (Sannigrahi *et al*., 2024). Strikingly, the lytic activity of the’I’ state tracked tightly with membrane order: compositions enriched in SM and CHOL (particularly equimolar mixtures) exhibited the highest GP values and supported maximal pore formation (**Figure 3I**). This strong correlation confirms that the molten globule ‘I’ intermediate relies on the specific high-order architecture of SM–CHOL domains to drive efficient membrane rupture.

### ClyA Assembly Drives Reciprocal Membrane Remodeling And Lipid Scrambling

While our data establish that the SM–CHOL environment chaperones toxin folding, we reasoned that this interaction must be thermodynamically coupled: the energy released during protein refolding should, in turn, perform work on the lipid bilayer. To test this reciprocal remodeling hypothesis, we tracked the spatial organization of lipids during the β-tongue’s transition from the molten globule (MG) to the folded state.

Atomistic simulations revealed a distinct “lipid scrambling” event coincident with protein refolding. In the vicinity of the MG intermediate, sphingomyelin and cholesterol initially form tight complexes. However, as the protein matures towards the protomer state (∼400 ns), we observed a progressive increase in the distance between the sphingosine nitrogen of SM and the hydroxyl oxygen of cholesterol (**Figure 4A**). Visual analysis of the trajectory highlights this reciprocal dynamics: snapshots at 200 ns and 400 ns show that the structural maturation of the β-tongue is coupled to a spatial reorganization of the immediately adjacent SM–CHOL complexes (**Figure 4B**). This molecular separation is accompanied by a reduction in protein–lipid interaction energies (**Figure 4C**), suggesting that the folding polypeptide actively performs mechanical work to disrupt local SM–CHOL packing, effectively “loosening” the membrane to facilitate insertion. Such structural rearrangement is likely instrumental for ClyA N-terminal domain insertion, consistent with prior mechanistic models (Rojko & Anderluh, 2015).

**Figure 4:**
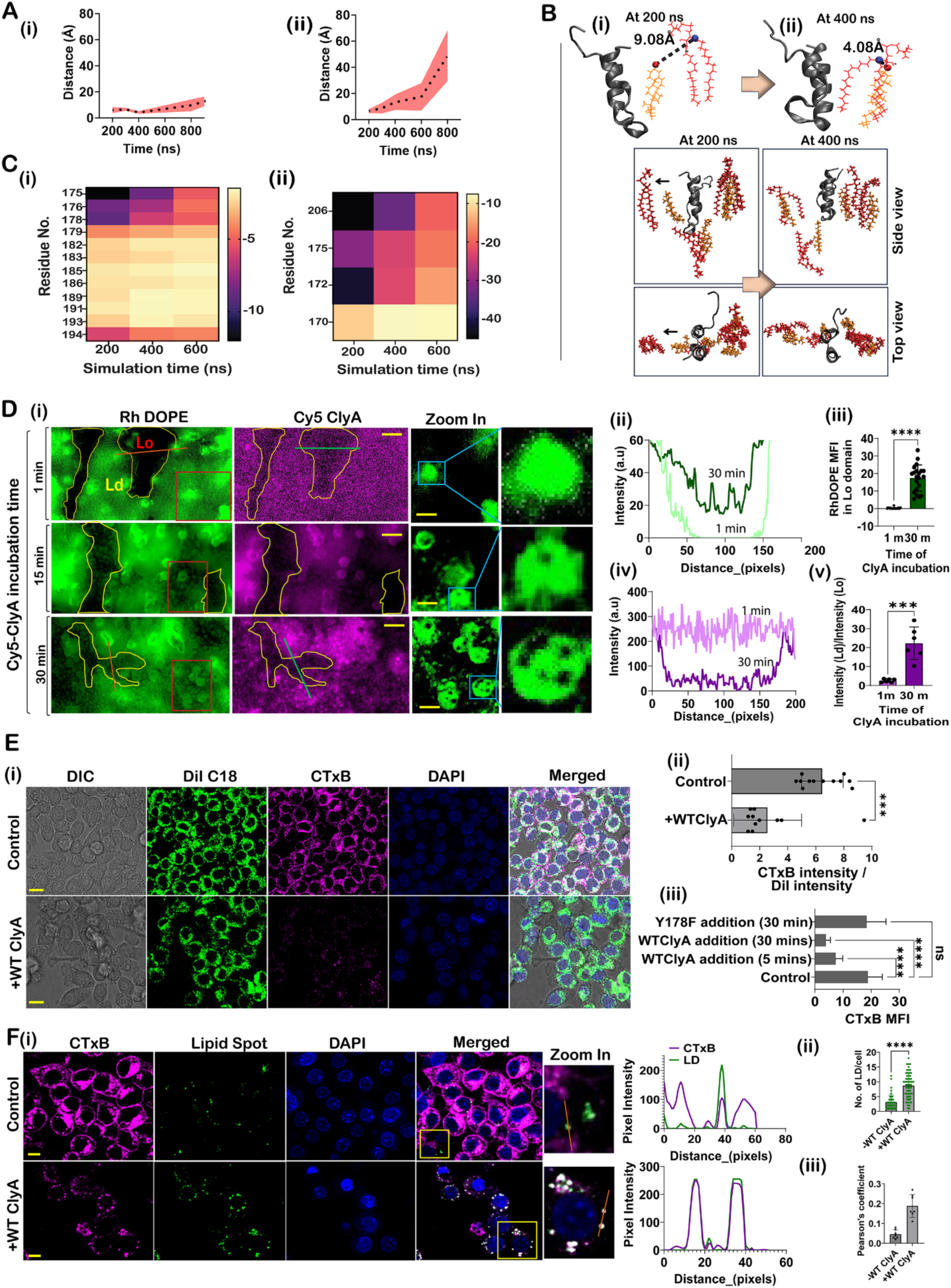
SM-CHOL complex behavior in ternary membranes during WT ClyA conformational transition. (A). (i) Distance between sphingosine nitrogen of SM and hydroxyl oxygen of CHOL during the conformational transition of the molten globule (MG)-like intermediate state of WT ClyA, as analyzed by all-atom MD simulations. Distances were measured for the SM-CHOL complexes located within a 15 Å radius of the WT ClyA MG state and the data were presented as mean ± SD from three simulation replicates. (ii) Distances between SM and CHOL (sphingosine nitrogen and CHOL hydroxyl oxygen) measured in ternary bilayers in regions distant from the WT ClyA MG state. Data were presented as mean ± SD from three simulation replicates. (B) Representative snapshots showing the MG-like state of WT ClyA and nearby SM-CHOL complexes at 200 ns and 400 ns of the MD simulation. (C) (i) Heatmap showing residue-wise interaction energies between the MG-like state and CHOL across three different MD time points in the ternary membrane. (ii) Heatmap of residue-wise interaction energies between the MG-like state and SM at the same time points. Color bars represent interaction energy in kcal/mol. (D) Ternary membrane composition was used to generate SuLB platforms with visible liquid-ordered (Lo) and liquid-disordered (Ld) phase separation, visualized via Rhodamine-DOPE labeling. (i) Fluorescence micrographs show SuLB platforms exposed to 50 nM Cy5-labeled ClyA for 1, 15, and 30 minutes. Dimmer Lo domains are marked in yellow. Right panels show zoomed-in views (red boxes). Scale bars: 5 µm (main images), 3 µm (zoomed-in). (ii) Line scan profiles of Rhodamine-DOPE fluorescence along red lines at 1 and 30 minutes post Cy5-ClyA exposure. (iii) Quantification of mean Rhodamine-DOPE fluorescence intensity in Lo domains at different time points. Data are shown as mean ± SD (n = 3). ****P < 0.0001. (iv) Line scan profiles of Cy5-ClyA fluorescence along green lines at 1 and 30 minutes of exposure. (v) Partition of Cy5-ClyA in Ld domains at 1 and 30 minutes post incubation. Data are shown as mean ± SD (n = 3). ****P < 0.0001. (E) Lipid raft organization in RAW 264.7 cells visualized using the raft marker CTxB (cholera toxin B). (i) Confocal images of CTxB and DiI C 18 labeled cells with or without WT ClyA treatment. DAPI (blue) marks nuclei. (ii) Plot of CTxB mean fluorescence intensity (MFI) per cell membrane with or without WT ClyA incubation. Data are presented as mean ± SD. ***P < 0.0023. (iii). Plot of CTxB mean fluorescence intensity (MFI) per cell membrane with or without WT ClyA incubation for different time points (5 mins and 30 mins) or Y178F incubation for 30 mins. Data are presented as mean ± SD. ***P < 0.0001. Scale bar is 15 μm. (F) (i). Confocal images showing lipid raft (CTxB, magenta), LD marker (lipid spot, green) and nucleus (DAPI,blue) with or without WT ClyA treatment. Scale bar is 10 μm. Line scan profiles (orange solid lines) show overlapping fluorescence intensities and the intensity plots (right) show the overlapping intensity profiles of raft (magenta) and LD (green).Scale bars: 10 µm. (ii). Plot showing the number of LD/cell in Raw cell with or without WT ClyA treatment. (iii). Plot shows the Pearson’s coefficient for CTxB and LD signal with or without WT ClyA treatment.

To visualize this remodeling at the macroscopic scale, we utilized our ternary SuLB platform, where segregated liquid-ordered (Lo) and liquid-disordered (Ld) domains can be imaged by fluorescence (**Figure 4D**). SuLB was labeled with the liquid-disordered (Ld) domain-specific fluorophore Rhodamine-DOPE (RhDOPE). Time-lapse imaging revealed that ClyA binding triggers a dramatic reorganization of this phase architecture. Upon toxin exposure, we observed the emergence of Ld-like patches within previously stable Lo domains and a concurrent redistribution of the Lo-marker Rhodamine-DOPE (**Figure 4D**). Additionally, small Lo patches emerged on the suspended areas of SuLB after 30 minutes (zoomed inset in **Figure 4Di**). These findings suggest that ClyA induced domain disruption and lipid mixing on ternary SuLB membranes. Temporal analysis indicated that ClyA initially displayed no selectivity for Lo or Ld domains, but with prolonged exposure, segregated to Ld regions (**Figure 4Di, iv, v**). Quantitatively, the toxin increasingly partitioned into the disordered phase over time, confirming a physically disruptive “mixing” of lipid domains.

This phenomenon extends to the native cellular context. we also examined the temporal dynamics of lipid domain architecture of RAW cells upon exposure to WT ClyA (50 nM). In RAW 264.7 macrophages, WT ClyA treatment significantly reduced the surface binding of Cholera Toxin B (CTxB), a marker for GM1-rich lipid rafts (**Figure 4Ei–ii**). Crucially, this raft disruption was absent in cells treated with the non-lytic Y178F mutant, confirming that the remodeling is driven specifically by the conformational work of the folding toxin (**Figure 4Eiii**, **Figure S19A**). The reduction in CTxB signal was not attributable to membrane rupture, as DiI C-18 fluorescence confirmed that membrane integrity was not compromised (**Figure 4E**). The simultaneous decrease in CTxB binding following WT ClyA exposure suggests that ClyA’s conformational changes upon membrane insertion are closely linked to reorganization of lipid domains in the plasma membrane. Furthermore, tracking of fluorescently labeled ClyA revealed a dynamic translocation: the toxin initially colocalizes with raft markers (5 min) but subsequently migrates to non-raft regions (30 min) as it disrupts the local order (**Figure S19B, C**) (Sarangi & Basu, 2018).

Finally, we confirmed this toxin-induced fluidization using phase-sensitive Laurdan generalized polarization (GP). Laurdan fluorescence spectra of untreated membranes exhibited a single peak at 440 nm (**Figure S19D**). However, both in ternary liposomes and live cells, WT ClyA treatment induced a marked spectral redshift and a corresponding reduction in GP values, indicative of increased membrane fluidity (**Figure S19D, E**). Time-dependent reduction in Laurdan GP confirmed increased membrane fluidity induced by WT ClyA (**Figure S19D**), in agreement with the CTxB imaging. In contrast, the rigid Y178F mutant failed to alter membrane order (**Figure S19D, E**). Collectively, these findings suggest that ClyA disrupts cellular lipid raft structures during its assembly, resulting in lipid mixing and domain redistribution through increased lipid mobility in the bilayer.

The physiological consequences of this lipid scrambling are profound. We observed that the dismantling of plasma membrane rafts by ClyA triggers the rapid biogenesis of intracellular lipid droplets (LDs). In particular, RAW cells exposed to WT ClyA exhibited a pronounced increase in lipid droplet (LD) formation (**Figure 4Fi, ii**). Following toxin exposure, surface-derived raft components (CTxB-positive) were internalized and relocalized to nascent lipid droplets (**Figure 4Fi, ii, iii**). This suggests that raft-associated sphingomyelin-cholesterol (SM-CHOL) domains could be relocalized into nascent LDs upon toxin exposure (Gonzalez *et al*, 2011; Lakshminarayan *et al*, 2014). We conjecture that the “lipid chaperone” mechanism comes at a metabolic cost to the host− by exploiting the SM–CHOL gradient to fold, the toxin also mechanically destabilizes the plasma membrane, forcing the cell to sequester displaced lipids into storage organelles to maintain homeostasis.

### SM–CHOL Complexes Engage Conserved Lysine Sensors To Drive Productive Membrane Insertion

The profound impact of membrane composition on ClyA activation suggested that the SM–CHOL complex provides a specific chemical signature recognized by the toxin. Prior studies have established lysine residues as key mediators of protein–lipid interactions; lysine side chains can preferentially associate with phospholipid headgroups and, in the context of pore-forming toxins such as lysenin, are critical for sphingomyelin binding and membrane disruption (Li *et al*, 2013; Yamaji *et al*, 1998; Yilmaz *et al*, 2018). Therefore, we hypothesized that specific interactions between SM-CHOL and β-tongue lysines are essential for the refolding events that govern ClyA lytic activity. We identified three specific residues—K172, K175, and K206—that showed significant interactions with SM during the transition from the MG-like state of β-tongue to the protomer-like conformation in our MD simulations (**Figure 5Ai-iii**). Notably, during the initial interaction phase, K175, K172, and K206 exhibited distinct interaction patterns with SM and CHOL (**Figure 5A**). Energy decomposition revealed a distinct hierarchy of interaction: K175 emerged as a dual-specificity sensor, maintaining robust, high-affinity interactions with both SM and CHOL (E_SM_=-23.52Kcal/mol; E_CHOL_=-8.6 kcal/mol) throughout the trajectory (**Figure 5A ii**). In contrast, K206 functioned primarily as an SM-anchor (E_SM_=-33.52Kcal/mol), while K172 exhibited negligible interaction with CHOL (E_CHOL_=-0.37 Kcal/mol), despite having a similar interaction with SM as K175(E_SM_=-22.87 Kcal/mol). Temporal analysis revealed that the SM-CHOL complex maintained consistent interaction with K175 throughout the refolding process. However, the interaction energy between SM and both K172 and K206 decreased as refolding progressed (**Figure 5Aii**).

**Figure 5:**
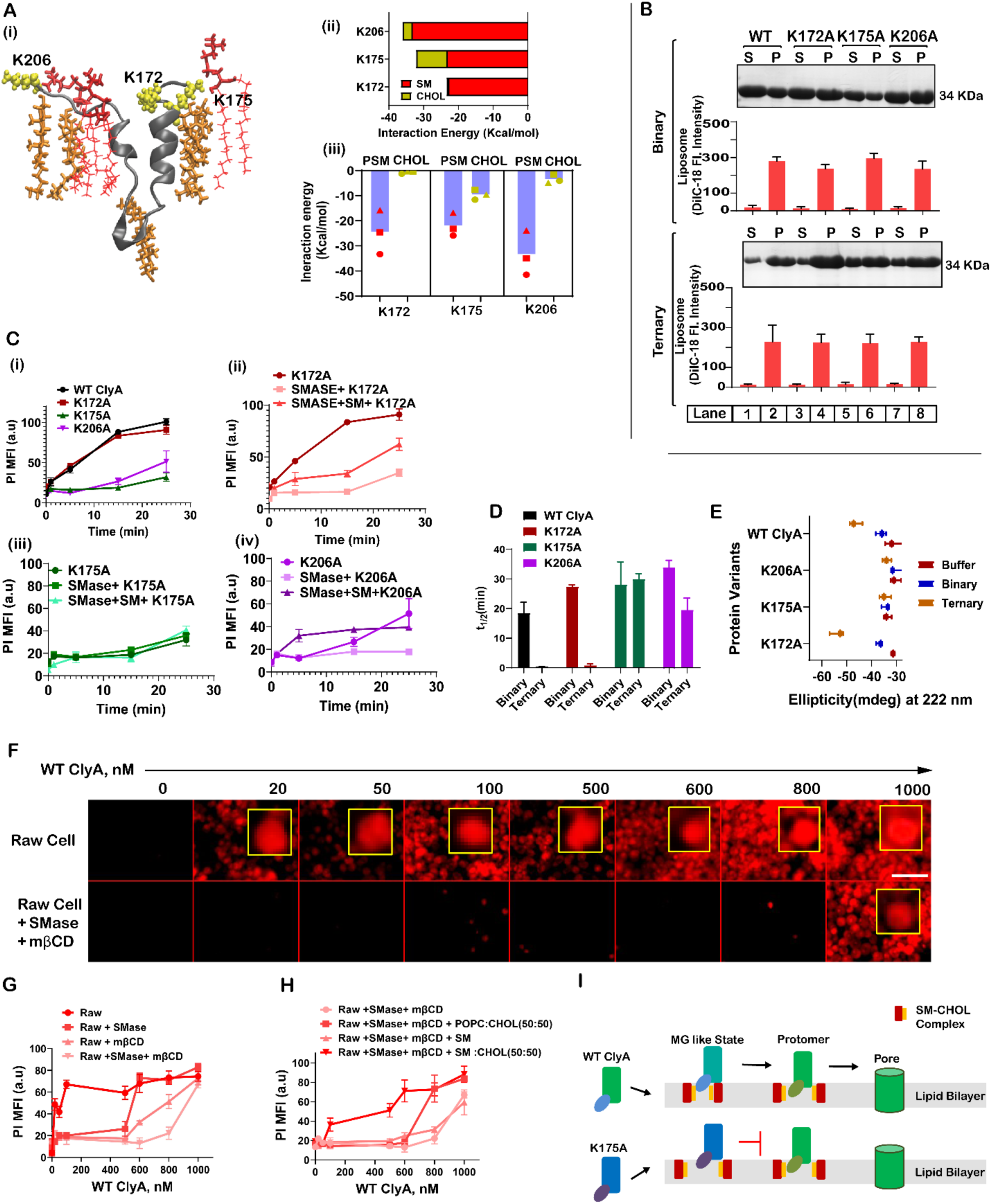
SM-CHOL complex is essential for the conformational transition and membrane rupture activity of ClyA. (A) (i) A snapshot from the 200 ns time point of a molecular dynamics simulation illustrates the SM-CHOL complex within a ternary membrane, localized near the peripheral region of the MG-like state of the B-tongue segment. Sphingomyelin (SM) is shown in red, cholesterol (CHOL) in yellow, and the mutation sites (K172A, K175A, and K206A) are highlighted as blobs. (ii) Interaction energy plot at 450 ns of simulation timepoint (where the jump of alpha helical conformation of the molten-globule segment happens) depicts the binding energies of different lysine residues of MG like state with PSM and CHOL in ternary lipid bilayer. (iii). Nested graph indicates the interaction energy of PSM and CHOL with three lysine residues at different simulation tie points (round symbol denotes 200 ns, rectangular symbol for 400 ns and triangle symbol indicates 600 ns of simulation time points). Each data point is presented as the average of three individual datasets obtained from three simulation repeats. (B) Sedimentation assays evaluate the binding affinity of the three mutants with binary and ternary multilamellar vesicles (MLVs). Bar plots display DiI fluorescence intensity in the pellet (P) and supernatant (S) fractions under various conditions. Data are shown as mean ± SD (n = 3). (C) Pore-forming activity in RAW 264.7 cells was assessed by measuring propidium iodide (PI) uptake over time using mean fluorescence intensity (MFI) of captured images under different conditions: (i) WT and mutants K172A, K175A, and K206A; (ii–iv) Each mutant was tested with and without sphingomyelinase (SMase) pretreatment, and sequential SMase followed by SM liposome addition. Data represent mean ± SD (n = 3). n stands for the number of experimental repeats. (D) Half-lives of calcein release from binary and ternary liposomes were determined following exposure to WT ClyA and the three mutants. Data are presented as mean ± SD (n = 3).n denotes the number of experimental repeats. (E) Circular dichroism (CD) spectroscopy was used to monitor secondary structural changes in WT ClyA and the mutants in the presence and absence of binary or ternary liposomes. (F) Fluorescence micrographs of Raw cells PI uptake with increasing concentration of WT ClyA in absence and presence of sequential prior treatments of SMase (50 mU/mL) and mβCD (1 mM) before WT ClyA exposure. (G). Plot shows the PI uptake inside Raw cells by WT ClyA at different prior treatment conditions. (H). A recovery experiment on the doubly treated cells (first SMase and then mβCD treatment) to show the effect of the addition of liposomes consisted of different lipid components prior to WT ClyA addition. (I) A schematic illustration summarizes the mechanism: the SM-CHOL complex interacts with the MG-like conformation of the β-tongue region in WT ClyA, facilitating conformational refolding and membrane rupture via pore formation. In contrast, the K175A mutant fails to engage with the SM-CHOL complex in its MG state, preventing conformational transition and pore formation.

To validate this sensory mechanism, we generated alanine substitutions (K172A, K175A, K206A). Biophysical analysis confirmed that all three mutants (K172A, K175A, and K206A) retained conformational stability indistinguishable from WT ClyA, as assessed by DynaMut modeling(Rodrigues *et al*, 2018) and circular dichroism (CD) spectroscopy (**Figure S20**).

Sedimentation assays showed that each mutant is bound efficiently to both binary and ternary multilamellar vesicles (**Figure 5B**). Functionally, however, the mutations revealed a stark dichotomy. K175A and K206A exhibited a profound loss of lytic activity in PI uptake assays, whereas K172A retained wild-type potency and sensitivity to SMase treatment (**Figure 5Ci**). Notably, K172A activity was abrogated by sphingomyelinase (SMase) treatment but recovered upon reintroduction of SM, paralleling WT ClyA behavior (**Figure 5Cii**). In contrast, K175A remained functionally impaired under all tested membrane compositions, and K206A showed partial sensitivity to membrane lipid composition (**Figures 5Ciii, iv**). Calcein leakage assays further revealed increased leakage by K172A in ternary membranes, whereas K175A and K206A mutants displayed significantly reduced activity across all membrane types (**Figure 5D**). Structural analysis by CD indicated that only K175A failed to undergo the membrane-induced conformational changes typical of ClyA, while K172A exhibited increased α-helicity upon membrane binding and K206 showed slight increase in helicity after liposome binding (**Figure 5E, Figure S20C**). Collectively, these findings identify K175 and K206 as critical for the SM-CHOL mediated MG-to-protomer transition, which is essential for efficient membrane lysis by ClyA.

Finally, we interrogated the physiological relevance of this lipid requirement in live cells (**Figure 5F**). RAW cells depleted of CHOL and SM using methyl-β-cyclodextrin (mβCD) and sphingomyelinase (SMase) were monitored for ClyA lytic activity. mβCD treatment caused the strong attenuation of PI uptake, but sequential SMase + mβCD treatment produced the strongest inhibition of WT ClyA activity (**Figure 5G**). Recovery experiments revealed that only SM-CHOL (50:50) liposomes could efficiently restore lytic activity, whereas POPC-CHOL or SM alone failed to support pore formation (**Figure 5H**). Corroborating this specificity, an external competition assay showed that free SM–CHOL liposomes effectively acted as a “decoy,” sequestering the toxin and preventing cell lysis (**Figure S21**). Together, these results demonstrate that the SM–CHOL complex is not merely a passive receptor, but an indispensable cofactor that engages lysine sensors to trigger the lytic cascade (**Figure 5I**).

### Conserved Cationic Sensors and Hydrophobic Runs Define a Lipid-Guided Membrane Insertion Code

The strict requirement for specific lysine–lipid contacts in ClyA assembly points to a broader structural logic governing membrane recognition. Evolutionary analysis reveals that the essential sensor residues (K175 and K206) are strictly conserved across diverse bacterial ClyA homologs (**Figure 6A**) preserving a “molecular barcode” designed to detect ordered lipid environments. This barcode comprises a bipartite motif: the conserved cationic sensors that anchor the protein to the interface, and a contiguous 16-residue hydrophobic run within the β-tongue that executes the insertion (**Figure 6Bi**). We propose that these elements function in concert to target the toxin specifically to SM–CHOL-rich domains.

**Figure 6:**
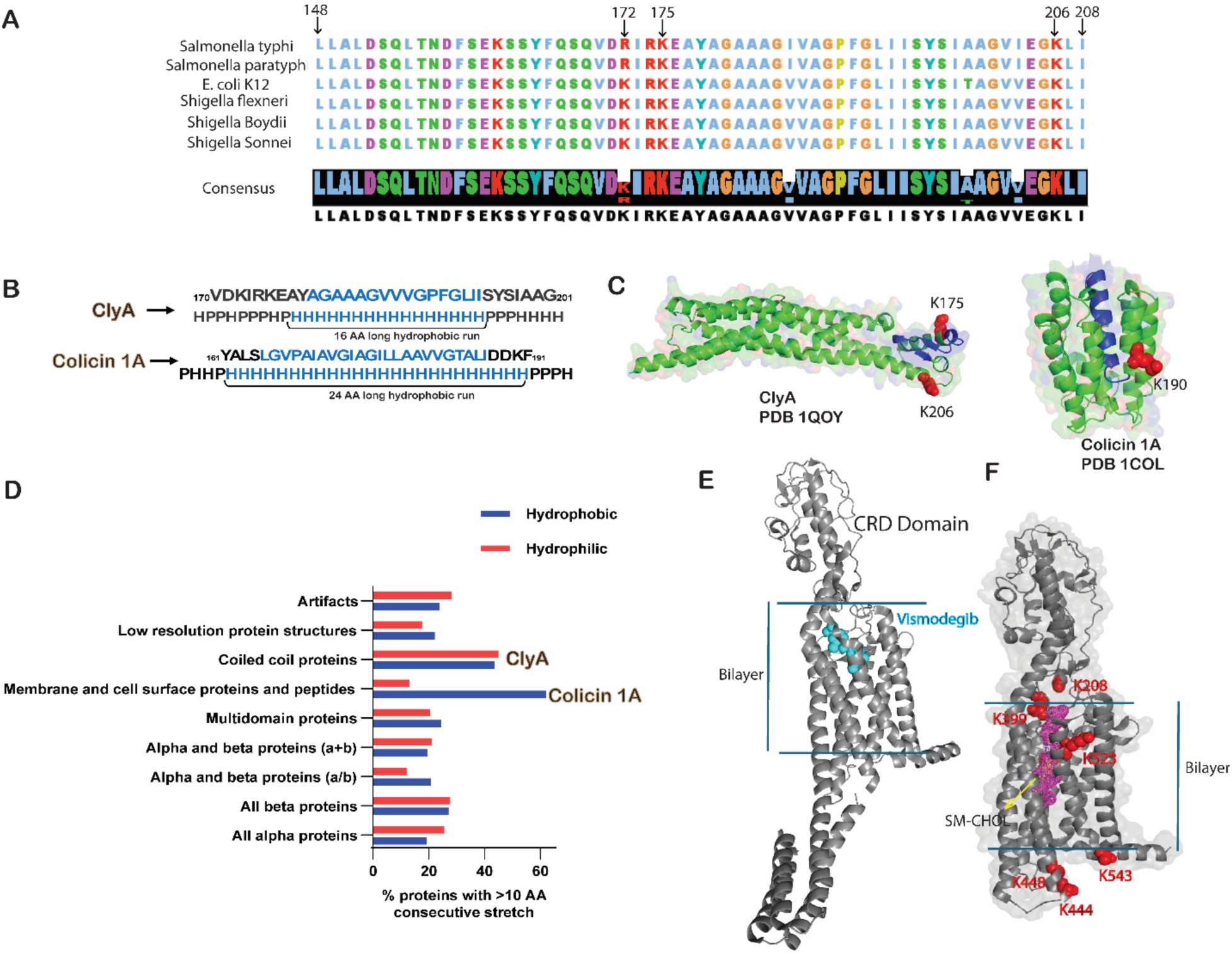
Conservation of cationic sensors and hydrophobic runs. **(A)** Sequence alignment of ClyA from diverse bacterial species reveals strong conservation of the cationic lysine residues K175 and K206. **(B)** Comparison of hydrophobic run sequences within the pore-forming domains of ClyA and colicin 1A.**(C)** Crystal structures of ClyA (PDB: 1QOY) and colicin 1A (PDB: 1COL), highlighting hydrophobic runs (blue) and cationic lysine sensors (red). **(D)** Frequency analysis of hydrophobic and hydrophilic run sequences derived from the SCOP protein class database. According to SCOP classification, ClyA belongs to the coiled-coil family, whereas colicin 1A is classified within membrane and cell-surface proteins. **(E)** Structure of Smoothened bound to the agonist vismodegib (PDB: 6O3C).**(F)** Molecular docking–derived model of Smoothened bound to SM-CHOL. Docking was performed using AutoDock Vina via the SwissDock platform with a docking grid defined by coordinates −3, 4, 18 to 22, 30, 24.

Strikingly, this “sensor-and-piston” architecture appears to be a convergent solution for membrane penetration. A parallel arrangement is found in the pore-forming toxin Colicin 1A, which features an extended 23-residue hydrophobic run flanked by a critical lysine residue (**Figure 6B–C**). Biophysically, this enrichment of lysines at the polar–apolar interface (Landolt-Marticorena *et al*, 1993; von Heijne, 1994), allows their side chains to “snorkel,” effectively locking the transmembrane segments in position while stabilizing lipid interactions (Gebhardt *et al*, 2012; Segrest *et al*, 1990; Strandberg & Killian, 2003). Our SCOP-based bioinformatic survey confirms that such long hydrophobic runs (Schwartz & King, 2006) are a genomic hallmark of membrane-associating proteins (**Figure 6D**). We argue that in spite of ClyA’s classification as a soluble coiled-coil protein, the hydrophobic β-tongue functions as a canonical membrane-insertion module.

Does this reliance on SM–CHOL dynamics extend beyond bacterial toxins? The principle of SM-mediated cholesterol regulation appears to be conserved in mammalian signaling, specifically in the Hedgehog pathway transducer Smoothened (SMO). SMO is activated by CHOL binding to its cysteine-rich domain (CRD) yet avoids constitutive activation despite high plasma membrane CHOL abundance (Radhakrishnan *et al*, 2020; Siebold *et al*, 2016). While structural studies confirm cholesterol binding sites in the SMO transmembrane (TM) cavity and CRD (Siebold *et al*., 2016) (**Figure 6E**), the regulatory trigger has remained elusive.

Applying the mechanistic insights from ClyA, we explored whether SM acts as a “capacitive buffer” for SMO. Molecular docking reveals that the SM–CHOL complex preferentially occupies the deep TM cavity of SMO rather than the bilayer interface (**Figure 6F**). We propose a model where SM inhibits signaling by sequestering cholesterol into stable complexes within the TM domain, preventing its transfer to the activating CRD site. Much like in ClyA, where the SM–CHOL interaction dictates the conformational switch, we suggest that lysine-rich regions in SMO read the SM–CHOL state to gate activity. Together, these findings posit that the coupling of conserved cationic sensors with hydrophobic motifs represents a fundamental, cross-kingdom mechanism for sensing and responding to membrane lipid composition.

## Discussion

Our findings resolve the paradox of how a soluble protein breaches a hydrophobic barrier, demonstrating that the host membrane acts as a conformational chaperone rather than a passive receptor. It sheds light on the longstanding disparity between ClyA’s slow kinetics in reductionist models (e.g., detergent micelles) versus its rapid efficacy against biological membranes like those of erythrocytes and macrophages. While classical models depict pore formation as a spontaneous insertion event, we demonstrate that the host membrane acts as an obligatory conformational chaperone. Specifically, ternary membranes mimicking the plasma membrane outer leaflet drive an 18-fold enhancement in lytic activity and a ∼5-fold acceleration in pore formation kinetics compared to binary controls. This dramatic potentiation occurs because the ternary environment resolves the energetic bottleneck of the transition. In binary membranes, the β-tongue remains frustrated, trapped in a shallow energy basin characterized by incomplete folding. The synergy of sphingomyelin and cholesterol (SM–CHOL) resolves this bottleneck by actively lowering the energetic barrier for the structural metamorphosis of the β-tongue. Thus, the SM–CHOL environment functionally mimics molecular chaperones, substituting chemical energy with the thermodynamic potential of the lipid bilayer.

The molecular mechanism of this lipidic chaperoning of ClyA is distinct from simple electrostatic recruitment and follows a sequential “unfold-to-refold” trajectory. First, interaction with CHOL catalyzes the local “melting” of the ClyA monomer’s hydrophobic β-tongue into a “molten globule” (MG) intermediate. This transient access to the MG intermediate, a disordered state that bridges the soluble and membrane-inserted forms is central to functional pore formation. This is exemplified by the rigid mutant Y178F that fails to melt its β-tongue and is inactive even though it assembles into a ring-shaped complex. However, we demonstrate that chemical induction of disorder using DDM effectively “rescues” its lytic potential, allowing the mutant to bypass the kinetic trap and assemble into functional pores. This supports our hypothesis that an ‘unfolded β-tongue’ intermediate is an obligatory step for functional pore formation (**Figure S22**). Further, SM assists the refolding of this intermediate into the functional helix-turn-helix motif by competitively engaging the cholesterol ligands of ClyA.

The mammalian plasma membrane functions as a protective “biophysical rheostat” (Simons & Ehehalt, 2002), where sphingomyelin sequesters cholesterol into ordered, low-activity complexes to defend against virulence factors. Because of this sequestration (Das *et al*., 2014), typical cholesterol-dependent cytolysins (CDCs)—such as LLO and PFO—are often unable to engage sterols on the native surface and require co-secreted enzymes (e.g., SMase) to chemically liberate accessible cholesterol for binding (Flores-Díaz *et al*, 2016; Heuck *et al*, 2010). ClyA, however, exemplifies a divergent evolutionary strategy that overcomes this barrier without enzymatic assistance. Lacking intrinsic lipid-modifying machinery (**Figure S23-25**), the toxin instead utilizes the sequestered SM–CHOL complex itself as a folding chaperone. This interaction is thermodynamically reciprocal: the free energy released during toxin refolding performs mechanical work on the bilayer, actively “scrambling” the local architecture to dismantle liquid-ordered domains and increase membrane fluidity. By forcing a transition from ordered to disordered states, the assembling pore mechanically disrupts the SM–CHOL sequestration equilibrium, effectively liberating cholesterol and imposing a metabolic cost that triggers the rapid biogenesis of lipid droplets to restore homeostasis.

The specificity of this interaction is encoded by a bipartite structural motif: a set of conserved cationic sensors coupled to a hydrophobic insertion sequence. We identified K175 and K206 as the obligate readers of this lipid code. K175 functions as a dual-specificity sensor, maintaining high-affinity contacts with both SM and CHOL to orient the folding domain, whereas K206 serves as an electrostatic anchor. These sensors work in concert with the contiguous 16-residue hydrophobic run of the β-tongue to target SM–CHOL-rich domains. The strict conservation of these residues across diverse ClyA homologs underscores their evolutionary importance. they constitute a “molecular barcode” that allows the toxin to distinguish the ordered, cholesterol-rich membranes of eukaryotic hosts from bacterial bilayers. The mechanism defined here could reflect a general principle of membrane protein folding and lipid interaction that extends beyond pathogenesis. Further, it suggests that pathogenic toxins have evolutionarily optimized a “cationic sensor” motif to detect the specific electrostatic and packing signature of SM-rich domains. By targeting SM—which acts as the physiological regulator of CHOL pool reactivity—ClyA bypasses the need for random collision, instead homing in on the specific membrane reservoirs where the thermodynamic potential for folding is highest.

This lipid-guided insertion logic appears to be an evolutionarily conserved strategy, extending beyond bacterial toxins to mammalian signaling machinery. A parallel “sensor-and-piston” architecture is observed in Colicin 1A, where flanking lysines position a hydrophobic run at the interface. Furthermore, we propose that this mechanism offers a new framework for understanding cholesterol sensing in the Hedgehog pathway transducer Smoothened (SMO). Much like ClyA, SMO contains cholesterol-binding sites buried within its transmembrane domain. Our docking analysis suggests that SM may inhibit SMO by sequestering cholesterol into stable complexes, thereby preventing its transfer to the activating cysteine-rich domain. Thus, the interplay between conserved cationic sensors, hydrophobic motifs, and the physical chemistry of the SM–CHOL complex represents a generalizable principle for regulating membrane protein function across kingdoms. This striking mechanistic convergence implies that bacteria have likely co-evolved to exploit a pre-existing eukaryotic regulatory code: the same thermodynamic principles that cells use to tune receptor sensitivity via lipid composition are hijacked by pathogens to drive the lethal assembly of virulence factors.

## Methods

### Computational Methods

#### Simulations of ClyA in DDM

The crystal structure of the ClyA monomer was taken from the PDB ID (1QOY). The initial structures of solvated monomers with water and two different numbers of DDM (n-Dodecyl β-D-maltoside) molecules, i.e., 50 and 100 molecules, were generated using CHARMM-GUI. An initial solvated structure of β-tongue and beta-tongue exposed to the micelle of 100 DDM molecules were also prepared using CHARMM-GUI, where β-tongue consisting of 38 residues (170-207) was truncated from the monomer structure. Details of all simulation setups are presented in Table S1.

All initial structures were solvated with TIP3P water and 𝑁𝑎^+^ and 𝐶𝑙^-^ ions were added to maintain the charge neutrality and 150 mM of physiological salt concentration. Initial solvated structures were energy minimized using the steepest. The energy minimized structures were then gradually heated up to 310 K, followed by 10 ns of NVT simulation and 20 ns of NPT simulations with position restraints on protein, and DDM molecules. The final configurations from the equilibration simulations were used as the starting structures for the production runs.

All simulations were performed in the NPT ensemble at 310 K using GROMACS-2018.6 with the force field parameters for the ClyA protein taken from CHARMM36 force field. The isotropic pressure control was achieved using the Parrinello-Rahman method with a time constant of 5 ps. Pressure was kept constant at 1 bar using the isothermal compressibility as 4.5 x 10^-5^ bar^-1^. The temperature was controlled using Nose-Hoover thermostat with a time constant of 1.0 ps. The solvent, protein, and DDM molecules were coupled separately to a temperature bath at 310 K. The long-range electrostatic interactions were treated using the particle mesh Ewald (PME) method(Essmann *et al*, 1995) with a real space cut-off of 1.2 nm and short ranged interactions were computed considering a cut-off radius of 1.2 nm. Hydrogen bonds were constrained using the LINCS constraint, which allowed a larger time step of 2 fs. Equilibration was monitored by evaluating the root mean squared deviation (RMSD) and secondary structure changes of the protein.

#### MD Simulation in lipid

The initial crystal structures for the Cytolysin A (ClyA) pore and monomer were retrieved from the Protein Data Bank under the PDB IDs: 2WCD (pore) and 2QOY (monomer). To focus on the change in conformation of the protein, the beta-tongue segment of the monomer (amino acid residues 170–207) was truncated based on insights from prior research. A previously reported melted beta-tongue segment corresponding to this truncated region (170–207) was also studied(Kulshrestha *et al*., 2023b).

The ClyA pore was embedded in two distinct lipid bilayers: (a) POPC:CHOL at 70:30 ratio and (b) POPC:SM:CHOL at 1:1:1 ratio, both constructed using the CHARMM-GUI membrane builder(Jo *et al*, 2009; Wu *et al*, 2014). Each setup was solvated with TIP3P (Jorgensen *et al*, 1983)water molecules, creating a water layer on either side of the membrane. A physiological salt concentration of 0.15 M KCl was added to mimic a biologically relevant ionic environment. Similar protocols were followed for systems incorporating each of the truncated beta-tongue segment and the molten globule segment within both lipid bilayers. Details of all simulation setups are presented in Table S2. AMBER20 (Case *et al*, 2023)package is used for all the simulations. The topology and coordinate files for all systems were prepared in AMBER format using CHARMM-GUI, with the ff14SB (Maier *et al*, 2015) force field applied to the protein and the lipid21 (Dickson *et al*, 2022)force field for the lipid molecules. Ion parameters for K⁺ and Cl⁻ were defined according to the Joung and Cheatham model(Joung & Cheatham III, 2008).

To optimize the systems, a two-step energy minimization protocol was applied. Initially, a 20,000-step minimization was performed with position restraints on the protein and lipid bilayers to ensure optimal water molecule placement. This was followed by another 20,000-step minimization without restraints, allowing full relaxation of the system. Each minimization phase employed the steepest descent method for the first 10,000 steps, followed by the conjugate gradient method for the remaining steps. Gradual heating was performed by linearly increasing the temperature from 0 K to 310 K over 2 ns in the NVT ensemble, with position restraints applied to the protein and lipids. Subsequently, the systems underwent equilibration for 10 ns under the same conditions, followed by an additional 4 ns equilibration in the NPT ensemble at 310 K and 1 atm, with weak restraints on the backbone atoms of the protein. Production simulations were then carried out for 50 ns in the NPT ensemble with these weak restraints, followed by unrestrained 1 μs NPT production runs to capture detailed dynamics. Each system was simulated in triplicate to minimize statistical errors.

Pressure regulation was maintained using a Monte Carlo barostat (Åqvist *et al*, 2004) with a relaxation time of 2 ps, while temperature control was achieved using a Langevin thermostat (Hünenberger, 2005) with a collision frequency of 1 ps⁻¹. Semi-isotropic pressure scaling was applied to maintain a consistent surface tension, with the bilayer plane aligned along the XY axis(Omelyan & Kovalenko, 2013; Zhang *et al*, 1995). The SHAKE algorithm(Ryckaert *et al*, 1977) was employed to constrain hydrogen-involved covalent bonds, enabling a 2-fs integration time step. Short-range nonbonded interactions were truncated at 10.0 Å, while long-range electrostatics were calculated using the particle mesh Ewald (PME) method(Essmann *et al*., 1995). Periodic boundary conditions (PBC) were applied in all three directions. The trajectories were analyzed using the CPPTRAJ (Roe & Cheatham III, 2013)module of Amber20 package and Visual Molecular Dynamics (VMD) (Humphrey *et al*, 1996)was used for visualization purposes.

### DynaMut for predicting the impact of mutations

DynaMut(Rodrigues *et al*., 2018) combines graph-based structural signatures with normal mode analysis (NMA) to provide a consensus prediction of how mutations affect protein stability. NMA enables the exploration of harmonic motions within a system, offering valuable insights into protein dynamics and accessible conformational states. Unlike more computationally demanding molecular dynamics (MD) simulations, NMA offers a computationally efficient alternative for studying protein flexibility and motion. While MD generates detailed motion trajectories over time, NMA evaluates conformational fluctuations by analyzing normal modes (eigenvectors) and their corresponding frequencies (eigenvalues). To further reduce computational cost, NMA can use simplified protein representations, such as modeling residues with only their Cα atoms. This method has proven effective for studying the dynamic effects of mutations. Notably, ENCoM enhances NMA by incorporating amino acid-specific properties to evaluate how single-point mutations influence vibrational entropy (ΔS) and protein stability.

### Analysis of hydrophobic and hydrophilic runs in protein databases

The prevalence hydrophobic and hydrophilic in the various classes of proteins in the SCOP (Structural Classification of Proteins; https://scop.berkeley.edu/) and OPM (Orientations of Proteins in Membranes; https://opm.phar.umich.edu/) (Lomize *et al*, 2012) and DisProt (database of intrinsically disordered proteins; https://disprot.org/) databases were calculated as follows: All the sequences in each database were downloaded. Each sequence was converted into a binary string by mapping the hydrophilic (Cys, Asp, Glu, His, Lys, Asn, Gln, Arg, Ser, Thr, Tyr) and hydrophobic (Ala, Phe, Gly, Ile, Leu, Met, Pro, Val, Trp) residues to 0 and 1, respectively(Moore & Stanitski, 1998; Seddon, 1988). The lengths of the consecutive runs of 0 and 1 were counted in each binary string to obtain each protein’s hydrophilic and hydrophobic runs. The distribution of the length of runs was plotted for each class of proteins.

## Biophysical Methods

### Protein Expression and Purification

The ClyA plasmid, based on the pET11a backbone and containing a Q56C mutation that retains wild-type lytic activity, was generously provided by Prof. Benjamin Schuler (University of Zurich, Switzerland). Plasmids encoding wild-type ClyA and its mutants were transformed into *E. coli* BL21 (DE3) cells. Protein overexpression was induced with 0.5 mM isopropyl β-D-1-thiogalactopyranoside (IPTG) in an orbital shaker at 25 °C. After overnight incubation (12–14 hours), cells were harvested and lysed by probe sonication in lysis buffer composed of 50 mM potassium phosphate (pH 7.5), 150 mM NaCl, 5% glycerol, 10 mM imidazole, and 10 mM β-mercaptoethanol (βME).Following lysis, cell debris was removed by centrifugation at 20,000 RPM (Hermle centrifuge), and the resulting clear lysate was loaded onto a Ni-NTA affinity column connected to an FPLC system (GE Healthcare Life Sciences). The column was washed with buffer containing 50 mM potassium phosphate (pH 7.5), 400 mM NaCl, 10 mM imidazole, and 10 mM βME. Protein elution was performed using a gradient of imidazole, with the final elution buffer containing 500 mM imidazole, 50 mM potassium phosphate (pH 7.5), 150 mM NaCl, 20% glycerol, and 10 mM βME. Protein purity was assessed by SDS-PAGE, and concentrations were determined using the Bradford assay. Finally, purified protein samples were aliquoted and stored at –80 °C.

### Fluorescence labeling of ClyA

ClyA was site-specifically labeled at position 56, where a cysteine residue was present, using Cyanine3 (Cy3) or Cyanine5 (Cy5) dyes (GE Healthcare Life Sciences) via maleimide chemistry. To ensure the cysteine thiol group was available for labeling, the protein was first reduced using either 50 mM dithiothreitol (DTT) or a 100-fold molar excess of tris(2-carboxyethyl)phosphine (TCEP). When DTT was used, it was removed prior to labeling using a centrifugal concentrator (Amicon Ultra), whereas TCEP remained in the reaction mixture when employed. Cy3 dye, dissolved in DMSO, was added to the reduced ClyA at a fivefold molar excess, and the reaction was incubated overnight at 4 °C. Excess unbound dye was removed by dialysis at 4 °C, followed by further purification through a Sephadex G-25 column. Labeling efficiency was evaluated by absorbance measurements using a Nanodrop spectrophotometer (Thermo Fisher), typically yielding 85–95% efficiency.

### Preparation of liposomes

An appropriate volume of lipid stock solution (25 mg/mL in chloroform) was transferred to a 10 mL glass bottle, and the organic solvent was evaporated by gently passing dry nitrogen gas. To remove residual solvent, the sample was placed in a desiccator connected to a vacuum pump for several hours. The dried lipid film was then hydrated with 20 mM sodium phosphate buffer (pH 7.4) to achieve a final lipid concentration of 10 mM, followed by overnight incubation at 4 °C to ensure efficient hydration of the phospholipid headgroups. The hydrated lipid film was vortexed for approximately 30 minutes to produce multilamellar vesicles (MLVs), with extended vortexing used when needed for uniform mixing. These MLVs were used for sedimentation assays. For liposome preparation, the MLVs were sonicated using a probe sonicator at 45% amplitude for 30 minutes, centrifuged at 5000 rpm to remove tungsten artifacts, and then filtered through a 0.22 μm filter unit. The resulting small unilamellar vesicles (SUVs) were characterized by dynamic light scattering (DLS), yielding an average diameter of approximately 70 nm(Bandyopadhyay *et al*, 2021; Halder *et al*, 2020; Sannigrahi *et al*, 2019).

### Gel-Assisted Synthesis of GUVs

The PVA-gel-assisted synthesis of giant unilamellar vesicles (GUVs) was carried out as follows: a 5% (w/v) solution of poly(vinyl alcohol) (PVA) was prepared by stirring the polymer in Milli-Q water while heating at 90 °C. Approximately 800–1200 µL of this solution was spread onto a 30 mm cell-culture dish and dried in a hot-air oven at 60 °C for 30 minutes to form a thin gel layer. A chloroform-based lipid mixture (20–25 µL) containing various combinations of POPC, cholesterol (CHOL), and sphingomyelin (SM) was then evenly spread over the dried PVA surface. The lipid film was vacuum-dried for ∼30 minutes to remove residual solvent. For hydration, 1000 µL of 1× PBS was gently added to the dish at room temperature, and the setup was left undisturbed for 1 hour. To compensate for evaporation during this period, an additional 300–500 µL of phosphate buffer (pH 7.4) was added after 60 minutes. GUVs were then carefully harvested using a 1 mL pipette tip (with the end trimmed to widen the opening) and transferred into a clean glass vial(Weinberger *et al*, 2013).

### Suspended lipid bilayer (SuLB) formation

Pore-spanning suspended lipid bilayers (SuLBs) were prepared for visualization using fluorescence microscopy and for sulforhodamine B leakage assays. To enable imaging, the GUV solution was doped with either 25 nM DiD (DiIC18(5); 1,1′-dioctadecyl-3,3,3′,3′-tetramethylindodicarbocyanine, 4-chlorobenzene sulfonate salt) or 5 mol% Rhodamine-DOPE, a liquid-disordered (Ld) phase-specific dye. For leakage experiments, 40 µL of 100 nM sulforhodamine B dye was added to the microwells of a pore-spanning polycarbonate track-etch (PCTE) membrane substrate. GUVs were then introduced and incubated on this substrate to facilitate bilayer formation. To assemble the SuLB platform, PVP-coated, continuous-pore, hydrophilic PCTE membranes were placed onto thoroughly cleaned glass coverslips (24 × 50 × 0.13 mm, VWR International). One milliliter of PBS-hydrated GUV solution (pH 7.5) was pipetted onto the membrane surface and incubated at 37 °C for 30 minutes to promote vesicle rupture and bilayer formation on the hydrophilic PVP-coated surface. Excess lipid material was gently removed by rinsing with sodium phosphate buffer. The successful formation of suspended bilayers was confirmed using atomic force microscopy (AFM), fluorescence microscopy, and total internal reflection fluorescence (TIRF) microscopy(Sannigrahi *et al*., 2022).

### Atomic Force Microscopy

Atomic force microscopy (AFM) imaging of pore-spanning suspended lipid bilayers was performed using a Park NX-10 AFM system in contact mode, employing a cantilever with a spring constant of 0.2 N/m. Initially, AFM images of the blank porous substrate were acquired to serve as a control. These were subsequently compared with images of substrates incubated with GUV solutions composed of binary and ternary lipid mixtures, in order to confirm the successful formation of pore-spanning suspended lipid bilayers.

### ClyA pore formation/Leakage study on suspended lipid bilayer

Sulforhodamine B (SRB) dye was trapped within the microwells of a polycarbonate track-etch (PCTE) membrane, which was then sealed on the top and bottom by a lipid bilayer and a coverslip, respectively. Upon perforation of the suspended lipid bilayer (SuLB) by the pore-forming toxin ClyA, SRB diffused through the pores formed by the assembly and insertion of ClyA oligomers. To prepare the SuLB, a 1 µm PCTE membrane (Avanti, 610010) was placed onto a cleaned coverslip, ensuring the glossy side faced upward for inverted microscopy. A solution of SRB (100 nM, 30 µL; Sigma-Aldrich, S9012) was spread onto the membrane and dried for 30 minutes at 37 °C. Subsequently, 50 µL of GUV suspension was applied onto the coverslip, followed by 5 µL of 0.3 M CaCl₂ evenly distributed over the membrane. The membrane-coverslip assembly was incubated undisturbed at 37 °C for 30 minutes, then gently washed with 100 µL of 1× PBS by spreading the buffer and absorbing excess liquid with tissue paper. For the dye-leakage assay, the coverslip was first imaged under a 100× objective to establish baseline fluorescence. ClyA was then added, and the sample was imaged continuously for up to 60 minutes with a fixed field of view. Imaging was performed on an Olympus IX81 microscope equipped with an Andor EMCCD detector (DU-897U), with excitation provided by a 532 nm Coherent Sapphire laser.

### TIRF Microscope Setup and Analysis

A customized imaging setup built around the Olympus IX81 microscope was used to image the PCTE SULB wells containing SRB dye (100nM). A 532nm laser (Sapphire; Coherent) was used to excite the SRB molecules. A combination of 25.4- and 300-mm biconvex lenses (Thor Laboratories) was used to expand the laser beam before a lens of focal length 150 mm was used to focus the (10 mW) laser beam on the back focal plane (BFP) of the objective (UAPON 100X OTIRF; Olympus). The laser spot at the BFP was translated away from the optical axis to achieve inclined illumination to reduce background. Fluorescence emission from an 80 μm × 80 μm area was collected by the objective and passed through a dichroic mirror (FF545/650-Di01-25 × 36; Semrock) and a long-pass filter (BLP02-561R-23.3-D; Semrock) before detection on an electron multiplying charge-coupled device (Andor ixon Ultra 897). Shutter (LS6; Vincent Associates) was used to control the laser illumination. The final imaged area corresponded to 160 nm x 160 nm physical area.

16-bit 512×512 images were captured at different timepoints with 100 ms exposure. The image captured before the addition of protein or the one just after the addition of protein served as a negative control for all the analyses. Well segmentation was achieved as follows. All the images in the timeseries were registered with the negative control for stage-drift correction using either the SIFT algorithm or the spatial intensity cross-correlation-based registration algorithm, whichever gave the least error(Huang *et al*, 2008).This was followed by generating a binary image from the negative control. Circular Hough transform(Meng *et al*, 2018) was applied to this binary image for locating the wells on the negative control. The size and location of the wells extracted from this binary image were utilized to generate a mask. This mask was imposed on all the images in the time series. The average intensity of the well was calculated after isolating the single wells (based on the size and perimeter-to-diameter ratio). Unresolvable wells were omitted from the analysis. The resultant intensity histograms were plotted with time as a heatmap, which demonstrated the pore-forming activity by ClyA under different conditions. We collected the data in duplicates to ensure robust statistics.

### Fluorescence imaging of different domains in SuLB

Giant unilamellar vesicles (GUVs) were synthesized using a ternary lipid mixture doped with 5 mol% Rhodamine-DOPE, and a suspended lipid bilayer (SuLB) platform was formed via GUV rupture to visualize phase separation in the bilayer. Fluorescence microscopy revealed two distinct domains: dimmer regions corresponding to the liquid-ordered (Lo) phase and brighter regions corresponding to the liquid-disordered (Ld) phase. The phase-separated SuLBs were treated with 100 nM Cy5-labeled ClyA to investigate colocalization and morphological changes in the lipid bilayer over various incubation time points. Image analysis was performed using Fiji (ImageJ) software.

### Isolation of DDM induced different preassembled states of ClyA

Three distinct populations—referred to as the I state, InP state, and cP state—corresponding to different stages of DDM-induced preassembled ClyA intermediate states were isolated as follows. A 100 µL reaction was prepared in a 1.5 mL microcentrifuge tube by mixing ClyA (1 µM) with 0.1% (w/v) DDM in 1× PBS. After 60 seconds, a 25 µL aliquot of the reaction mixture was transferred into a separate 1.5 mL tube containing 475 µL of 1× PBS and mixed thoroughly. Subsequent 25 µL aliquots were collected at 15, 40, and 60 minutes post-reaction initiation and each transferred into individual tubes containing 475 µL of 1× PBS. All collected samples were immediately stored at –80 °C until further use.

### Circular Dichroism Measurement

Far-UV CD spectra of wild-type (WT) ClyA and its mutants were recorded in the absence and presence of binary and ternary liposomes using a JASCO J-720 spectropolarimeter (Japan Spectroscopic Ltd, Ishikawa-cho, Hachioji-shi, Japan). Measurements were performed between 200 and 260 nm using a 1 mm path-length cuvette at a protein concentration of 20 μM. The scan speed was set to 50 nm/min with a response time of 2 s and a bandwidth of 1 nm. Three scans were recorded in continuous mode and averaged. A typical protein-to-lipid molar ratio of 1:50 was maintained. CD spectra of individual DDM-induced ClyA populations (I, InP, and cP states) were collected using a JASCO J-715 spectropolarimeter with a 0.1 cm path-length cuvette. Ellipticity was measured from 200 to 260 nm in 0.5 nm increments using a 2 nm bandwidth. Temporal changes in helicity upon DDM treatment were monitored at 222 nm for up to 60 minutes.

### TEM imaging

ClyA samples, both in the presence and absence of binary or ternary liposomes, as well as DDM-induced preassembled ClyA states, were visualized using negative staining electron microscopy (EM) to assess particle homogeneity and distribution. Samples were prepared using standard negative staining protocols. Carbon-coated copper EM grids (300 mesh; Ted Pella) were glow-discharged for 30 seconds at 20 mA using a GloQube glow discharge system (Quorum). A 3.5 μL aliquot of sample (0.1 mg/mL) was applied to each glow-discharged grid and incubated for 30 seconds before excess liquid was blotted off. Grids were then stained with 1% (w/v) uranyl acetate (98% purity; ACS Reagent, Polysciences Inc., Warrington, PA, USA) for 20 seconds and allowed to air-dry. Imaging was performed at room temperature on a Tecnai T12 electron microscope (FEI) equipped with a LaB₆ filament, operating at 120 kV. Images were acquired using a side-mounted Olympus VELITA CCD camera (2000 × 2000 pixels) at a magnification of 220,000×, corresponding to a pixel size of 2.54 Å.

### Preparation of Calcein Entrapped liposomes

To prepare dye-entrapped vesicles, the required amount of lipids dissolved in chloroform was added to a cleaned glass vial. The solvent was gently evaporated under a stream of nitrogen to form a uniform lipid film on the vial walls. After complete drying, 1 mL of 1× PBS was added to hydrate the lipid film. Calcein AM (Merck, 17783), pre-solubilized in 1 M KOH, was added to the lipid suspension to achieve a final dye concentration of 70 mM. The resulting mixture was probe-sonicated for 30 minutes using a Branson Digital Sonifier-450 (Branson, USA) to form dye-entrapped vesicles. To remove unencapsulated dye, the vesicle suspension was subjected to dialysis for 24 hours at 4 °C in 1× PBS using SnakeSkin™ dialysis tubing (3.5 kDa MWCO, 16 mm; Thermo Fisher Scientific, 88242) under gentle stirring.

### Assay for Permeabilization of Lipid Vesicles

The ability of ClyA, different mutants and DDM induced ClyA preassembled states to promote the release of calcein from entrapped SUVs composed of different lipids was checked by monitoring the increase in fluorescence intensity of calcein. Calcein-loaded liposomes (70 mM concentration) were separated from non-encapsulated (free) calcein by gel filtration on a Sephadex G-75 column (Sigma) using an elution buffer of 10 mM MOPS, 150 mM NaCl and 5 mM EDTA (pH 7.4), and lipid concentrations were estimated by complexation with ammonium ferro-thiocyanate Fluorescence intensity was measured at room temperature (25°C) using a PTI spectro-fluorometer. A path length of 1cm was used for each measurement. The excitation wavelength was 490 nm and emission was set at 520 nm. Excitation and emission slits with a nominal bandpass of 3 and 5 nm were used, respectively. The high concentration (70 mM) of the entrapped calcein led to self-quenching of its fluorescence resulting in low fluorescence intensity of the vesicles (I_B_). Release of calcein caused by the addition of protein samples led to dequenching of the dye into the medium, which could therefore be monitored by an enhancement of fluorescence intensity (I_F_). This enhancement of fluorescence is a measure of the extent of vesicle permeabilization. The experiments were normalized relative to the total fluorescence intensity (I_T_) corresponding to the total release of calcein after complete disruption of all the vesicles by addition of Triton X-100 (2% v/v). The percentage of calcein release in the presence of proteins was calculated using the equation:

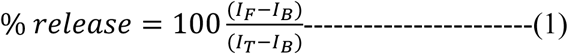

where, I_B_ is the background (self-quenched) intensity of calcein encapsulated in vesicles, I_F_ represents the enhanced fluorescence intensity resulting from the dilution of dye in the medium caused by the protein induced release of entrapped calcein. I_T_ is the total fluorescence intensity after complete permeabilization is achieved upon addition of Triton X-100. Typical SUVs concentration was taken 100 μM and 0.5μM concentration of each protein sample was used.

For temperature-dependent lysis measurements using wild-type (WT) ClyA, calcein-loaded vesicles composed of various lipid components were first assessed for their baseline (quenched) fluorescence intensity. The vesicles were then incubated with WT ClyA at a final concentration of 0.5 µM for 2 minutes at different temperatures. Following incubation, the fluorescence intensity of dequenched calcein was measured. To determine total dye release, 2% (v/v) Triton X-100 was added to each sample, and fluorescence intensity corresponding to complete vesicle disruption was recorded. The percentage of calcein leakage at each temperature was calculated using Equation 1, as described above.

### Erythrocyte Lysis Assay

For kinetic measurements of erythrocyte lysis, ClyA variants were incubated with 200 μL of a 1% (v/v) suspension of rabbit erythrocytes in phosphate-buffered saline (PBS) at a final protein concentration of 0.5 µM. Turbidity reduction, indicative of cell lysis, was monitored by measuring optical density at 620 nm over time using a microplate reader (Tecan) with intermittent orbital shaking at 37 °C. The resulting lysis kinetics data were fitted to a Boltzmann sigmoidal function to extract the half-life of lysis (t₁/₂)(Sathyanarayana *et al*., 2018).

### Probing Secondary Conformations using ATR FTIR

Fourier-Transform Infrared (FTIR) spectroscopy was used to monitor the secondary structure of protein–detergent intermediates and lipid-membrane-induced conformational changes. Measurements were performed using a Perkin-Elmer Spectrum 3 Dual Range MIR/NIR Spectrometer (L1280138) operated in attenuated total reflectance (ATR) mode. Spectra were acquired over the wavenumber range of 1600–1700 cm⁻¹, with a scan rate of 32 scans/sec and an integration interval of 0.5 seconds. At designated time points, 10 μL of the reaction mixture (containing either protein–detergent or protein–lipid complexes) was applied directly to the ATR crystal to assess temporal structural changes. Spectral data were baseline-corrected and subjected to appropriate curve fitting (Sannigrahi *et al*., 2021) for peak analysis.

### Laurdan Generalized polarization (GP) Measurement in model membrane

Laurdan fluorescence measurements were performed using a PTI fluorescence spectrophotometer. Liposomes of varying lipid compositions were labeled by incubating them with Laurdan (dissolved in DMSO) at a final concentration of 5 µM for 20 minutes at room temperature(Sannigrahi *et al*., 2024). The Laurdan-labeled liposomes were then excited at 340 nm, and emission spectra were recorded from 400 to 550 nm at a scan speed of 60 nm/min. Membrane order was quantified using the generalized polarization (GP) parameter, defined as:

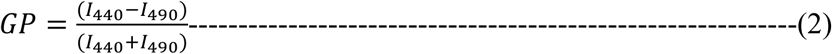

were obtained. Here *I*_440_ and *I*_490_ stand for the intensities at the wavelength at 440 nm and 490 nm respectively.

### Fluorescence recovery after photobleaching (FRAP) of lipid bilayers

We prepared our SULB platform using binary and ternary lipid compositions doped with lipophilic tracer DiI C-18. The sample was imaged on a Leica STED SP5 confocal microscope with an Argon laser and the emission was separated and collected between 550 and 580 nm by adjusting the acousto-optical beam splitter. Fluidity of the membrane was assessed by performing FRAP analysis. A circular area (radius, w = 1 μm) was bleached using high intensity illumination. The recovery kinetics of the lipophilic tracer molecule in this area was monitored. The recovery kinetics was fitted to an exponential of the form:

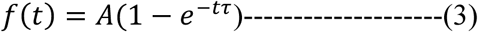

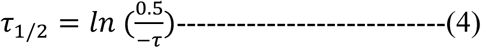

Diffusion coefficient (D) was estimated by the relation below.

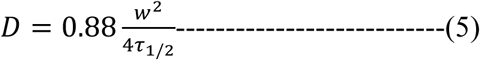

Where,Ꚍ1/2 and w denote the half time of recovery and radius of uniform bleach laser respectively.

### Sedimentation Assay to measure binding

We assessed protein–lipid binding using multilamellar vesicles (MLVs) by incubating the purified unlabeled proteins (WT ClyA and the mutants) with pre-formed MLVs labeled with 0.5 mol% of DiI C-18 (1 mM lipid concentration) in binding buffer (PBS, pH 7.4) at room temperature for 30 minutes. Following incubation, we centrifuged the samples at 23000 × g for 45 minutes at 4°C to sediment the MLVs. We then carefully separated the supernatant and resuspended the pellet in an equal volume of SDS loading buffer. Both fractions were analyzed by SDS-PAGE and visualized using Coomassie staining. Protein binding was indicated by the presence of protein in the pellet fraction compared to the protein-only control, and band intensities were quantified by densitometry. The fraction of bound protein was calculated using the equation:

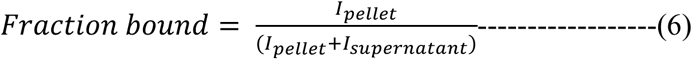

Where, *I_pellet_* and *I_supernatant_* are the band intensities of the pellet and supernatant, respectively. The fluorescence intensities of the isolated supernatant fraction and pellet fraction were measured to compare the similar pelleting potential of MLVs irrespective of lipid compositions.

For time-dependent binding studies, a similar protocol was followed using Cy5-labeled ClyA (Q56C) and MLVs composed of different lipid components. The labeled protein was incubated with MLVs at a final lipid concentration of 1 mM for varying time intervals. After incubation, samples were subjected to high-speed centrifugation at 23,000 × g for 45 minutes to pellet the lipid-bound protein. The unbound protein remaining in the supernatant was quantified by measuring the fluorescence intensity of Cy5-labeled ClyA. The decrease in supernatant fluorescence over time was used to assess the extent and kinetics of protein binding to the lipid vesicles.

### Cell Biology methods

#### Cell Culture and treatments

RAW 264.7 macrophages were maintained in DMEM supplemented with 10% FBS and 1% penicillin–streptomycin in a humidified incubator at 37 °C with 5% CO₂. On Day 0, approximately 2 × 10⁴ cells were seeded into 96-well plates and allowed to adhere overnight under standard culture conditions. On Day 1 of treatment, cells were pre-incubated with either sphingomyelinase (SMase; final concentration 50 mU/mL) for 30 min, sphingomyelin (SM) liposomes (final concentration 20 µM) for 6 h, methyl-β-cyclodextrin (MβCD; final concentration 5 mM) for 30 min, or cholesterol liposomes (final concentration 20 µM) for 6 h. Following these pre-treatments, ClyA was added at a final concentration of 100 nM for the indicated durations before subsequent analysis.

Following the treatments, cells were gently washed with PBS and incubated with propidium iodide (PI; final concentration 3 µg/mL) at the designated time points to assess membrane integrity and cell viability. Fluorescence imaging was performed at each time point using a Thermo Fisher EVOS M7000 Cell Imaging System under identical acquisition settings, ensuring comparability across conditions. The acquired images were analyzed using ImageJ software to quantify PI-positive cells, enabling evaluation of the effects of SMase or SM pre-treatment on ClyA-induced cytotoxicity.

### Confocal Microscopy to detect membrane lipid raft

RAW 264.7 macrophages (1 × 10^4^ cells per well) were seeded into μ-Slide 8 Well chambered slides (ibidi, Cat. No. 80826) and cultured overnight in DMEM supplemented with 10% FBS at 37 °C. Following incubation, cells were treated with either Cy3-labelled or unlabelled ClyA (50 nM) for the indicated time periods. Subsequently, cells were washed three times with 1× PBS and stained for lipid rafts using cholera toxin subunit B conjugated to Alexa Fluor 647 (CTxB-AF647; Invitrogen, C34778) or for lipid droplets using LipidSpot-610 (Biotium), following the manufacturer’s instructions. Briefly, adherent cells were incubated with CTxB-AF647 (4 µg/mL) for 30 min in dark, washed three times with 1× PBS, fixed with 4% paraformaldehyde for 15 min at room temperature, and stained with DAPI. Confocal images were acquired on a Leica TCS SP8 FALCON microscope using a 63× objective and analyzed using Fiji ImageJ.

### Statistics and reproducibility

All results were confirmed in at least three independent experiments performed on different days with different batches of samples. The results were expressed as the Mean ± SD derived from multiple data points. Statistical significance in the observed differences was determined through analysis of variance (ANOVA) and unpaired t-test utilizing GraphPad prism (9.0) software. A significance level of p < 0.05 was considered indicative of meaningful differences.

## Data availability

No data has been deposited in any public database.

## Supporting information

Additional text of SuLB platform characterization, Tables S1, S2, and Figures S1 to S22 have been provided in the supplementary file.

## Acknowledgement

AS acknowledges SERB (PDF/2020/000678), Govt of India for providing National Postdoctoral Fellowship. RR acknowledges financial support from the Human Frontier Science Program (HFSP/RGP0047/2022). We acknowledge IISc Central AFM, EM and bio-imaging facilities for their support for instrumentation.

## Supplementary Material SM1

### Characterization of High-Throughput Pore-Spanning Suspended Lipid Bilayer (SuLB) Platform

To investigate membrane pore formation by ClyA, we employed SuLB systems incorporating both binary and ternary lipid compositions. Characterization of these SuLB systems (**Figure S4A**) was performed using atomic force microscopy (AFM), which confirmed successful bilayer formation spanning the porous PCTE (polycarbonate track-etched) substrate (Sannigrahi *et al*, 2022). AFM measurements revealed a significant reduction in depth—from 126 ± 18 nm for bare microwells to 20–70 nm after membrane coverage—indicating the formation of suspended bilayers (**Figure S4B i–iii**). The reduction in axial depth (∼50–100 nm) and contrast in AFM images further supported the formation of the bilayer via ruptured GUVs spreading across the microwell array (**Figure S4B i–iii**). AFM also revealed membrane deflection consistent with the increased bending stiffness of suspended regions compared to the supported ones, reflecting the higher spring constant of the suspended membranes. We define membrane curvature as the ratio of maximum depression at the membrane center to the pore radius. Among the tested compositions, ternary membranes exhibited the lowest curvature (0.047 ± 0.014), followed by binary (0.068 ± 0.017), and POPC-only membranes (0.10 ± 0.01) (**Figure S4C**), indicating relatively uniform bending across all membrane types. The binary membranes, primarily in a liquid-disordered (Ld) phase at room temperature, exhibited increased order upon the addition of sphingomyelin, forming more liquid-ordered (Lo) domains. This ordering leads to increased membrane stiffness and reduced bending. Furthermore, the ternary SuLBs showed greater surface roughness, suggesting the presence of Lo/Ld phase segregation (**Figure S4B**). We also evaluated SuLB formation on PCTE substrates with larger (3 µm) pore diameters and found lipid-dependent differences in membrane bending. Ternary SuLBs showed reduced deflection compared to binary systems (**Figure S4D**), reinforcing the role of lipid composition in determining mechanical properties. The increased roughness in ternary membranes (**Figure S4D**) again pointed to phase separation. Fluorescence imaging with rhodamine-DOPE, a probe for the Ld phase, further supported phase separation. Binary membranes showed uniform fluorescence, while ternary membranes exhibited both bright (Ld) and dim (Lo) regions, confirming coexistence of distinct domains (**Figure S4E, F**). Notably, Lo domains were predominantly located outside the pore cavities, with only occasional small Lo domains appearing in the suspended regions—an observation consistent with previous studies(Schütte *et al*, 2017). Together, these results demonstrate that the SuLB platform effectively replicates lipid composition–dependent effects on curvature and stiffness in suspended bilayers. To assess bilayer fluidity, both binary and ternary SuLB membranes were labeled with DiI C18 and subjected to fluorescence recovery after photobleaching (FRAP). The similar recovery rates observed suggest that both membrane types maintain high lateral mobility and remain sufficiently fluid (**Figure S4G**). To probe ClyA-induced pore formation, SuLBs with binary and ternary compositions were assembled on PCTE membranes with 1 µm pores. Microwells were preloaded with hydrophilic sulforhodamine B (SRB) dye and sealed with lipid bilayers formed by GUV rupture (**Figure S4H**). Pore formation was assessed by tracking SRB release via a reduction in fluorescence intensity over time and analyzing the frequency of dye-retaining wells. Time-lapse TIRF microscopy enabled kinetic monitoring of SRB leakage in individual SuLB areas.

---

**Supplementary Table S1:** Complete simulation details for ClyA simulations in presence of DDM.

| System | No. of DDM molecules | No. of protein residues | No. of water molecules | Simulation time (ns) |
| --- | --- | --- | --- | --- |
| Monomer with 50 DDM | 50 | 303 | 67161 | 500 |
| Monomer with 100 DDM | 100 | 303 | 67161 | 180 |
| β-tongue with micelle | 100 | 38 | 10992 | 1000 |
| β-tongue in solution | 0 | 38 | 5020 | 1000 |

**Supplementary Table S2.** Detailed information of all the Lipid-MD Simulation systems studied.

| Protein | Lipid | No. of lipid molecules |  |  | No. of water | Simulation time (ns) |
| --- | --- | --- | --- | --- | --- | --- |
|  |  | POPC | CHOL | SM |  |  |
| Pore | Binary | 341 | 143 | - | 77532 | 300 |
|  | Ternary | 176 | 174 | 172 | 76609 | 300 |
| β-tongue | Binary | 140 | 60 | - | 8471 | 1000 |
|  | Ternary | 66 | 66 | 66 | 7433 | 1000 |
| Unfolded β-tongue | Binary | 140 | 60 | - | 5298 | 1000 |
|  | Ternary | 66 | 66 | 66 | 4578 | 1000 |

## Supplementary Methods and Materials

Supplementary Methods and Materials:
| Reagent/resource | Source/Reference | Identifier/catalog number |
| --- | --- | --- |
| ClyA Q56C | Benke et al, 2015 | Gift from Benjamin Schuler (University of Zurich) |
| K172A,K175A,K206A | DX/DT Enzymes | N/A |
| Raw 264.7 | RAW 264.7 | Cat.#TIB-71 |
| Vero cell | Vero E6 | Cat.#CRL-1586 |
| BL21(DE3) pLs S Escherichia coli competent cells | Invitrogen | Cat.#C606003 |
| Coomassie brilliant Blue R250 staining Solution | Bio-Rad | Cat.#161-0436 |
| Dulbecco's phosphate buffered solution | Corning | Cat.#D0632-25G |
| Dulbecco's modified Eagle's medium | DMEM, High Glucose, Gibco | Cat.# 11995065 |
| Quik Change II XL Site-Directed Mutagenesis Kit | Agilent | Cat.#200522 |
| Penicillin and streptomycin sulfate | Corning | Cat. #30-002-C1 |
| Phenylmethanesulfonyl fluoride (PMSF) | Sigma-Aldrich | Cat.#P7626-25G |
| Cy3 Maleimide | GE Healthcare life sciences |  |
| Cy5 Maleimide | GE Healthcare life sciences |  |
| N-palmitoyl-D-erythro-sphingosylphosphorylcholine | Avanti Polar Lipids | Cat.#860584 |
| 1-palmitoyl-2-oleoyl-glycero-3-phosphocholine | Avanti Polar Lipids | Cat.#850457 |
| Cholesterol | Sigma-Aldrich | Cat.# C8667 |
| DAPI | SRL | Cat.#28488 |
| Paraformaldehyde | Himedia | Cat.# TC703 |
| Cholera Toxin Subunit B (Recombinant), Alexa Fluor™ 647 Conjugate | Invitrogen | Cat.#C34778 |
| LipidSpot™610 | Biotium | Cat.#70069 |
| Laurdan | Invitrogen | Cat.# D250 |
| 1,2-dioleoyl-sn-glycero-3-phosphoethanolamine-N-(lissaminerhodamine B sulfonyl) (ammonium salt) | Avanti Polar Lipids | Cat.#810150 |
| Sphingomyelinase from *Staphylococcus aureus* | Millipore Sigma | Cat.# S8633 |
| Urea | Millipore Sigma | Cat.# U5128 |
| Poly(vinyl alcohol) | Millipore Sigma | Cat.# P1763 |
| Sulforhodamine B | Millipore Sigma | Cat.#230162 |
| Dil Stain (1,1'-Dioctadecyl-3,3,3',3'-Tetramethylindocarbocyanine Perchlorate ('DiI'; DiIC<sub>18</sub>(3))) | Invitrogen | Cat.#D3911 |
| n-dodecyl-β-D-maltoside (DDM) | Avanti Polar lipids | Cat.#850520P |
| Polycarbonate (PCTE) Membrane Filters, 1.0 Micron, 25mm, 100/Pk | SterliTech | Cat.#PCT1025100 |
| Polycarbonate (PCTE) Membrane Filters, 3.0 Micron, 25mm, 100/Pk | SterliTech | Cat.#PCT3025100 |
| GraphPadPrism9.0 |  | N/A |
| Adobe Photoshop & Adobe Illustrator |  | N/A |
| ImageJ | https://imagej.nih.gov/ij/ | N/A |
| Endnote |  | N/A |
| BioRender | Scientific Image and Illustration Software\| | N/A |
| Olympus VELITA charge-coupled device camera | Olympus | N/A |

---

**Supplementary Figure S1.**
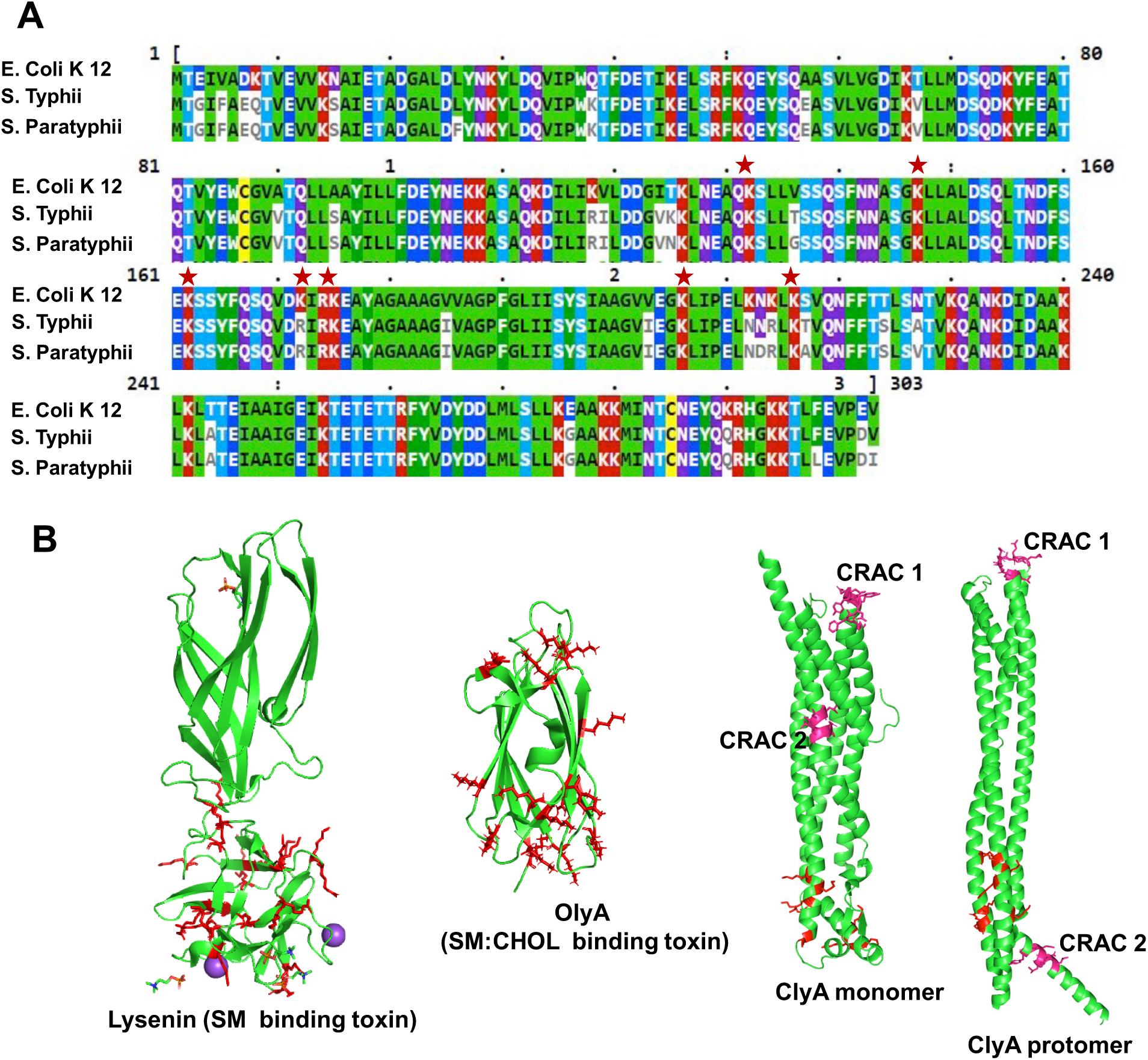
(A) Sequence similarity analysis of ClyA from different organisms was performed using Clustal Omega. Asterisks indicate conserved arginine (R) and lysine (K) residues near the β-tongue region. (B) Structures of sphingomyelin-binding protein Lysenin (PDB: 3ZX7), sphingomyelin–cholesterol-binding protein OlyA (PDB: 6MYI), and the monomeric and protomeric forms of ClyA (PDB: 1QOY, 6MRT). Red-highlighted residues indicate lysine and arginine residues, while pink-colored residues represent the CRAC (cholesterol recognition/interaction amino acid consensus) domain.

**Supplementary Figure S2.**
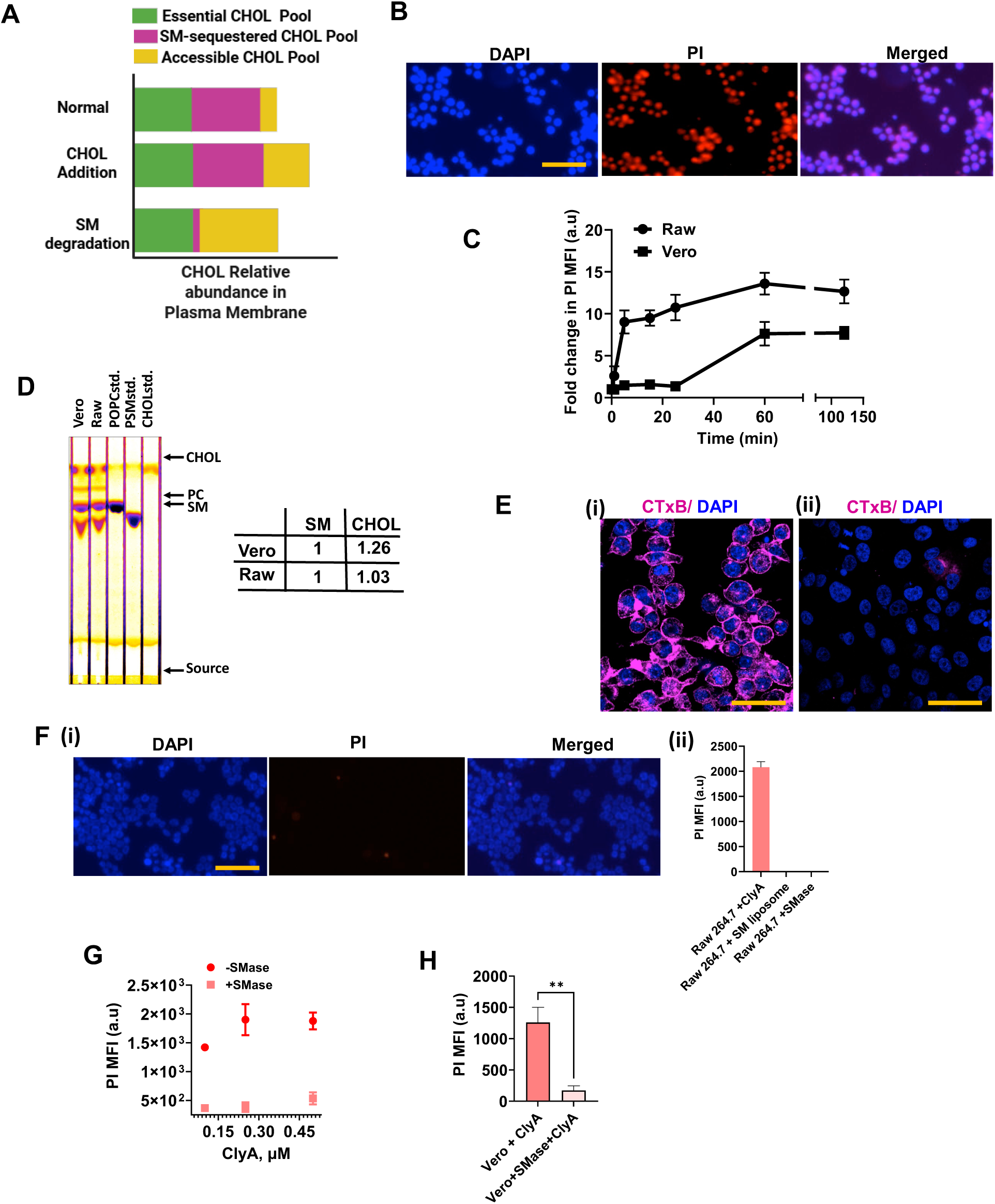
(A) Schematic representation of cholesterol distribution within the mammalian plasma membrane. The chemical accessibility of cholesterol is modulated by its association with SM, creating distinct functional pools. (B) Pore formation in RAW 264.7 cells treated with WT ClyA (100 nM) was assessed by monitoring propidium iodide (PI) uptake. Increased intracellular PI fluorescence indicates significant pore formation induced by WT ClyA. Scale bar is 100 μm. (C) Temporal analysis of PI intensity in RAW 264.7 and Vero cells following exposure to 100 nM WT ClyA. The data show that WT ClyA induces pore formation more rapidly in RAW 264.7 cells than in Vero cells. Data were represented as mean ± SD (n = 3). n stands for number of experimental repeats. (D) Thin-layer chromatography (TLC) analysis of total lipids extracted from 6 × 10⁶ RAW and Vero cells. The TLC image shows the relative levels of sphingomyelin (SM), cholesterol (CHOL), and phosphatidylcholine (PC). The accompanying table summarizes the SM and CHOL ratios estimated from the TLC. (E) Lipid raft detection in (i) RAW 264.7 and (ii) Vero cells using Alexa 647-labeled cholera toxin subunit B (CTxB, magenta). Nuclei were counterstained with DAPI (blue). Scale bar: 50 μm. (F) (i) PI fluorescence in RAW cells treated with sphingomyelinase (SMase, 50 mU/mL for 30 min), showing minimal pore formation. Scale bar: 100 μm. (ii) Quantification of PI mean fluorescence intensity (MFI) under different treatment conditions in RAW cells. (G) The plot shows PI MFI values in Raw Cells across different doses of WT ClyA under treated and untreated conditions with 50 mU/mL SMase. (H) PI MFI in Vero cells (2×10^4^/well) treated with WT ClyA alone or with sequential SMase and WT ClyA. Data are presented as mean ± SD (n = 3) Statistical significance was determined using unpaired t-tests (GraphPad Prism v9), **P =0.0017.

**Supplementary Figure S3:**
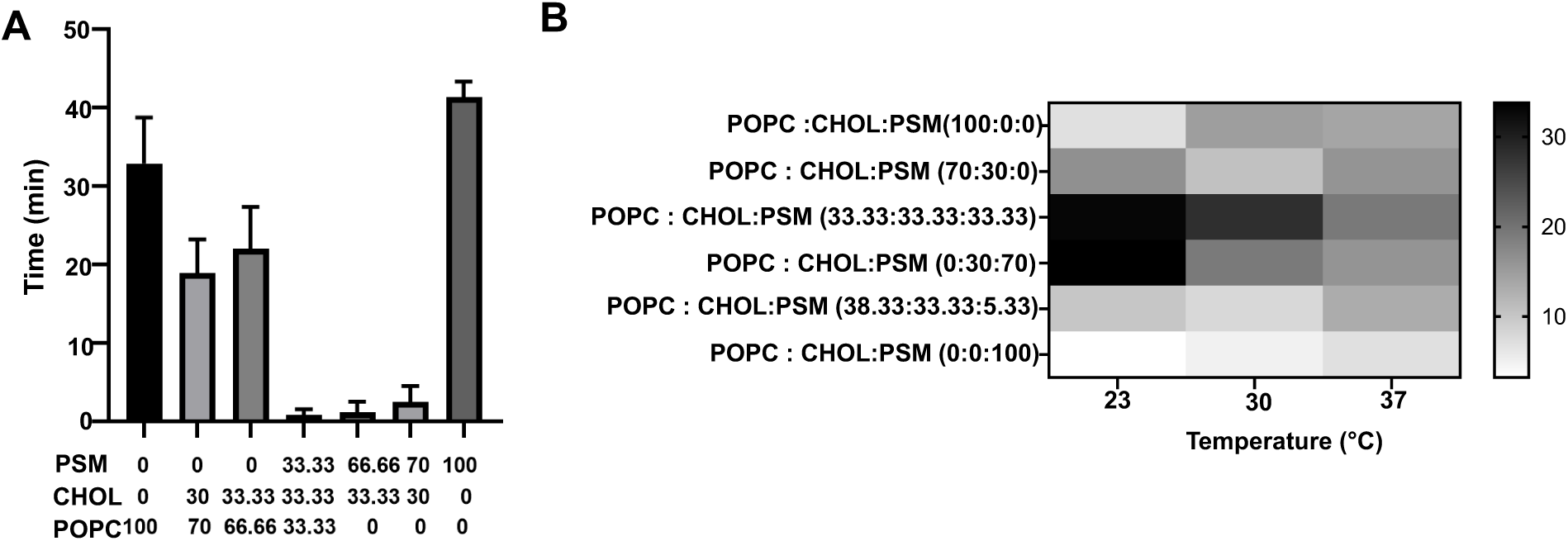
(A) Half-lives of calcein leakage from calcein-entrapped liposomes composed of different lipid components(molar percentages of each liposome composition are mentioned below the bar graph). Liposomes were treated with 500 nM wild-type ClyA to monitor dye leakage kinetics. Data are presented as mean ± SD (n = 3), where n indicates the number of independent experimental repeats. (B) Heatmap displays the extent of calcein dye leakage from liposomes of various compositions at three different temperatures. Liposomes composed of different lipid components were exposed to WT ClyA for 2 minutes at the indicated temperatures, and leakage was measured using fluorescence readouts. Color bar indicates the extent of dye release.

**Supplementary Figure S4.**
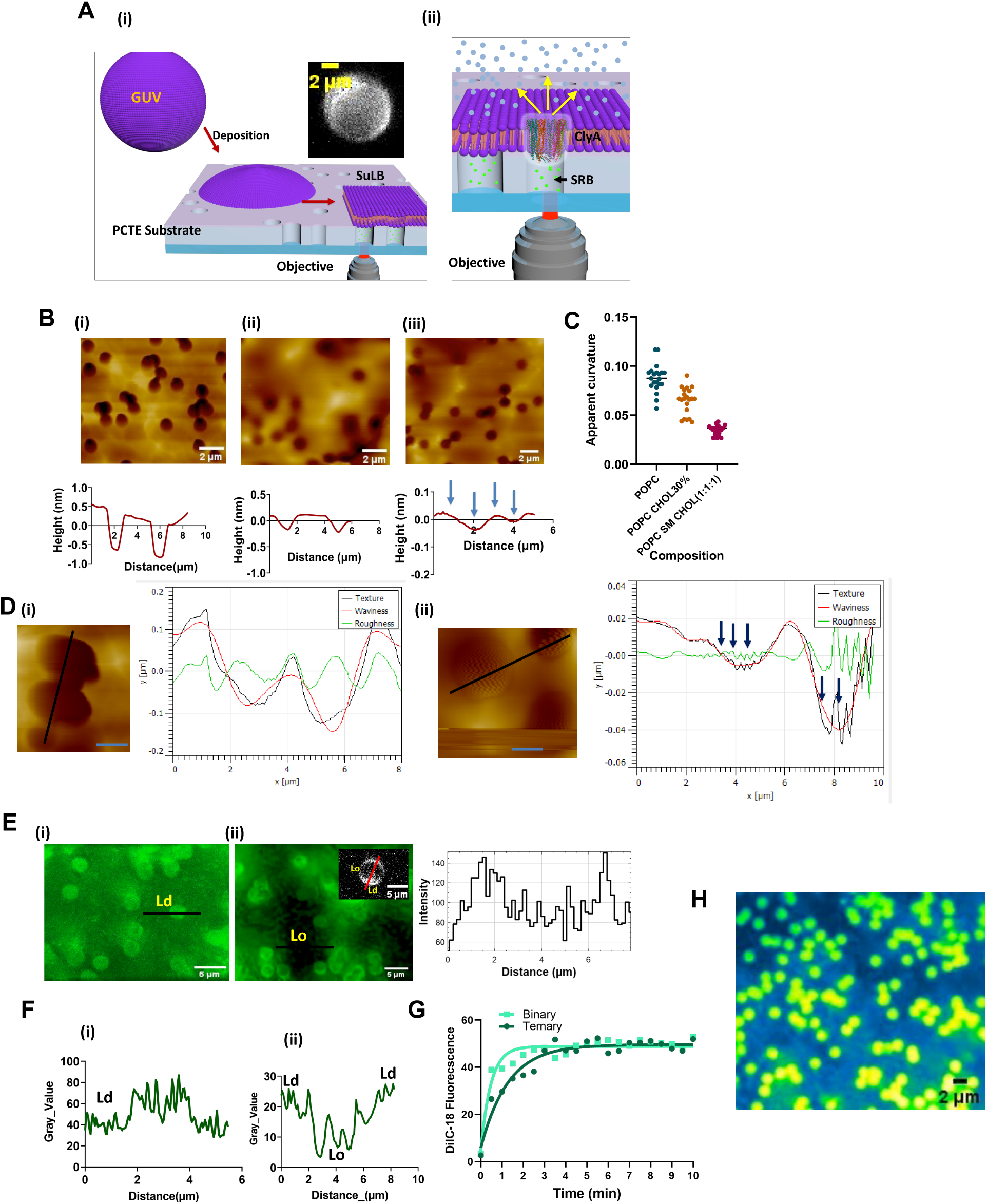
Generation and characterization of the SuLB (Suspended Lipid prepared via the gel-assisted method are ruptured and spread over a PCTE polymeric substrate placed on a glass coverslip. (ii) Schematic showing the formation of ClyA pores in the bilayer of the SuLB system. (B) AFM topographic images of (i) PCTE substrate alone, (ii) SuLB composed of binary lipid components, and (iii) SuLB composed of ternary lipid components. Corresponding AFM height profiles show bilayer depth and microwell cavity size. Arrows indicate surface roughness in the ternary system, attributed to phase-separated lipid domains. PCTE substrates used contained 1μm sized microwell cavities. (C) Plot of apparent curvature in blank, binary, and ternary SuLB systems. (D) AFM micrographs of SuLB platforms composed of (i) binary and (ii) ternary lipid mixtures. Height profiles to the right of each image indicate increased surface roughness and reduced depth in the ternary system compared to the binary system. PCTE substrates used contained 3μm sized microwell cavities. Scale bar is 1μm. (E) Fluorescence micrographs of (i) binary and (ii) ternary SuLB membranes labeled with Rhodamine-DOPE, a marker for liquid-disordered (Ld) phase. Green regions represent Ld domains; darker (black) regions in (ii) indicate liquid-ordered (Lo) phases. The intensity plot on the right shows a line profile across the red solid line in the phase-separated GUV in (ii). (F) Fluorescence line scan profiles of (i) binary and (ii) ternary SuLB systems. (G) Fluorescence recovery after photobleaching (FRAP) of binary and ternary SuLB membranes labeled with DiI C-18. The plot compares FRAP recovery curves, indicating similar membrane fluidity in both systems. (H) Representative TIRF image of the SuLB platform entrapping sulforhodamine B (SRB). The SRB dye is confined within the microwell cavity beneath the suspended bilayer. Microwell size is 1 μm.

**Supplementary Figure S5:**
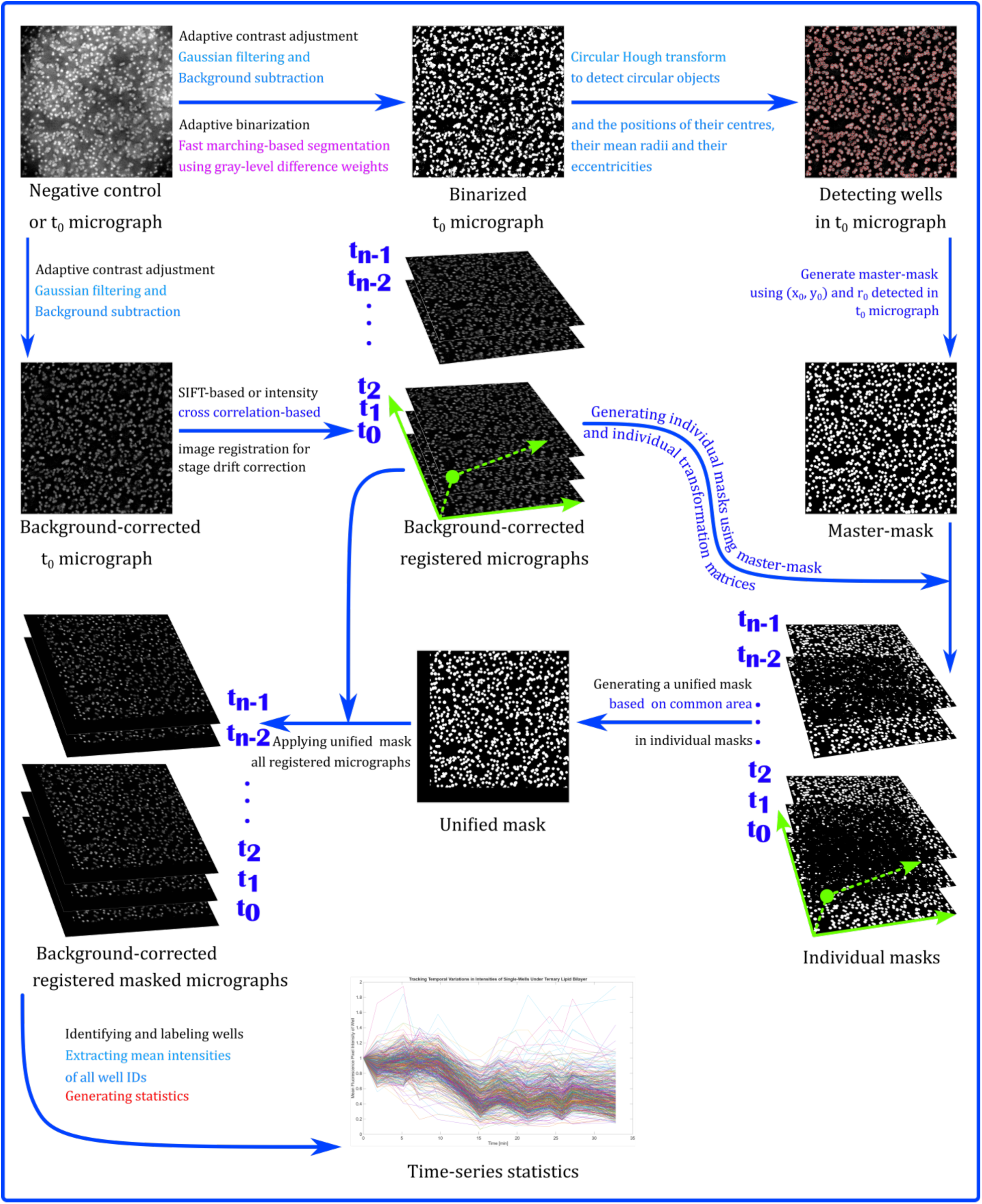
Workflow of image processing of the SuLB systems to track pore

**Supplementary Figure S6:**
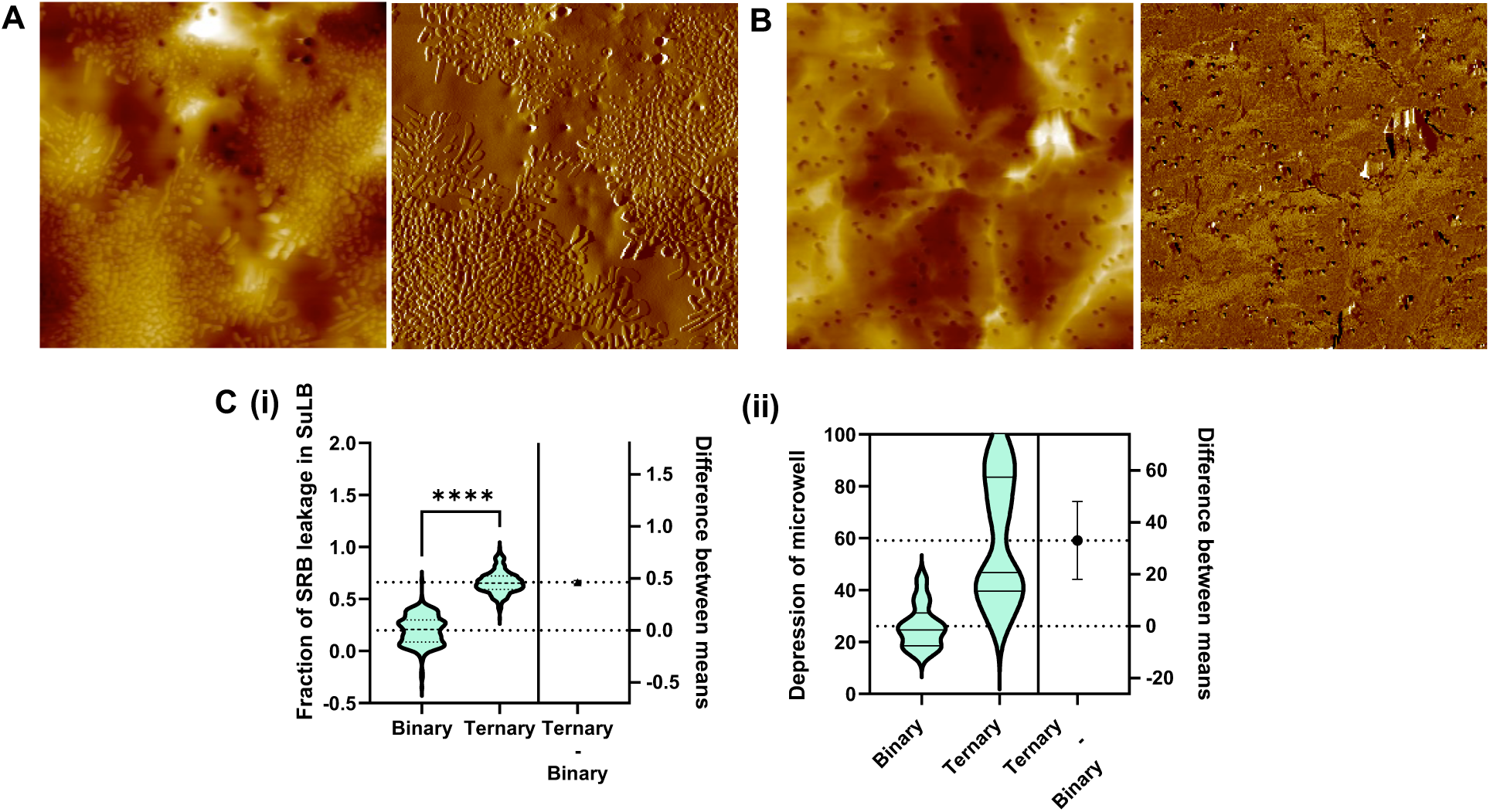
AFM images of SuLB to visualize the oligomerization status of WT ClyA. (A) AFM tomographic image (left) and the phase image (right) of ternary SuLB on PCTE substrate. A significant number of WT ClyA oligomers were found on the suspended bilayer. Here ternary SuLB platform was treated with WT ClyA (100 nM) and incubated for 10 minutes and then AFM imaging was performed. Microwell diameter was 1 μm. (B) AFM tomographic image (left) and the phase image (right) of binary SuLB on PCTE substrate. No oligomer was found on the suspeneded bilayer. Here binary SuLB platform was treated with WT ClyA (100 nM) and incubated for 10 minutes and then AFM imaging was performed. Microwell diameter was 1 μm. (C) (i) plot shows the fraction of SRB leakage from binary and ternary SuLB after 100 nM WT ClyA exposure for 10 mins. (ii) Plot shows the depression of microwell in binary and ternary SuLB after WT ClyA exposure for 10 mins. The microwell depressions were calculated from the AFM images.

**Supplementary Figure S7.**
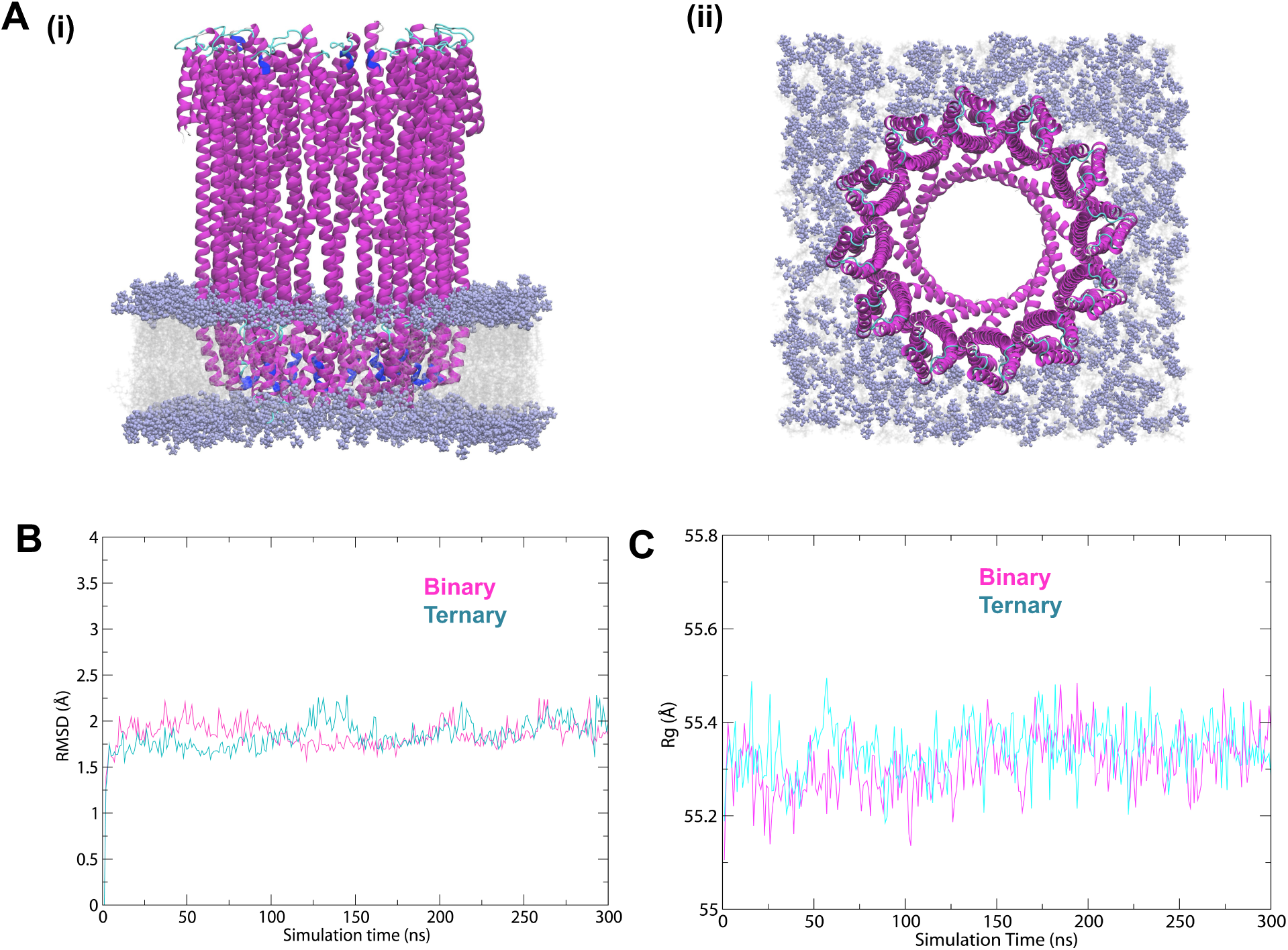
(A) ClyA dodecameric pore embedded in a POPC:CHOL (70:30) lipid bilayer after 300 ns NPT molecular dynamics simulation. (i) Side view, (ii) Top view. (B) Root mean square deviation (RMSD) of backbone atoms of the ClyA pore in binary (POPC:CHOL 70:30) and ternary (POPC:CHOL:SM, 33.3:33.3:33.3) over the course of the simulation. (C) Radius of gyration (Rg) of backbone atoms of the ClyA pore Versus simulation time.

**Supplementary Figure S8.**
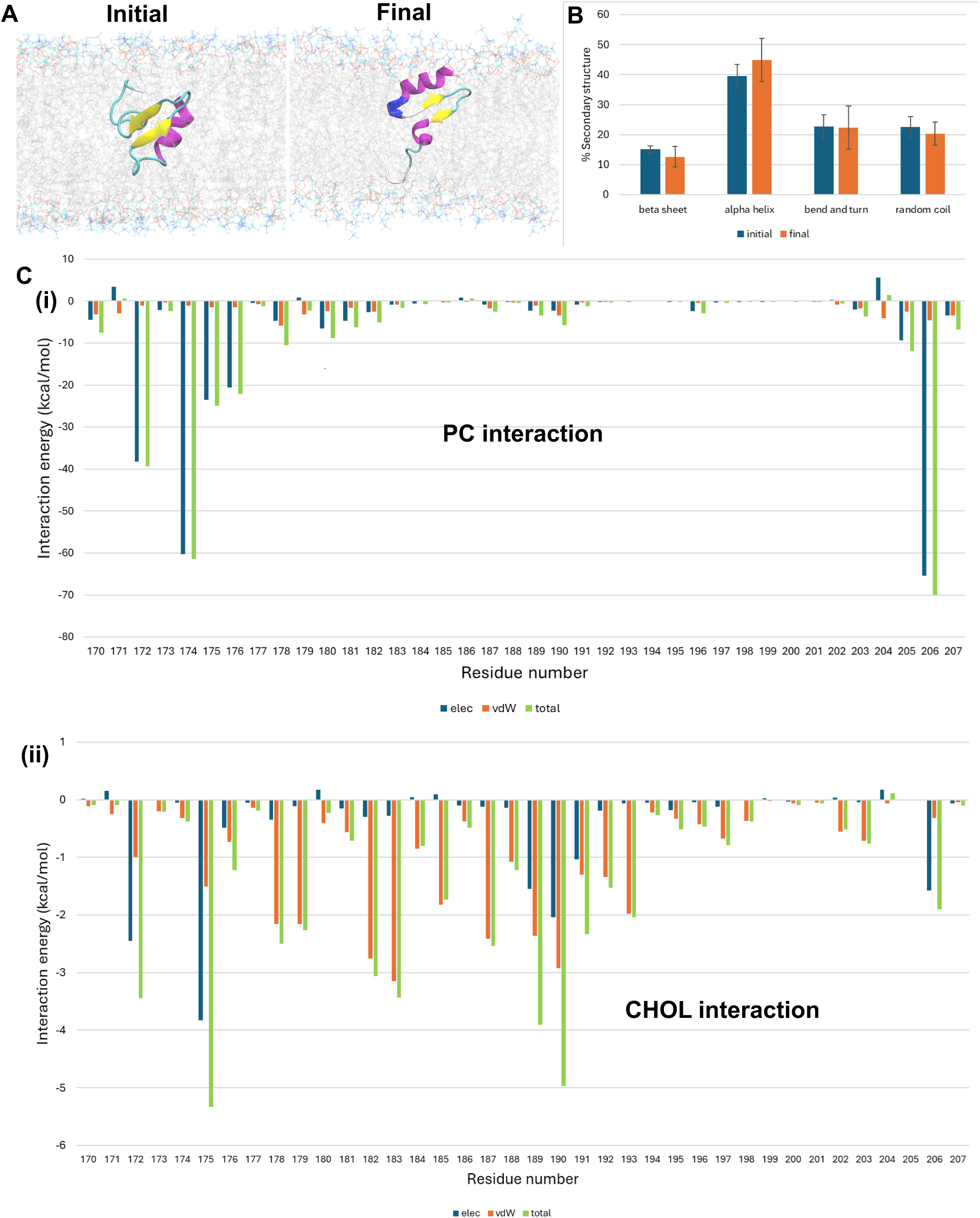
(A) Snapshots of the β-tongue segment embedded in a POPC:CHOL (70:30) lipid bilayer at the initial (0 ns) and final (300 ns) stages of molecular dynamics simulation. Structural conformations are color-coded as follows: yellow – extended β-sheet, light green – isolated bridge-β, pink – α-helix, deep blue – 3₁₀ helix, light blue – turns, and white – coil structures. The lipid bilayer is shown with headgroup atoms in blue-orange lines and tail atoms in silver. (B) Bar plot showing the percentage of each secondary structure element of the β-tongue at the initial and final simulation time points. (C) Interaction energy plots (in kcal/mol) between the β-tongue and (i) phosphatidylcholine (PC) and (ii) cholesterol (CHOL).

**Supplementary Figure S9.**
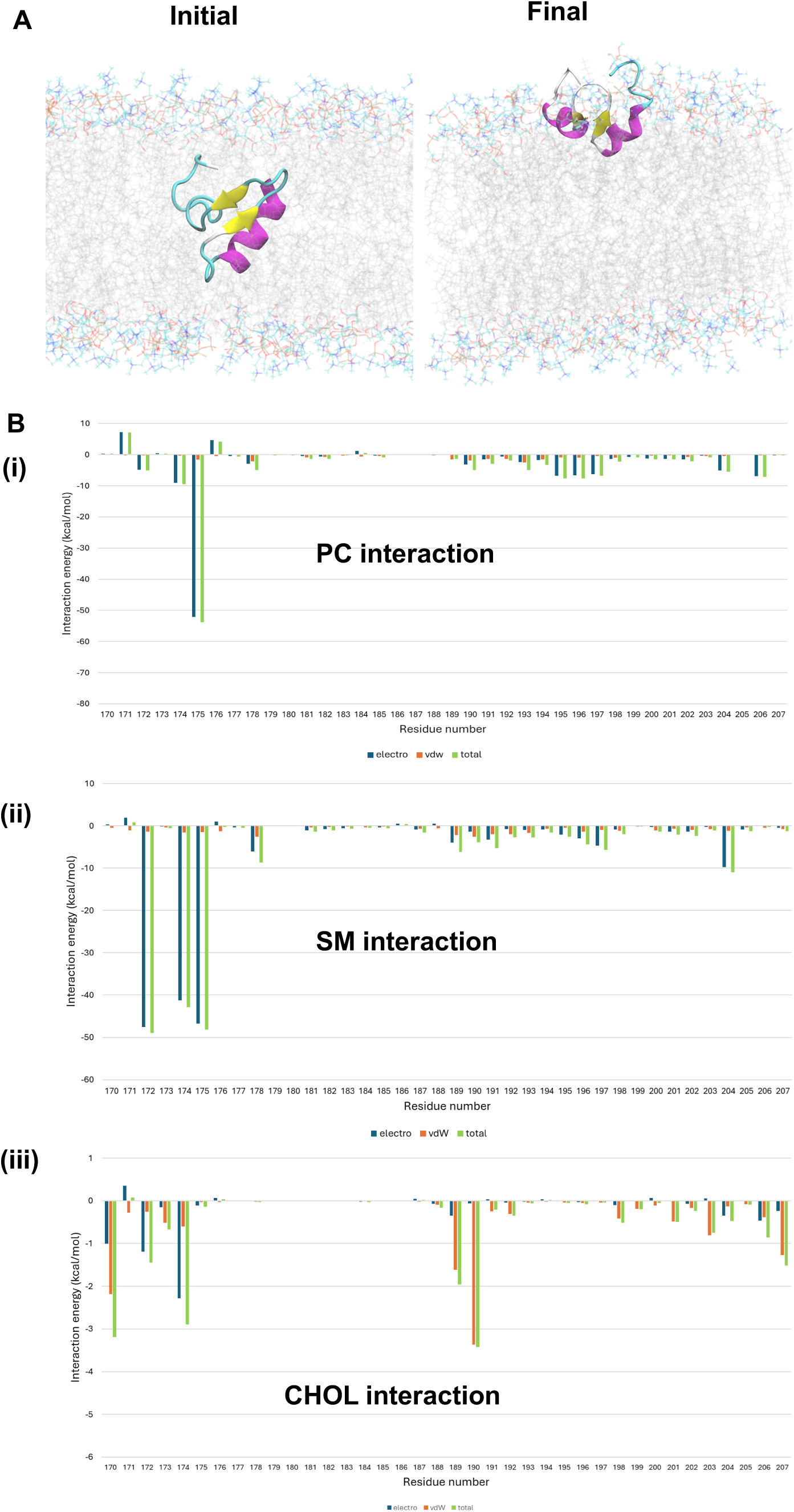
(A) Snapshots of the β-tongue segment embedded in a ternary POPC:CHOL:SM (equimolar) lipid bilayer at the initial (0 ns) and final (300 ns) stages of simulation. Structural conformations are color-coded as follows: yellow – extended β-sheet, light green – isolated bridge-β, pink – α-helix, deep blue – 3₁₀ helix, light blue – turns, and white – coil structures. The lipid bilayer is depicted with headgroup atoms in blue-orange lines and tail atoms in silver lines. (B) Interaction energy plots (kcal/mol) between the β-tongue and (i) phosphatidylcholine (PC), (ii) sphingomyelin (SM), and (iii) cholesterol (CHOL).

**Supplementary Figure S10.**
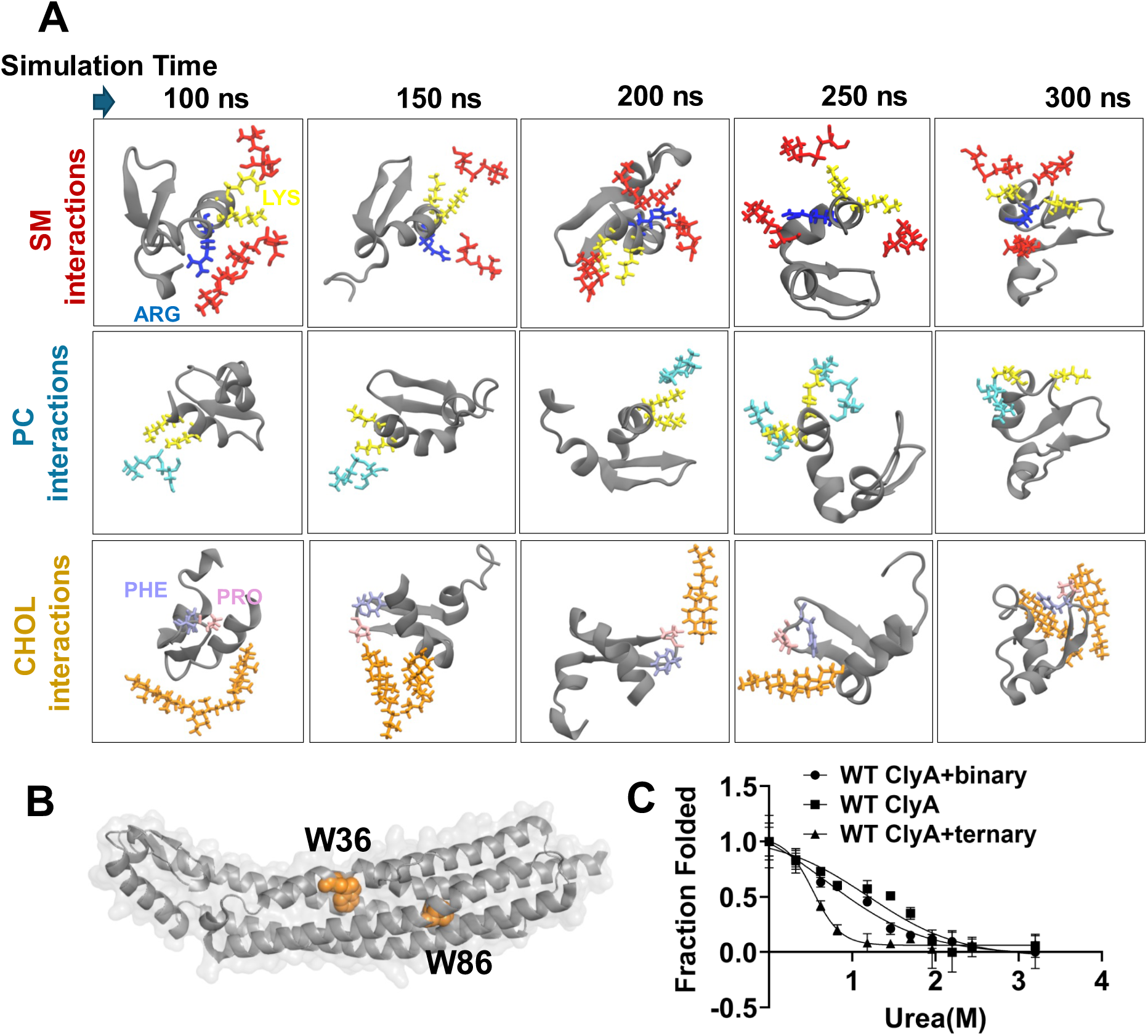
(A) Snapshots depicting interactions between lipid headgroups and specific amino acid residues of the β-tongue segment embedded in a ternary lipid membrane at various simulation time points. Color codes: grey – β-tongue segment, red – sphingomyelin (SM) headgroup, cyan – phosphatidylcholine (PC) headgroup, orange – cholesterol (CHOL) headgroup, blue – arginine (ARG), yellow – lysine (LYS), ice blue – phenylalanine (PHE), pink – proline (PRO) (B) Tryptophan residues positions within the ClyA structure. (C) Urea-induced unfolding of ClyA in the absence and presence of binary and ternary liposomes, monitored by tryptophan fluorescence as a function of denaturant concentration.

**Supplementary Figure S11.**
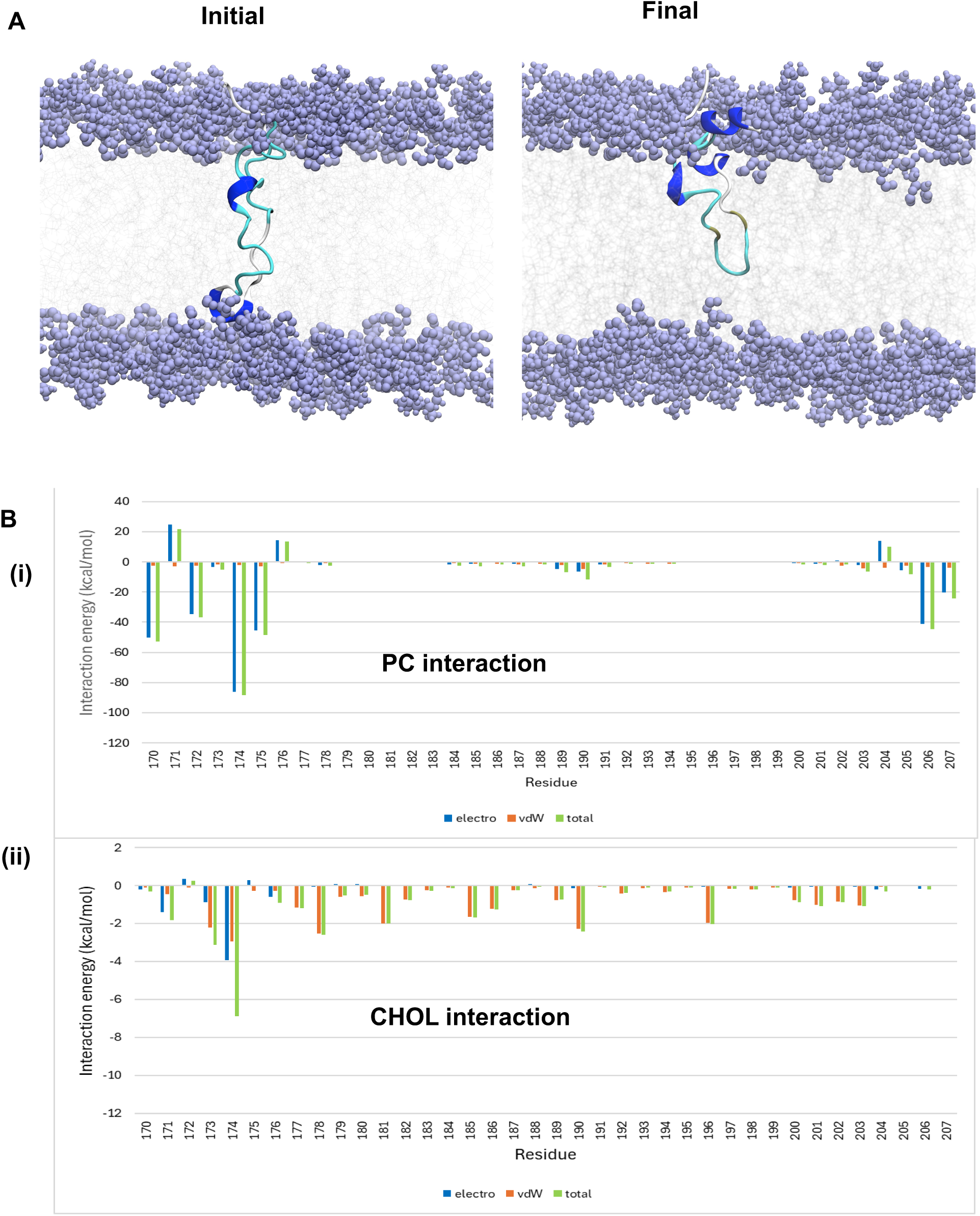
(A) Snapshots of the molten β-tongue segment of ClyA embedded in a POPC:CHOL (70:30) lipid bilayer at 200 ns and 1000 ns time points of simulation. Structural conformations are color-coded as follows: yellow – extended β-sheet, light green – isolated bridge-β, pink – α-helix, deep blue – 3₁₀ helix, light blue – turns, and white – coil structures. Lipid head atoms are shown as purple spheres, with tail atoms depicted as faint silver lines. (B) Residue-wise initial interaction energy plots (kcal/mol) between lipid components e.g. (i) phosphatidylcholine (PC) and (ii) cholesterol (CHOL) and the molten segment of the β-tongue motif of ClyA.

**Supplementary Figure S12.**
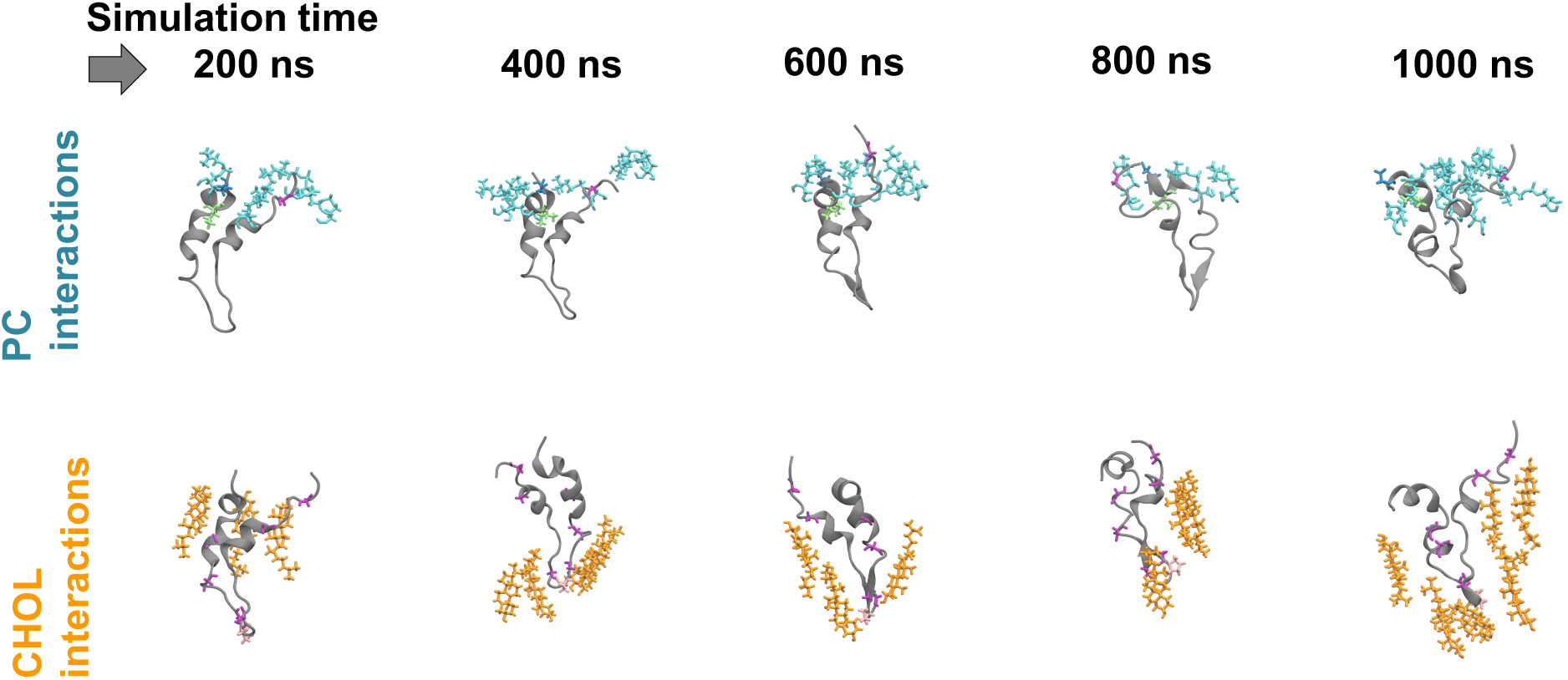
Time evolution of secondary structure changes for each residue of the molten segment within the binary lipid bilayer. Snapshots showing interactions between lipid components and specific amino acid residues of the molten segment of β-tongue embedded in a binary lipid membrane at different simulation time points. Color codes: grey – molten segment, cyan – phosphatidylcholine (PC), orange – cholesterol (CHOL), green – valine (VAL), mauve – alanine (ALA).

**Supplementary Figure S13.**
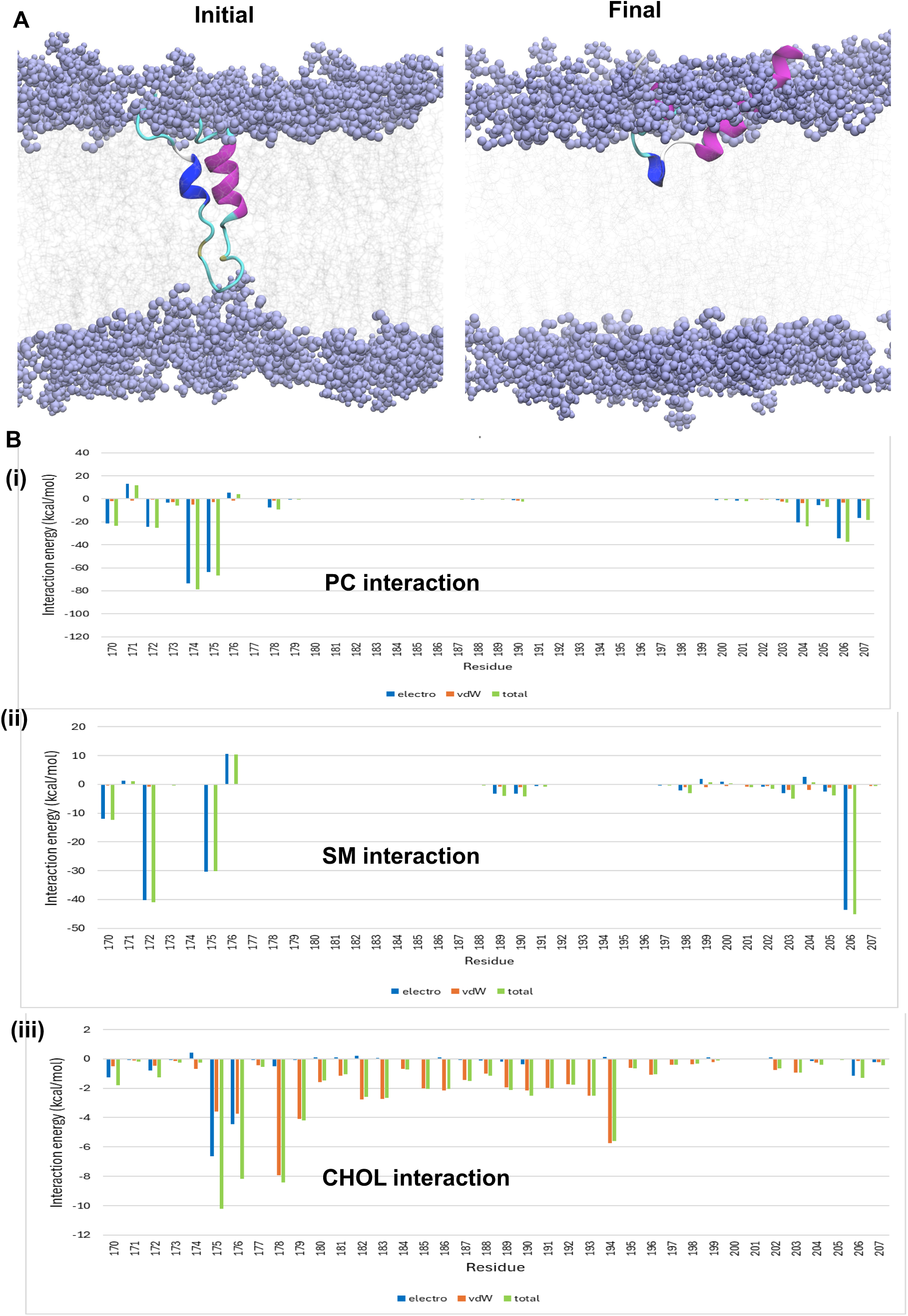
(A) Snapshots of the molten β-tongue segment of ClyA embedded in a ternary POPC:CHOL:SM (equimolar) lipid bilayer at 200 ns and 1000 ns of simulation time points. Structural conformations are color-coded as follows: yellow – extended β-sheet, light green – isolated bridge-β, pink – α-helix, deep blue – 3₁₀ helix, light blue – turns, and white – coil structures. Lipid head atoms are shown as purple spheres, and tail atoms as faint silver lines. (B) Initial interaction energy plots (kcal/mol) between lipid components and amino acid residues of the molten β-tongue segment of ClyA. Interaction profiles of molten β-tongue segment of ClyA with (i) phosphatidylcholine (PC), (ii) sphingomyelin (SM), and (iii) cholesterol (CHOL) are shown.

**Supplementary Figure S14.**
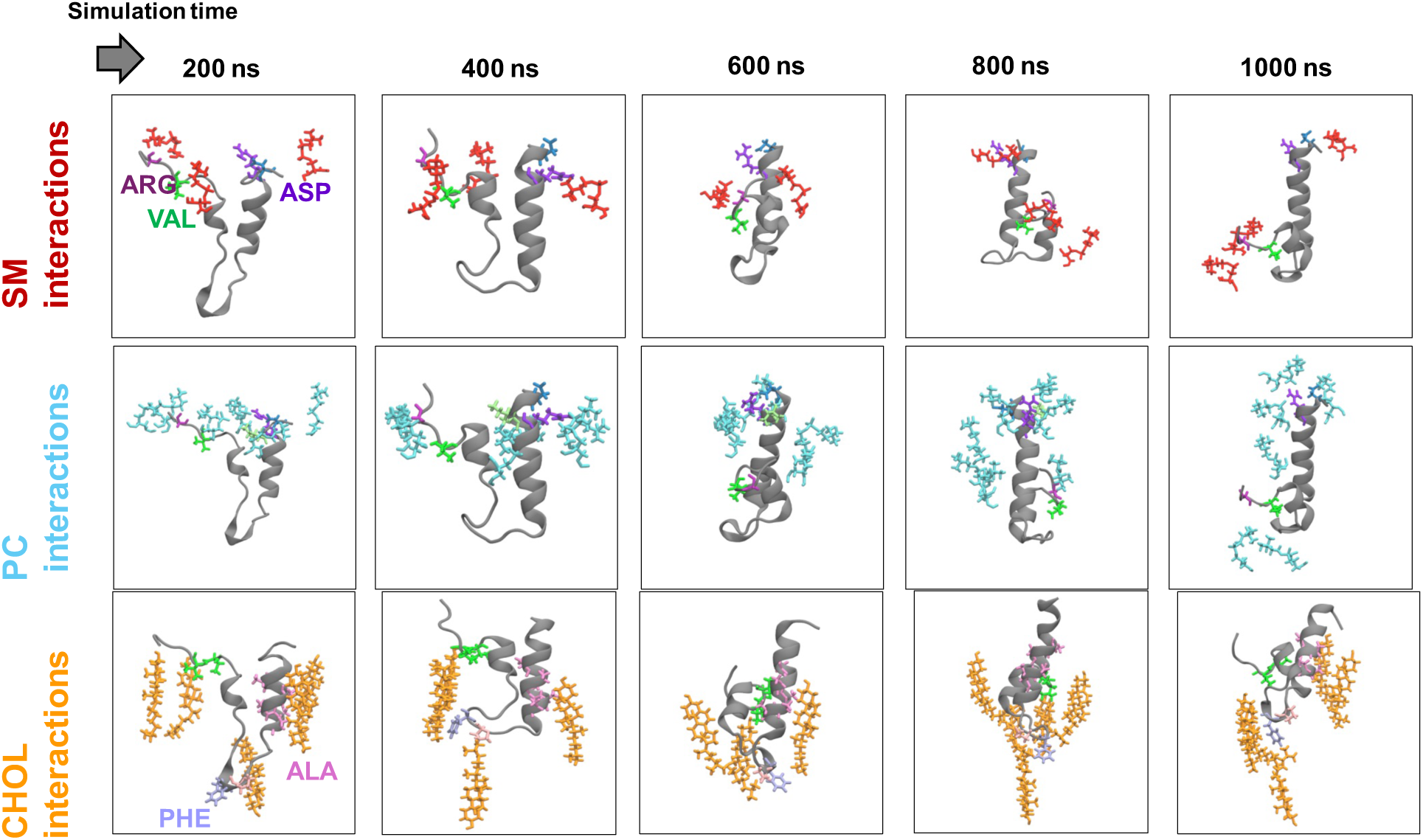
Time evolution of structural changes for specific residues of the molten β-tongue segment of ClyA within the ternary lipid bilayer. Snapshots showing interactions between lipid components and specific amino acid residues of the molten β-tongue segment of ClyA embedded in a ternary lipid membrane at various simulation time points. Color codes: grey – molten segment, red – sphingomyelin (SM) headgroup, cyan – phosphatidylcholine (PC) headgroup, orange – cholesterol (CHOL) headgroup, green – valine (VAL), violet – arginine (ARG), blue – aspartic acid (ASP), pink – alanine (ALA), ice blue – phenylalanine (PHE).

**Supplementary Figure S15.**
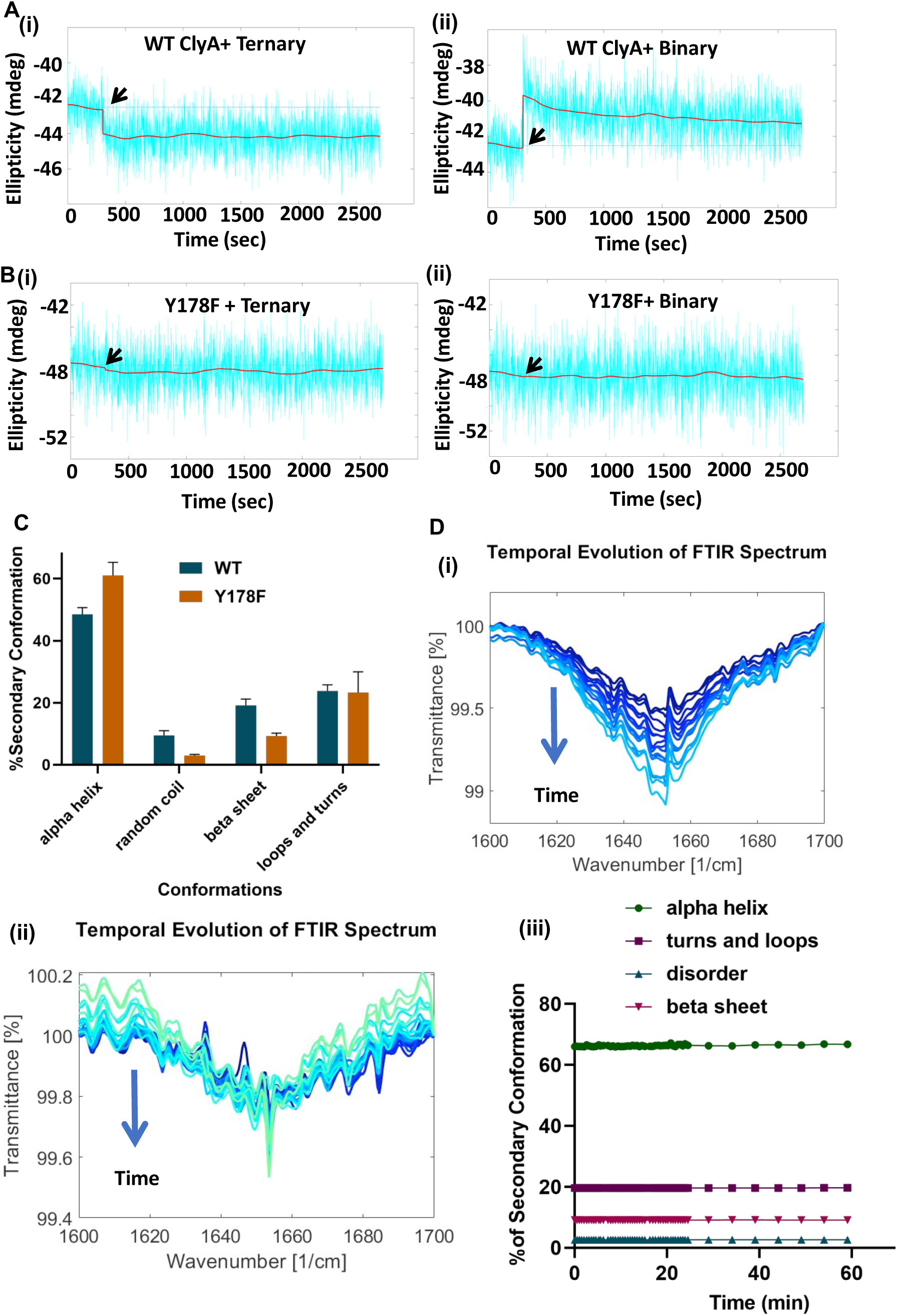
Conformational changes of WT ClyA and Y178F ClyA in the presence of membranes. (A) CD spectra showing the kinetics of α-helical content (at 222 nm) in WT ClyA following immediate addition (indicated by arrow mark) of (i) ternary (POPC:SM:CHOL equimolar) and (ii) binary (POPC:CHOL 70:30) liposomes. (B) CD spectra showing α-helical kinetics in Y178F ClyA after immediate addition (indicated by arrow mark) of (i) ternary(POPC:SM:CHOL equimolar) and (ii) binary (POPC:CHOL 70:30) liposomes. (C) Relative contents of different conformational states of WT ClyA and Y178F ClyA determined by ATR-FTIR spectroscopy. (D) Temporal evolution of FTIR spectra for (i) WT ClyA and (ii) Y178F ClyA after addition of binary liposomes; data indicate insignificant spectral changes for Y178F ClyA in the membrane environment. (iii) Temporal evolution of secondary structure elements for Y178F ClyA.

**Supplementary Figure S16.**
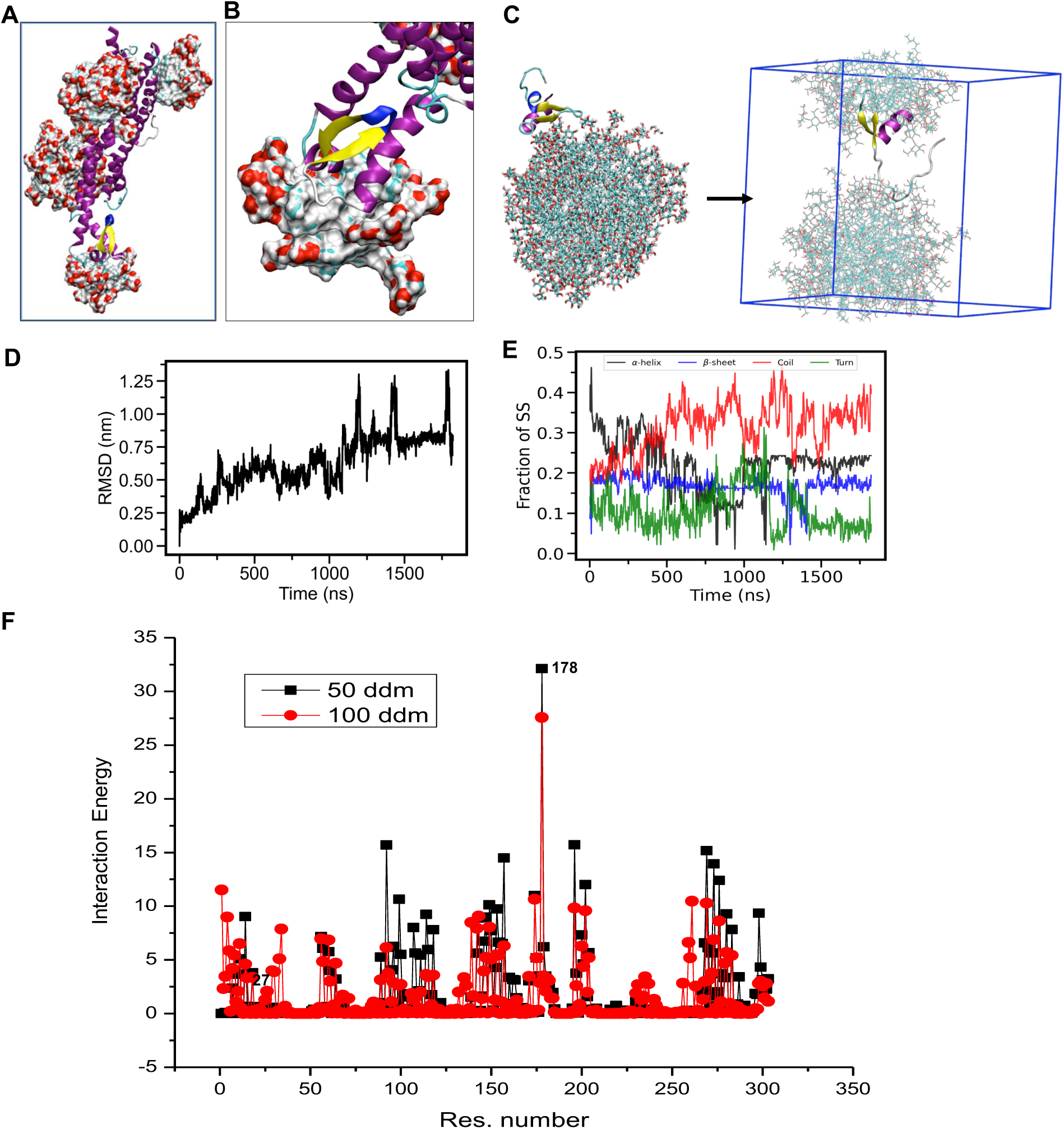
Molecular dynamics simulation of WT ClyA in DDM micelles. (A) Final snapshot showing the WT ClyA monomer and surrounding DDM molecules; (B) Zoomed-in view of the β-tongue region with DDM molecules displayed in surface representation. (C) Initial (left) and final (right) configurations of the β-tongue exposed to the DDM micelle; the box indicates periodic boundary conditions. (D) Root mean square deviation (RMSD), (E) fraction of secondary structure content, and (F) relative interaction energy (Kcal/mol) of the β-tongue segment in presence of DDM during the simulation.

**Supplementary Figure S17.**
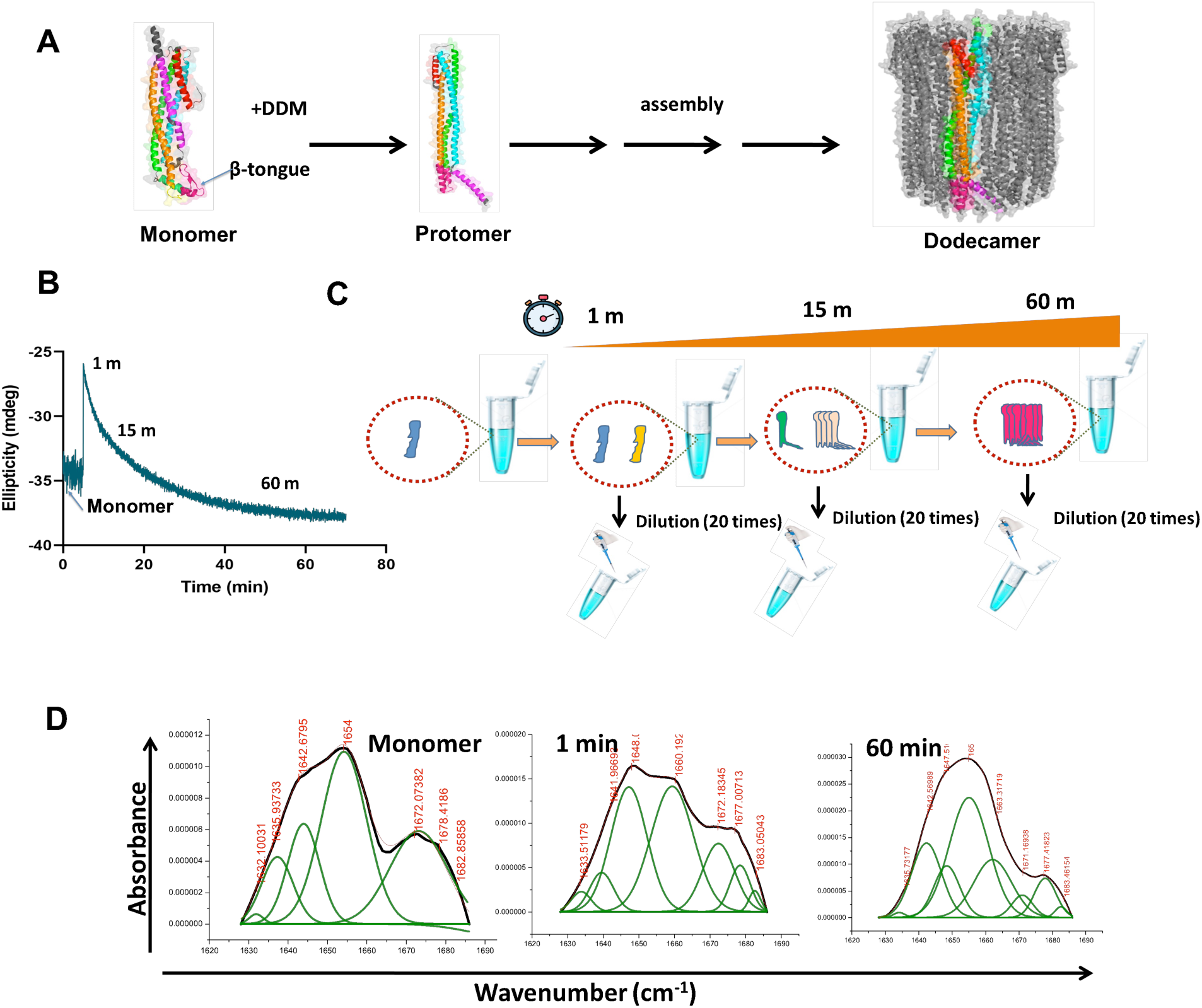
Different states of WT ClyA during the assembly process. (A) Conformational transition of ClyA from monomer to protomer and finally to the dodecameric pore structure upon interaction with 0.1% (w/v) DDM. (B) CD spectra showing the temporal changes in α-helical content of WT ClyA after addition of DDM at different time points (1 min, 15 min, and 60 min), representing three distinct conformational states: 1 min = Intermediate(I), 15 min = Incomplete pore (InP), and 60 min = Complete pore (cP) (C) Pictorial diagram illustrating the three WT ClyA states during transition; samples were diluted 20-fold in PBS to halt the assembly process. (D) ATR-FTIR spectra of monomeric WT ClyA and its three states after exposure to 0.1% DDM. (E) Deconvoluted FTIR spectra of (i) monomer WT ClyA, (ii) intermediate state ‘I’(1 min sample), and (iii) complete pore state cP(60 min sample).

**Supplementary Figure S18.**
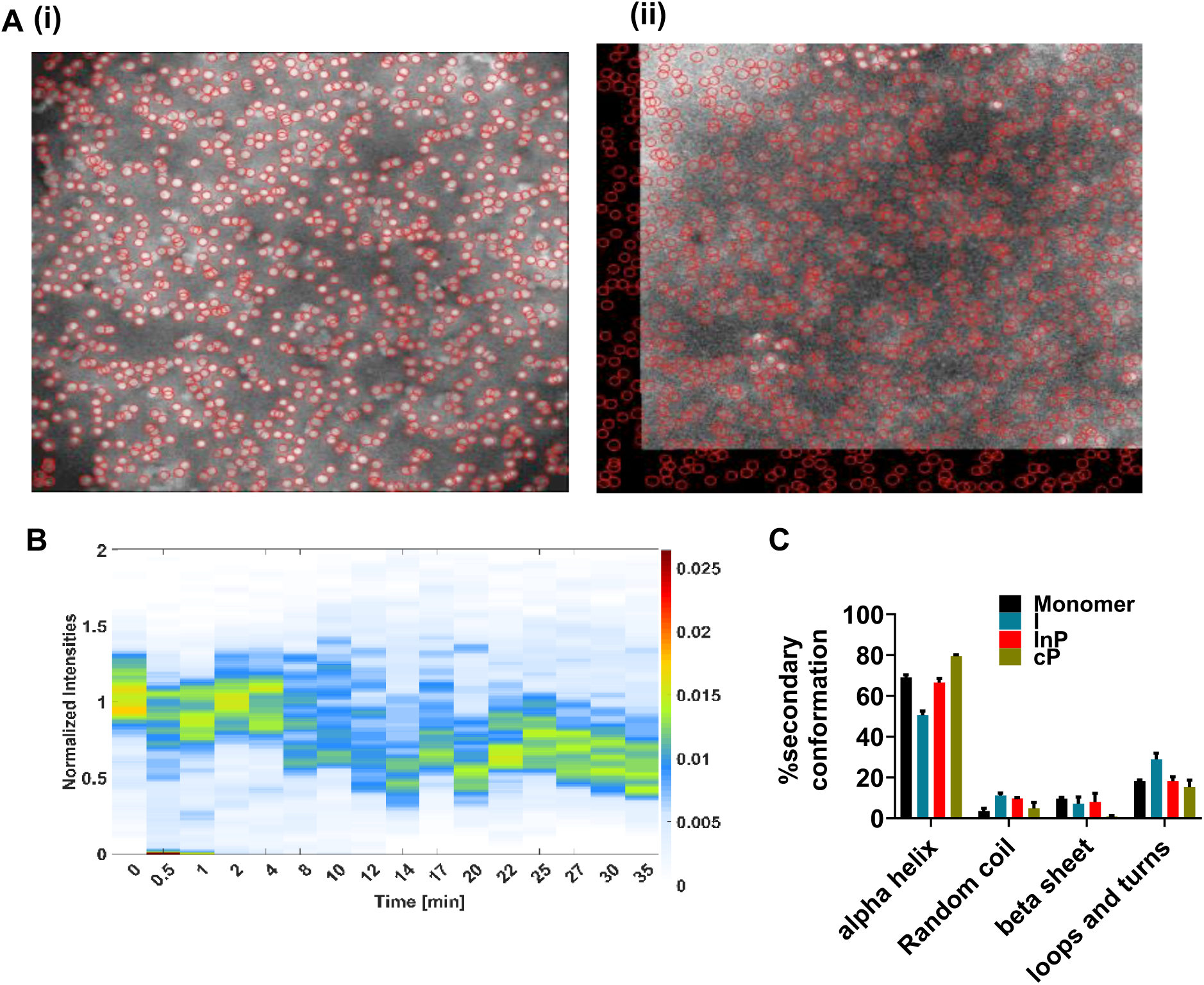
(A) Fluorescence micrographs of SRB-entrapped ternary SuLB membrane (POPC:SM:CHOL equimolar) before (i) and after (ii) addition of WT ClyA ‘I’ state (30 min post-addition) (B) Temporal SRB leakage from ternary SuLB (POPC:SM:CHOL equimolar) induced by the ‘cP’ state; color bar indicates probability density. (C) Percentage of different secondary structures of Y178F ClyA and its three DDM-induced states: I (Y178F), InP (Y178F), and cP (Y178F). Data are presented as Mean ± SD (n = 3), where n denotes the number of experimental repeats.

**Supplementary Figure S19.**
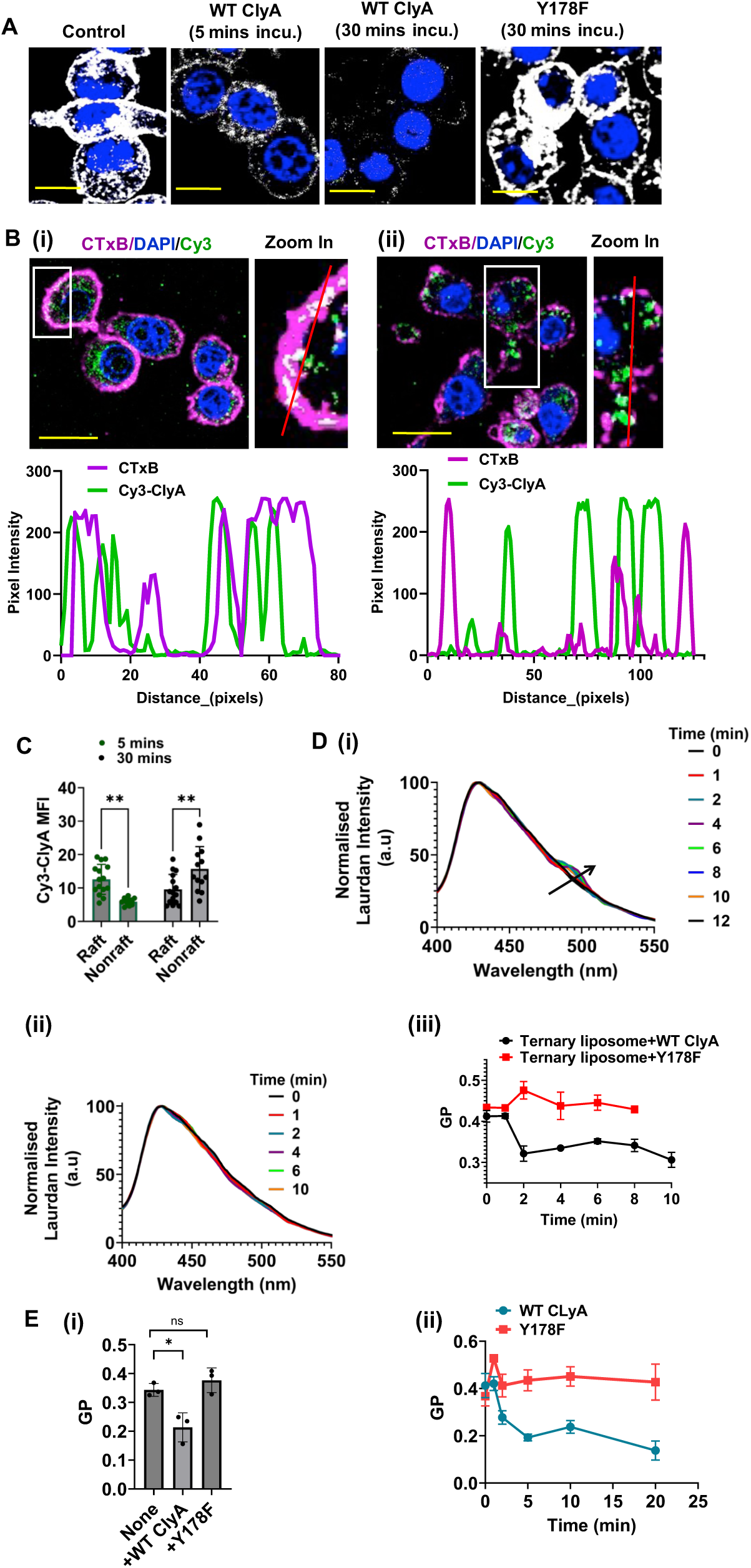
(A) Confocal microscopy images show the lipid raft organization in RAW cells after treatment with WT ClyA for 5 and 20 minutes, or with the Y178F ClyA for 20 minutes, as visualized by CTxB-Alexa 647 binding. Scale bar: 10 µm. (B) Confocal images display co-localization of CTxB (magenta), Cy3-ClyA (green), and DAPI (blue) following incubation with Cy3-ClyA for (i) 5 minutes and (ii) 30 minutes. Zoomed-in panels highlight membrane regions showing co-localization of CTxB and Cy3-ClyA. Line scan profiles (red lines) indicate overlapping fluorescence intensities of CTxB and Cy3-ClyA. Scale bars: 10 µm. (C) Quantitative analysis comparing the mean fluorescence intensity (MFI) of Cy3-ClyA in raft (CTxB-positive) versus non-raft regions at 5 and 30 minutes post-treatment. Data are shown as mean ± SD (n = 3). Statistical significance was determined by two-way ANOVA using GraphPad Prism (version 9); ****P = 0.0018 (5 min group) and ****P = 0.0024 (30 min group). (D) Temporal fluorescence spectra of laurdan-labeled ternary liposomes (POPC:SM:CHOL equimolar) after treatment with (i) WT ClyA and (ii) Y178F ClyA, along with (iii) a plot showing time-dependent changes in laurdan generalized polarization (GP) following treatment. Data are presented as mean ± SD (n = 3), where *n* represents the number of independent experimental replicates. (E) (i) Raw cells labeled with laurdan were treated with WT ClyA or Y178F ClyA at 100 nM concentration and incubated for 30 mins and generalized polarization (GP) was measured. Data were represented as mean ± SD (n = 3). Statistical significance was determined by two-way ANOVA using GraphPad Prism (version 9); *P = 0.02. (ii) Plot shows the change in GP values with time after treatment of Raw cells with 100 nM WT ClyA or Y178F ClyA.

**Supplementary Figure S20.**
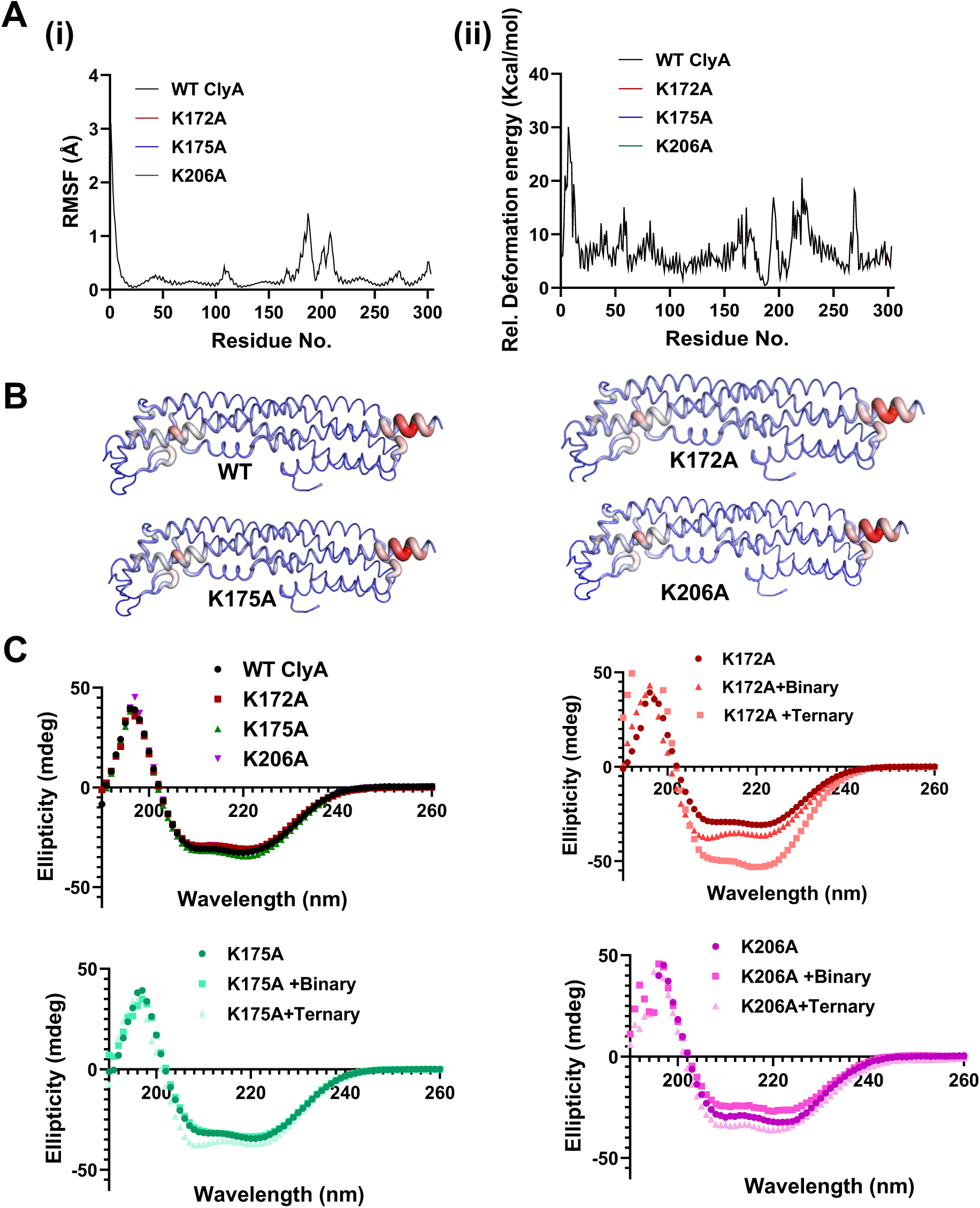
A (i) Plot showing the residue wise fluctuations of WT ClyA and four lysine-alanine mutants. (ii) Relative deformation energy per residue has been plotted for WT and four mutants of ClyA. (B) Visual analysis of Deformation Energies. Deformation energy provides a measure for the amount of local flexibility in the protein. Calculations performed over the first 10 non-trivial modes of the molecule. The magnitude of the fluctuation is represented by thin to thick tube colored blue (low), white (moderate) and red (high). (C) Far-UV CD spectra of WT ClyA and three alanine mutants: K172A, K175A, and K206A. Secondary structure changes were monitored by measuring CD signatures of K172A, K175A, and K206A in the presence and absence of binary(POPC:CHOL 70:30) or ternary(POPC:SM:CHOL equimolar) liposomes. Background subtraction was performed for each protein sample.

**Supplementary Figure S21.**
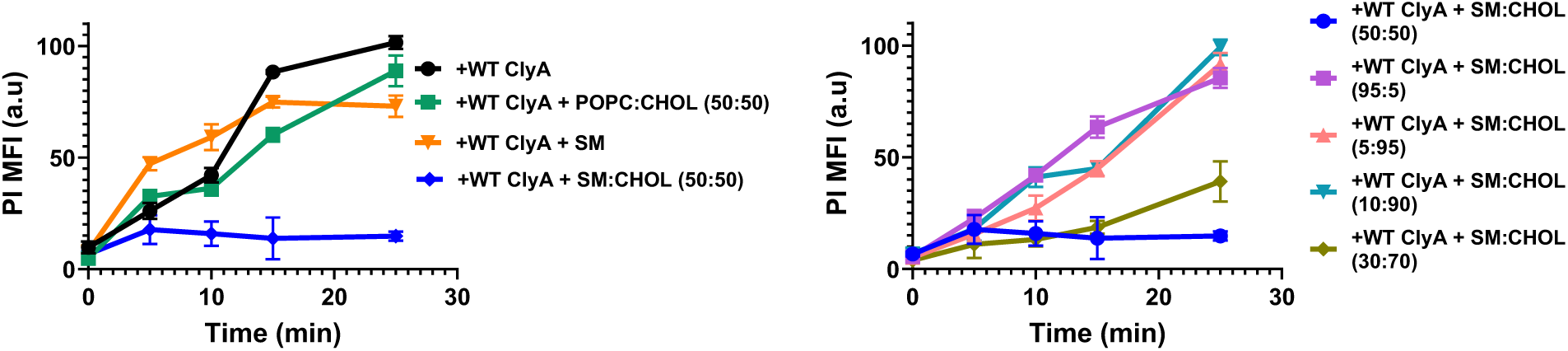
Membrane pore-formation assay by WT ClyA by monitoring the PI uptake in RAW cell. RAW 264.7 cells (2×10^4^/well) were seeded in 96-well plates, cultured in DMEM supplemented with 10% FBS and 1% penicillin-streptomycin (Day 0). Day 1: cells were treated with WT ClyA (100 nM) along with liposomes composed of different lipid components and the temporal PI intensities inside the cells were recorded by fluorescence microscopic imaging to track the pore-formation kinetics of WT ClyA.

**Supplementary Figure S22.**
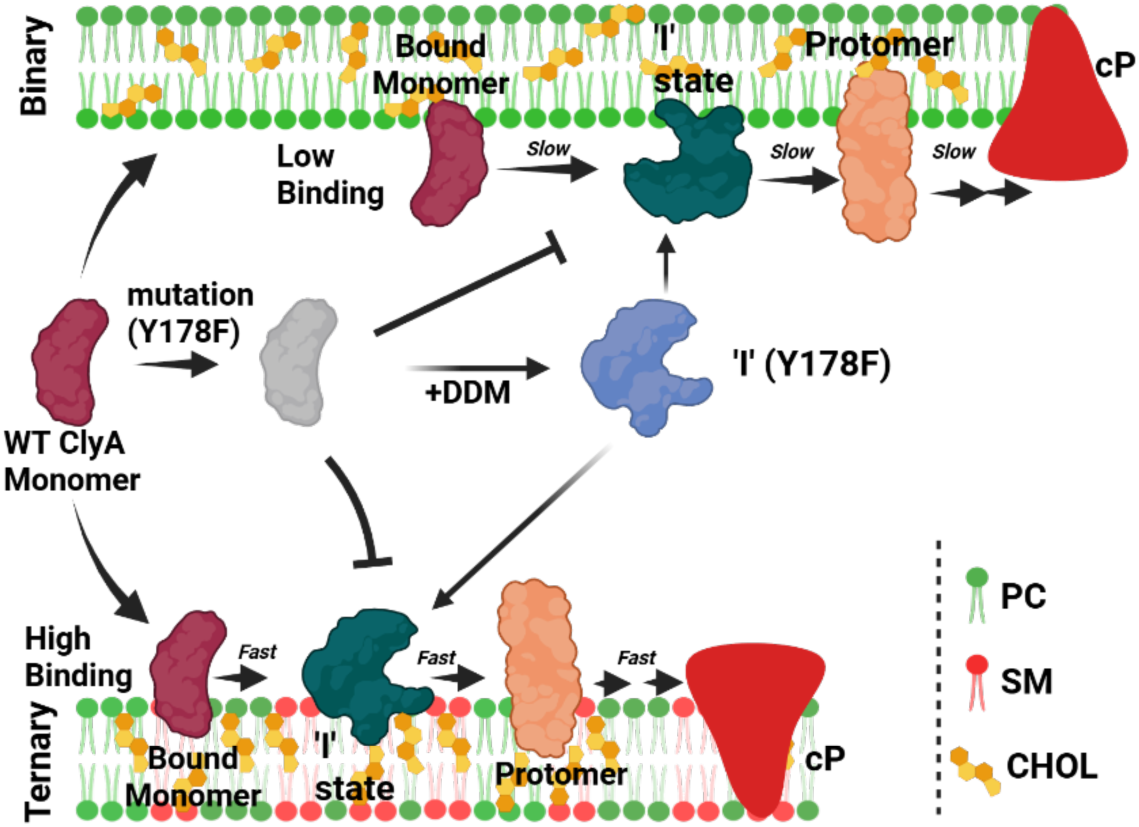
Schematic illustration highlighting that early melting and the presence of sphingomyelin in the bilayer are crucial for the conformational transition of ClyA. The Y178F mutant, which is defective in lysing cell membranes, regains its lytic activity upon detergent (DDM)-induced conformational melting.

**Supplementary Figure S23.**
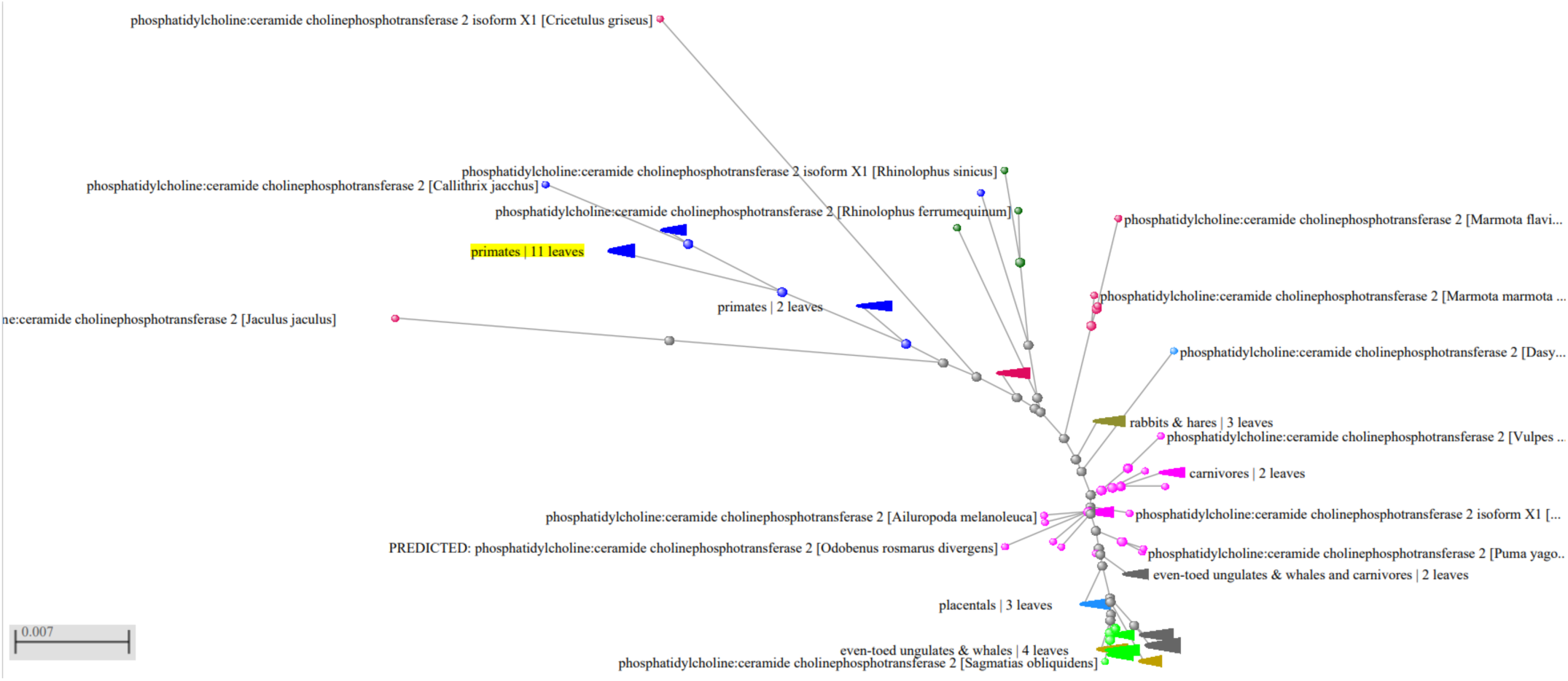
Phylogenetic tree of the sphingomyelin synthase 2 (SMS2) protein.

**Supplementary Figure S24.**
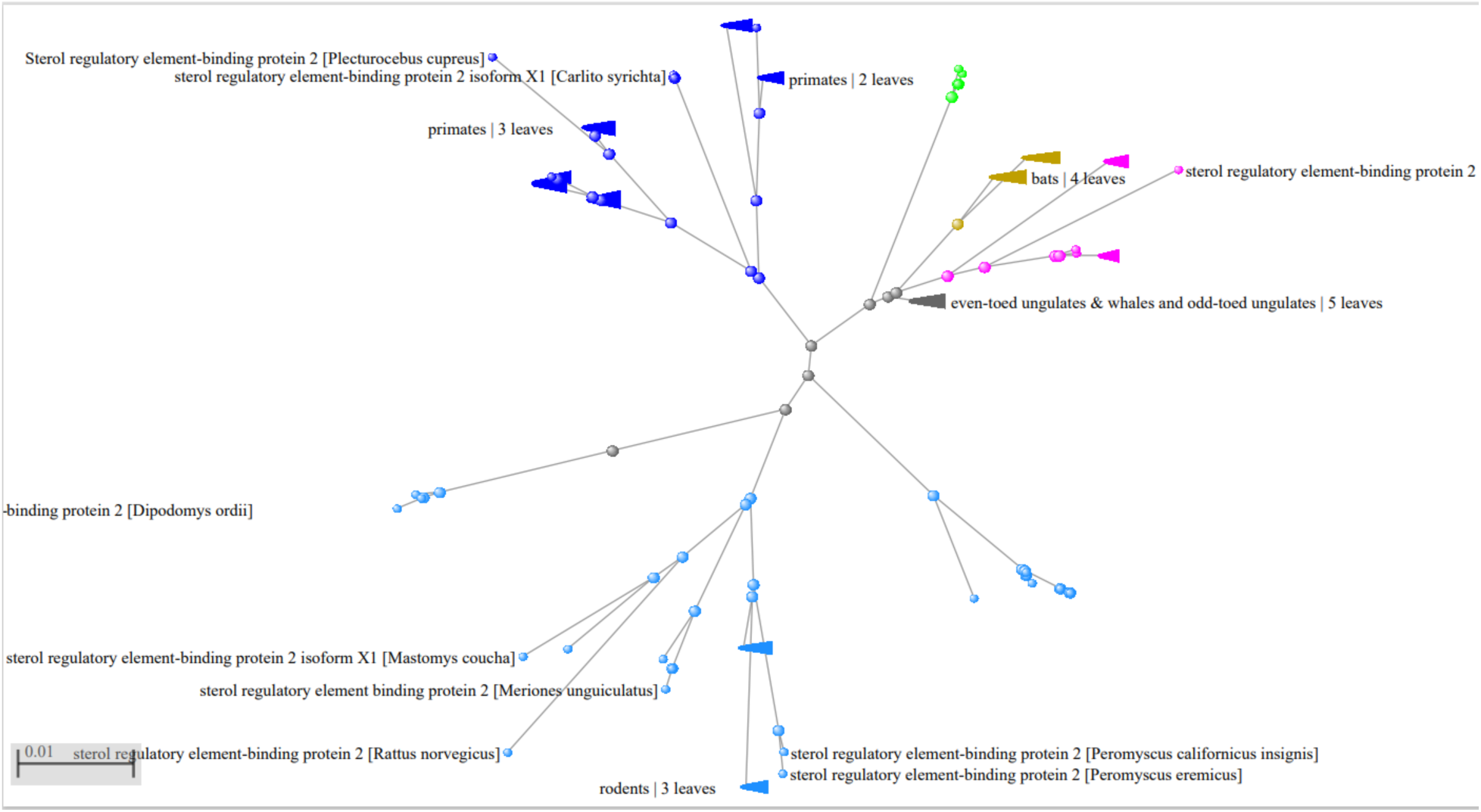
Phylogenetic tree of SREBP2, a key protein in cholesterol biosynthesis.

**Supplementary Figure S25.**
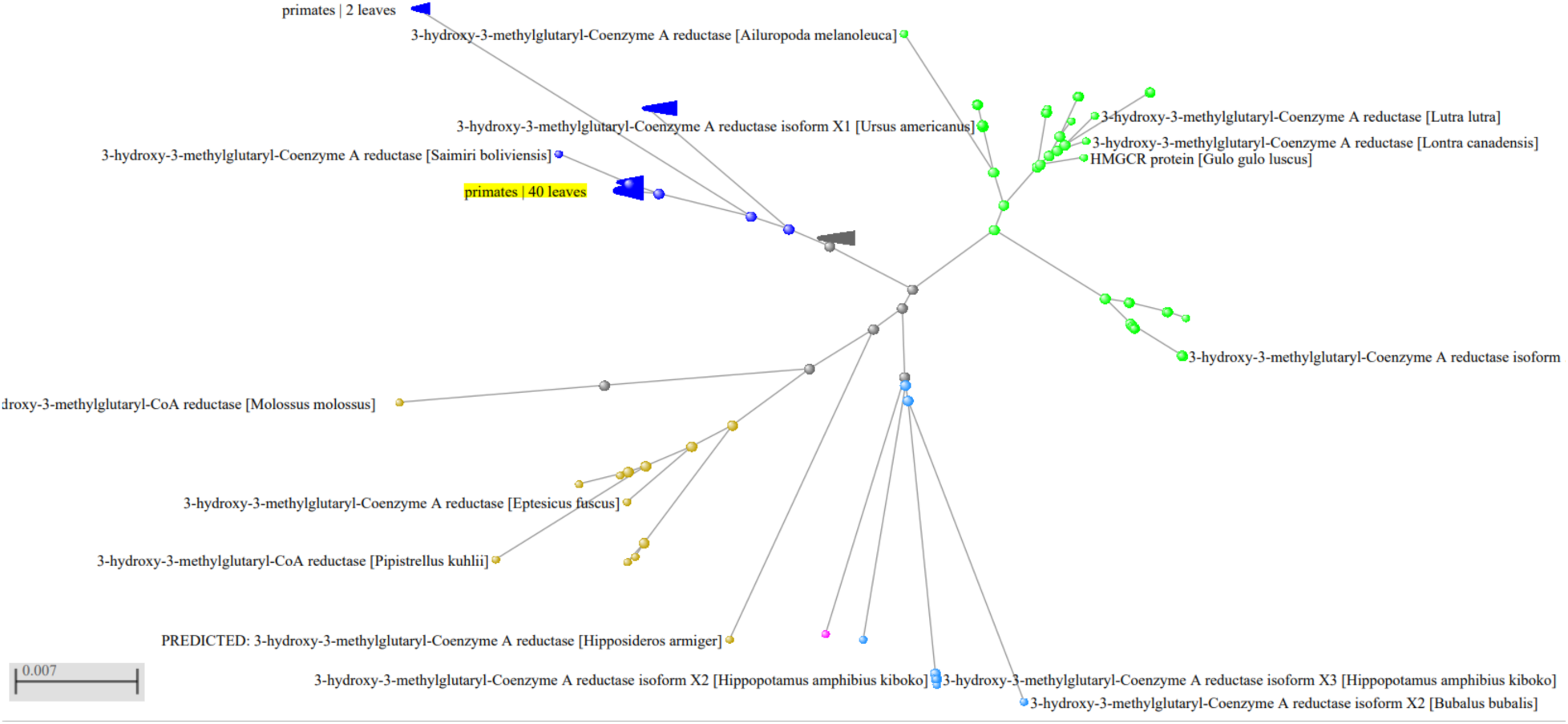
Phylogenetic tree of HMGCR, a key enzyme in cholesterol biosynthesis.

## References

Åqvist J, Wennerström P, Nervall M, Bjelic S, Brandsdal BO (2004) Molecular dynamics simulations of water and biomolecules with a Monte Carlo constant pressure algorithm. Chemical physics letters 384: 288–294

Bandyopadhyay A, Sannigrahi A, Chattopadhyay K (2021) Membrane composition and lipid to protein ratio modulate amyloid kinetics of yeast prion protein. RSC Chemical Biology 2: 592–605

Becucci L, Vizza F, Duarte Y, Guidelli R (2014) The GM1 ganglioside forms GM1-rich gel phase microdomains within lipid rafts. Coatings 4: 450–464

Benke S, Roderer D, Wunderlich B, Nettels D, Glockshuber R, Schuler B (2015) The assembly dynamics of the cytolytic pore toxin ClyA. Nature communications 6: 6198

Casadevall A, Pirofski L-a (1999) Host-pathogen interactions: redefining the basic concepts of virulence and pathogenicity. Infection and immunity 67: 3703–3713

Case DA, Aktulga HM, Belfon K, Ben-Shalom IY, Berryman JT, Brozell SR, Cerutti DS, Cheatham III TE, Cisneros GA, Cruzeiro VWD (2023) *Amber* 2023. University of California, San Francisco

Das A, Brown MS, Anderson DD, Goldstein JL, Radhakrishnan A (2014) Three pools of plasma membrane cholesterol and their relation to cholesterol homeostasis. Elife 3

Dickson CJ, Walker RC, Gould IR (2022) Lipid21: complex lipid membrane simulations with AMBER. Journal of chemical theory and computation 18: 1726–1736

Endapally S, Frias D, Grzemska M, Gay A, Tomchick DR, Radhakrishnan A (2019) Molecular discrimination between two conformations of sphingomyelin in plasma membranes. Cell 176: 1040–1053. e1017

Essmann U, Perera L, Berkowitz ML, Darden T, Lee H, Pedersen LG (1995) A smooth particle mesh Ewald method. The Journal of chemical physics 103: 8577–8593

Fahie MA, Liang L, Avelino AR, Pham B, Limpikirati P, Vachet RW, Chen M (2018) Disruption of the open conductance in the beta-tongue mutants of Cytolysin A. Sci Rep 8: 3796

Flores-Díaz M, Monturiol-Gross L, Naylor C, Alape-Girón A, Flieger A (2016) Bacterial Sphingomyelinases and Phospholipases as Virulence Factors. Microbiol Mol Biol Rev 80: 597–628

Gatt S, Bierman EL (1980) Sphingomyelin suppresses the binding and utilization of low density lipoproteins by skin fibroblasts. Journal of Biological Chemistry 255: 3371–3376

Gebhardt M, Henkes LM, Tayefeh S, Hertel B, Greiner T, Van Etten JL, Baumeister D, Cosentino C, Moroni A, Kast SM (2012) Relevance of lysine snorkeling in the outer transmembrane domain of small viral potassium ion channels. Biochemistry 51: 5571–5579

Giri Rao VV, Desikan R, Ayappa KG, Gosavi S (2016) Capturing the Membrane-Triggered Conformational Transition of an alpha-Helical Pore-Forming Toxin. J Phys Chem B 120: 12064–12078

Gonzalez MR, Bischofberger M, Frêche B, Ho S, Parton RG, van der Goot FG (2011) Pore-forming toxins induce multiple cellular responses promoting survival. Cellular microbiology 13: 1026–1043

Ha JH, Loh SN (2012) Protein conformational switches: from nature to design. Chemistry 18: 7984–7999

Halder A, Sannigrahi A, De N, Chattopadhyay K, Karmakar S (2020) Kinetoplastid membrane protein-11 induces pores in anionic phospholipid membranes: effect of cholesterol. Langmuir 36: 3522–3530

Hartl FU, Hayer-Hartl M (2002) Molecular chaperones in the cytosol: from nascent chain to folded protein. Science 295: 1852–1858

Heuck AP, Moe PC, Johnson BB (2010) The cholesterol-dependent cytolysin family of gram-positive bacterial toxins. Subcell Biochem 51: 551–577

Huang H, Guo W, Zhang Y, 2008. Detection of copy-move forgery in digital images using SIFT algorithm, 2008 IEEE Pacific-Asia Workshop on Computational Intelligence and Industrial Application. IEEE, pp. 272–276.

Humphrey W, Dalke A, Schulten K (1996) VMD: visual molecular dynamics. Journal of molecular graphics 14: 33–38

Hünenberger PH (2005) Thermostat algorithms for molecular dynamics simulations. Advanced computer simulation: Approaches for soft matter sciences I: 105–149

Ishitsuka R, Yamaji-Hasegawa A, Makino A, Hirabayashi Y, Kobayashi T (2004) A lipid-specific toxin reveals heterogeneity of sphingomyelin-containing membranes. Biophysical journal 86: 296–307

Jo S, Lim JB, Klauda JB, Im W (2009) CHARMM-GUI Membrane Builder for mixed bilayers and its application to yeast membranes. Biophysical journal 97: 50–58

Jorgensen WL, Chandrasekhar J, Madura JD, Impey RW, Klein ML (1983) Comparison of simple potential functions for simulating liquid water. The Journal of chemical physics 79: 926–935

Joung IS, Cheatham III TE (2008) Determination of alkali and halide monovalent ion parameters for use in explicitly solvated biomolecular simulations. The journal of physical chemistry B 112: 9020–9041

Kostow N, Welch MD (2023) Manipulation of host cell plasma membranes by intracellular bacterial pathogens. Current opinion in microbiology 71: 102241

Krone L, Mahankali S, Geiger T (2024) Cytolysin A is an intracellularly induced and secreted cytotoxin of typhoidal Salmonella. Nat Commun 15: 8414

Kulshrestha A, Maurya S, Gupta T, Roy R, Punnathanam SN, Ayappa KG (2023a) Conformational Flexibility Is a Key Determinant for the Lytic Activity of the Pore-Forming Protein, Cytolysin A. J Phys Chem B 127: 69–84

Kulshrestha A, Punnathanam SN, Roy R, Ayappa KG (2023b) Cholesterol catalyzes unfolding in membrane-inserted motifs of the pore forming protein cytolysin A. Biophysical Journal 122: 4068–4081

Lakshminarayan R, Wunder C, Becken U, Howes MT, Benzing C, Arumugam S, Sales S, Ariotti N, Chambon V, Lamaze C (2014) Galectin-3 drives glycosphingolipid-dependent biogenesis of clathrin-independent carriers. Nature cell biology 16: 592–603

Landolt-Marticorena C, Williams KA, Deber CM, Reithmeier RA, 1993. Non-random distribution of amino acids in the transmembrane segments of human type I single span membrane proteins. Elsevier, pp. 602–608.

Li L, Vorobyov I, Allen TW (2013) The different interactions of lysine and arginine side chains with lipid membranes. The journal of physical chemistry B 117: 11906–11920

Li Y-J, Chen C-Y, Yang J-H, Chiu Y-F (2022) Modulating cholesterol-rich lipid rafts to disrupt influenza A virus infection. Frontiers in Immunology 13: 982264

Lingwood D, Simons K (2010) Lipid rafts as a membrane-organizing principle. science 327: 46–50

Lomize MA, Pogozheva ID, Joo H, Mosberg HI, Lomize AL (2012) OPM database and PPM web server: resources for positioning of proteins in membranes. Nucleic acids research 40: D370–D376

Lorent JH, Levental KR, Ganesan L, Rivera-Longsworth G, Sezgin E, Doktorova M, Lyman E, Levental I (2020) Plasma membranes are asymmetric in lipid unsaturation, packing and protein shape. Nature chemical biology 16: 644–652

Maier JA, Martinez C, Kasavajhala K, Wickstrom L, Hauser KE, Simmerling C (2015) ff14SB: improving the accuracy of protein side chain and backbone parameters from ff99SB. Journal of chemical theory and computation 11: 3696–3713

Makino A, Abe M, Murate M, Inaba T, Yilmaz N, Hullin-Matsuda F, Kishimoto T, Schieber NL, Taguchi T, Arai H (2015) Visualization of the heterogeneous membrane distribution of sphingomyelin associated with cytokinesis, cell polarity, and sphingolipidosis. The FASEB Journal 29: 477–493

Meng Y, Zhang Z, Yin H, Ma T (2018) Automatic detection of particle size distribution by image analysis based on local adaptive canny edge detection and modified circular Hough transform. Micron 106: 34–41

Moore JW, Stanitski CL (1998) The chemical world: concepts and applications. (No Title)

Mueller M, Grauschopf U, Maier T, Glockshuber R, Ban N (2009) The structure of a cytolytic α-helical toxin pore reveals its assembly mechanism. Nature 459: 726–730

Omelyan I, Kovalenko A (2013) Multiple time step molecular dynamics in the optimized isokinetic ensemble steered with the molecular theory of solvation: Accelerating with advanced extrapolation of effective solvation forces. The Journal of Chemical Physics 139

Radhakrishnan A, Rohatgi R, Siebold C (2020) Cholesterol access in cellular membranes controls Hedgehog signaling. Nature chemical biology 16: 1303–1313

Rodrigues CH, Pires DE, Ascher DB (2018) DynaMut: predicting the impact of mutations on protein conformation, flexibility and stability. Nucleic acids research 46: W350–W355

Roe DR, Cheatham III TE (2013) PTRAJ and CPPTRAJ: software for processing and analysis of molecular dynamics trajectory data. Journal of chemical theory and computation 9: 3084–3095

Róg T, Pasenkiewicz-Gierula M (2006) Cholesterol-sphingomyelin interactions: a molecular dynamics simulation study. Biophysical Journal 91: 3756–3767

Rojko N, Anderluh G (2015) How lipid membranes affect pore forming toxin activity. Accounts of chemical research 48: 3073–3079

Ryckaert J-P, Ciccotti G, Berendsen HJ (1977) Numerical integration of the cartesian equations of motion of a system with constraints: molecular dynamics of n-alkanes. Journal of computational physics 23: 327–341

Sannigrahi A, Chall S, Jawed JJ, Kundu A, Majumdar S, Chattopadhyay K (2018) Nanoparticle induced conformational switch between α-helix and β-sheet attenuates immunogenic response of MPT63. Langmuir 34: 8807–8817

Sannigrahi A, Chowdhury S, Das B, Banerjee A, Halder A, Kumar A, Saleem M, Naganathan AN, Karmakar S, Chattopadhyay K (2021) The metal cofactor zinc and interacting membranes modulate SOD1 conformation-aggregation landscape in an in vitro ALS model. Elife 10: e61453

Sannigrahi A, Ghosh S, Pradhan S, Jana P, Jawed JJ, Majumdar S, Roy S, Karmakar S, Mukherjee B, Chattopadhyay K (2024) Leishmania protein KMP-11 modulates cholesterol transport and membrane fluidity to facilitate host cell invasion. EMBO reports 25: 5561–5598

Sannigrahi A, Nandi I, Chall S, Jawed JJ, Halder A, Majumdar S, Karmakar S, Chattopadhyay K (2019) Conformational switch driven membrane pore formation by Mycobacterium secretory protein MPT63 induces macrophage cell death. ACS chemical biology 14: 1601–1610

Sannigrahi A, Rai VH, Chalil MV, Chakraborty D, Meher SK, Roy R (2022) A Versatile Suspended Lipid Membrane System for Probing Membrane Remodeling and Disruption. Membranes 12: 1190

Sarangi NK, Basu JK (2018) Pathways for creation and annihilation of nanoscale biomembrane domains reveal alpha and beta-toxin nanopore formation processes. Physical Chemistry Chemical Physics 20: 29116–29130

Sathyanarayana P, Maurya S, Behera A, Ravichandran M, Visweswariah SS, Ayappa KG, Roy R (2018) Cholesterol promotes Cytolysin A activity by stabilizing the intermediates during pore formation. Proc Natl Acad Sci U S A 115: E7323–e7330

Scheek S, Brown MS, Goldstein JL (1997) Sphingomyelin depletion in cultured cells blocks proteolysis of sterol regulatory element binding proteins at site 1. Proceedings of the National Academy of Sciences 94: 11179–11183

Schwartz R, King J (2006) Frequencies of hydrophobic and hydrophilic runs and alternations in proteins of known structure. Protein Sci 15: 102–112

Seddon N, 1988. Biochemistry by Lubert Stryer. pp 1136. WH Freeman, New York. 1988.£ 39.95 or£ 24.95 (pbk) ISBN 0-7167-1843-X or-1920-7. Wiley Online Library.

Segrest JP, De Loof H, Dohlman JG, Brouillette CG, Anantharamaiah G (1990) Amphipathic helix motif: classes and properties. Proteins: Structure, Function, and Bioinformatics 8: 103–117

Siebold C, Byrne E, Miller P, Sansom M, Hedger G, Newstead S, Sircar R, Luchetti G, Nachtergaele S, Tully M (2016) Structural basis for Smoothened regulation by its extracellular domains. Nature 535

Simons K, Ehehalt R (2002) Cholesterol, lipid rafts, and disease. J Clin Invest 110: 597–603

Slotte JP, Bierman EL (1988) Depletion of plasma-membrane sphingomyelin rapidly alters the distribution of cholesterol between plasma membranes and intracellular cholesterol pools in cultured fibroblasts. Biochem J 250: 653–658

Smaby JM, Brockman HL, Brown RE (1994) Cholesterol’s interfacial interactions with sphingomyelins and-phosphatidylcholines: hydrocarbon chain structure determines the magnitude of condensation. Biochemistry 33: 9135–9142

Strandberg E, Killian JA (2003) Snorkeling of lysine side chains in transmembrane helices: how easy can it get? FEBS letters 544: 69–73

Vadia S, Arnett E, Haghighat A-C, Wilson-Kubalek EM, Tweten RK, Seveau S (2011) The pore-forming toxin listeriolysin O mediates a novel entry pathway of L. monocytogenes into human hepatocytes. PLoS pathogens 7: e1002356

Vanderroost J, Avalosse N, Mohammed D, Hoffmann D, Henriet P, Pierreux CE, Alsteens D, Tyteca D (2023) Cholesterol and Sphingomyelin Polarize at the Leading Edge of Migrating Myoblasts and Involve Their Clustering in Submicrometric Domains. Biomolecules 13: 319

Verkleij AJ, Zwaal RF, Roelofsen B, Comfurius P, Kastelijn D, van Deenen LL (1973) The asymmetric distribution of phospholipids in the human red cell membrane. A combined study using phospholipases and freeze-etch electron microscopy. Biochimica et Biophysica Acta (BBA)-Biomembranes 323: 178–193

von Heijne G (1994) Membrane proteins: from sequence to structure. Annual review of biophysics and biomolecular structure 23: 167–192

Weinberger A, Tsai F-C, Koenderink GH, Schmidt TF, Itri R, Meier W, Schmatko T, Schröder A, Marques C (2013) Gel-assisted formation of giant unilamellar vesicles. Biophysical journal 105: 154–164

Wu EL, Cheng X, Jo S, Rui H, Song KC, Dávila-Contreras EM, Qi Y, Lee J, Monje-Galvan V, Venable RM, 2014. CHARMM-GUI membrane builder toward realistic biological membrane simulations. Wiley Online Library.

Yamaji A, Sekizawa Y, Emoto K, Sakuraba H, Inoue K, Kobayashi H, Umeda M (1998) Lysenin, a novel sphingomyelin-specific binding protein. Journal of Biological Chemistry 273: 5300–5306

Yilmaz N, Yamaji-Hasegawa A, Hullin-Matsuda F, Kobayashi T, 2018. Molecular mechanisms of action of sphingomyelin-specific pore-forming toxin, lysenin, Seminars in cell & developmental biology. Elsevier, pp. 188–198.

Yuan C, Furlong J, Burgos P, Johnston LJ (2002) The size of lipid rafts: an atomic force microscopy study of ganglioside GM1 domains in sphingomyelin/DOPC/cholesterol membranes. Biophysical journal 82: 2526–2535

Zhang Y, Feller SE, Brooks BR, Pastor RW (1995) Computer simulation of liquid/liquid interfaces. I. Theory and application to octane/water. The Journal of chemical physics 103: 10252–10266

Zidovetzki R, Levitan I (2007) Use of cyclodextrins to manipulate plasma membrane cholesterol content: evidence, misconceptions and control strategies. Biochimica et Biophysica Acta (BBA)-Biomembranes 1768: 1311–1324

## References

Schütte OM, Mey I, Enderlein J, Savić F, Geil B, Janshoff A, Steinem C (2017) Size and mobility of lipid domains tuned by geometrical constraints. Proceedings of the National Academy of Sciences 114: E6064–E6071

